# Pangenome biology and evolution in harmful algal-bloom-forming pelagophyte algae

**DOI:** 10.1101/2024.10.30.620910

**Authors:** Shannon J. Sibbald, Maggie Lawton, Charlotte Maclean, Andrew J. Roger, John M. Archibald

## Abstract

In prokaryotes lateral gene transfer (LGT) is a key mechanism leading to intra-species variability in gene content and the phenomenon of pangenomes. In microbial eukaryotes, however, the extent to which LGT-driven pangenomes exist is unclear. Pelagophytes are ecologically important marine algae that include *Aureococcus anophagefferens* – a species notorious for causing harmful algal blooms. To investigate genome evolution across Pelagophyceae and within *Aureococcus anophagefferens*, we used long-read sequencing to produce high-quality genome assemblies for five strains of *Ac. anophagefferens* (52-54 megabase-pairs; Mbp), a telomere-to-telomere assembly for *Pelagomonas calceolata* (32 Mbp), and the first reference genome for *Aureoumbra lagunensis* (41 Mbp). Using comparative genomics and phylogenetics, we show remarkable strain level genetic variation in *Ac. anophagefferens* with a pangenome (23,356 orthogroups) that is 81.1% core and 18.9% accessory. Although gene content variation within *Ac. anophagefferens* does not appear to be largely driven by recent prokaryotic LGTs (2.6% of accessory orthogroups), 368 orthogroups were acquired from bacteria in a common ancestor of all analyzed strains and are not found in *P. calceolata* or *Au. lagunensis*. 1,077 recent LGTs from prokaryotes and viruses were identified within Pelagophyceae overall, constituting 3.5-4.0% of the orthogroups in each species. This includes genes likely contributing to the ecological success of pelagophytes globally and in long-lasting harmful blooms.

## INTRODUCTION

A large fraction of the diversity of life resides within microbial eukaryotes. However, much of this diversity is not represented in genomic databases; a lack of sequencing depth and breadth within and between lineages of eukaryotic microbes hinders our ability to answer fundamental questions related to genome biology, evolution and ecology^1^. Of particular significance is the concept of the pangenome, i.e., the collection of genes within a given species, some of which are found in all strains (the ‘core’ genome) and others only in certain strains (the ‘accessory’ genome). Analyses of vast genome sequence datasets covering the full breadth of bacterial and archaeal diversity have revealed that pangenomes are a common feature of prokaryotes, where LGT results in substantial coding capacity differences within members of a species and contributes to niche differentiation and key phenotypic differences such as antibiotic resistance in bacteria (e.g., see refs. ^2–4^ and references therein).

In the eukaryotic realm, pangenomes have been studied and identified primarily in fungal and plant species (e.g., see refs. ^5–7^) where the genomic datasets are sufficient to make such inferences. A prominent exception is in haptophyte algae such as *Gephyrocapsa huxleyi* (previously *Emiliania huxleyi*) where a substantial accessory genome was identified and predicted to contribute to metabolic and phenotypic differences between strains^8^. Wisecaver *et al.*^9^ recently showed that in the harmful algal bloom (HAB)-associated haptophyte *Prymnesium parvum*, extensive accessory genome content is linked to toxin metabolism. In the chrysophyte alga *Poteriospumella lacustris,* Majda *et al*.^10^ identified a sizeable pangenome with accessory genes involved in secondary metabolic functions.

Stramenopiles (heterokonts) are a diverse collection of eukaryotes that include heterotrophs as well as single-celled and multicellular algae and have relatively good genome sequence availability, particularly in oomycete plant pathogens and diatoms. Pelagophyte algae (Pelagophyceae) are a notable exception; until recently, pelagophytes were only represented by a single genome sequence^11–14^. The ecological significance of pelagophytes like the HAB-causing taxa *Aureococcus anophagefferens* and *Aureoumbra lagunensis*^15^, as well as *Pelagomonas calceolata*, which is among the most abundant eukaryotes in the open ocean^13^, make them a priority for genomic investigations. Furthermore, given the availability of genomic data in related stramenopiles, pelagophytes are an excellent lineage in which to explore the role of LGT in microbial adaptation. While the extent to which recent LGT contributes to eukaryotic genomes has been questioned (e.g., see ref. ^16^ for a skeptical viewpoint), it is increasingly considered to be an important mechanism underpinning eukaryote biology and evolution (e.g., see refs. ^17–22^).

Here we present highly contiguous, long-read sequencing-based genome assemblies for five strains of *Ac. Anophagefferens* (CCMP1707, 1708, 1850, 1984 and 3368), the first reference genome for *Au. lagunensis* CCMP1510, and a six-chromosome telomere-to-telomere assembly for *P. calceolata* CCMP1756. Orthogroup inference, functional assessment and phylogenetic analyses of these data reveal substantial differences in genome structure and coding capacity between these pelagophytes, differences that likely contribute to their ecological success in the world’s oceans. We also investigated the presence and functional implications of a pangenome within *Ac. anophagefferens,* showing evidence for an accessory genome comprising almost 20% of all orthogroups. Using a rigorous phylogenetic approach, we assessed the extent to which both the *Ac. anophagefferens* pangenome and gene content differences between pelagophyte species were driven by recent prokaryotic LGTs. Our results provide insight into the nature of pangenomes in microbial eukaryotes and show how recent LGTs from prokaryotes have impacted the biology of pelagophyte algae.

## METHODS

### Cell Culturing

Cultures of pelagophyte algae (*Aureococcus anophagefferens* CCMP1707, 1708, 1850, 1984 and 3368; *Aureoumbra lagunensis* CCMP1510; and *Pelagomonas calceolata* CCMP1756) were obtained from the National Center for Marine Algae and Microbiota (NCMA, East Boothbay, ME, USA). *Ac. anophagefferens* strains were maintained as either axenic (CCMP1984, 3368) or uni-eukaryotic cultures (CCMP1707, 1708, 1850) in L1-Si media prepared with artificial seawater^23^. *P. calceolata* CCMP1756 was grown axenically in L1-Si media prepared with artificial seawater, while *A. lagunensis* CCMP1510 was grown axenically in h/2 media made with artificial seawater^23^. All pelagophytes were grown with 100 µmol quanta m^−2^s^−1^ light on a 12hr : 12hr light : dark cycle at either 20°C (*Au. lagunensis* and *Ac. anophagefferens*) or 22-23°C (*P. calceolata*).

### RNA and DNA extraction and Sequencing

Total RNA was extracted from 50 mL of algal cell cultures in early exponential growth centrifuged at 5,020 × g at room temperature for 2 min. RNA from *Ac. anophagefferens* CCMP1984 was isolated using a modified TRIzol Reagent-based protocol (Thermo Fisher Scientific) in which cells were lysed using 2 mL of TRIzol Reagent and homogenized with a glass homogenizer for 10 min. RNA from other *Ac. anophagefferens* strains and pelagophyte species was extracted using a PureLink RNA Mini Kit (Thermo Fisher Scientific), with cells homogenized similarly. Isolated RNA was subsequently treated with a DNA digestion using TURBO DNase (Thermo Fisher Scientific). RNA quality was assessed on a formamide 1% agarose gel and quantity was estimated using a Qubit RNA Broad Range Assay (Thermo Fisher Scientific). Total RNA was sequenced at Génome Québec (Montreal, Canada) using stranded 100-bp paired-end mRNA libraries on Illumina’s NovaSeq 6000 (CCMP1510, 1707, 1708, 1756, 1984) or HiSeq platform (CCMP1850 and 3368).

To extract total genomic DNA, 50 mL (*Ac. anophagefferens*) or 100 mL (*P. calceolata* and *Au. lagunensis*) of cell culture in early exponential growth was isolated by centrifugation at 5,020 × g at 4°C for 5 min. DNA was extracted using either the DNeasy PowerSoil or PowerSoil Pro kit (Qiagen). DNA quality and quantity was assessed using a Qubit Broad Range DNA Assay (Thermo Fisher Scientific) and NanoDrop Spectrophotometer (Thermo Fisher Scientific). DNA integrity was assessed using 1% agarose gel electrophoresis. Further methods to additionally clean-up genomic DNA were performed, including CTAB based extractions^24^ for *Ac. anophagefferens* strains and *Au. lagunensis* or using a DNeasy PowerSoil clean-up kit (Qiagen) with *P. calceolata*. *P. calceolata* genomic DNA was additionally treated with a Short Read Eliminator (SRE) kit (Circulomics) to remove fragments below 25 kbp. Genomic DNA quality was further assessed using restriction enzyme digests of total genomic DNA with *Nco*I (Thermo Fisher Scientific) to confirm that DNA would be amenable to library preparation protocols.

High-quality short read data were generated using Illumina sequencing of PCR-free 150 bp paired-end libraries at Génome Québec (Montreal, Canada) on either the NovaSeq 6000 or HiSeq platforms. Long read sequencing data were generated in-house using Oxford Nanopore Technologies’ Minion platform. Sequencing libraries were prepared using a 1D Genomic DNA by Ligation SQK-LSK109 kit (CCMP1984) or 1D Native Barcoding Genomic DNA by Ligation SQK-LSK109 kit with the barcoding EXP-NBD103 expansion (all other strains) (Oxford Nanopore Technologies). In total, four flow-cells (FLO-MIN106) were used for long-read sequencing, with species/strain combination on individual flow cells as follows: (1) *Ac. anophagefferens* CCMP1984, (2) *Ac. anophagefferens* CCMP1850 and CCMP3358, (3) *Ac. anophagefferens* CCMP1707 and CCMP1708, and (4) *P. calceolata* CCMP1756 and *Au. lagunensis* CCMP1510. In all cases flow-cells were run for 48 hrs.

### Sequence Processing and Genome Assembly

The quality of Illumina-based DNA and RNA sequencing short-reads was assessed using FastQC (v0.11.9) (https://www.bioinformatics.babraham.ac.uk/projects/fastqc/), which informed the trimming of adaptors and low-quality regions using Trimmomatic (v0.39)^25^. Long-read sequencing data from barcoded samples were demultiplexed using Deepbinner (v0.2.0) (https://github.com/rrwick/Deepbinner) and Guppy (v.3.3.0) (Oxford Nanopore Technologies). Long-reads were basecalled using Guppy (v.3.3.0) (Oxford Nanopore Technologies) and adaptors were trimmed using Porechop (v0.2.1) (https://github.com/rrwick/Porechop). Long-reads were filtered using Filtlong (v0.2.1) (https://github.com/rrwick/Filtlong) to remove reads with a Phred quality score lower than Q7 for all read-sets. The quality and statistics for each long-read dataset was then assessed using Nanoplot (v1.13)^26^. Depending on the depth of sequencing for a given sample, additional filtering of reads shorter than 5-20 kbp and those with lower quality was carried out to retain only a subset of sequencing depth for genome assembly.

Long-read based genome assemblies were trialed for each strain and species using a variety of assemblers (Flye v2.6^27^, Canu v2.2^28^ and Raven v0.0.1^29^), read-sets and assembly parameters. Resulting assemblies were assessed using QUAST (v5.0.2)^30^, BUSCO (v3.0, database eukaryota.obd9)^31^ and Assembly Likelihood Estimation (ALE)^32^. Genome assemblies selected for further curation were generated using Flye (v2.6)^27^ with minimum overlap parameters set to either 6.5 kbp (*Ac. anophagefferens* CCMP1984 and 1707) or 10 kbp.

Genome assemblies were manually curated to inspect for mis-assemblies, remove contamination and reduce duplicated regions that failed to collapse into a single haplotype in diploid species. To search for mis-assemblies, long reads were mapped onto the genome using ngmlr (v0.2.7)^33^ and visualized using Tablet (v1.21.02.08)^34^. In the case of the *Ac. anophagefferens* reference strain (CCMP1984), long read mapping was also used in combination with assembly visualization using Bandage^35^ to scaffold initially assembled contigs. Contigs that were deemed contamination were identified and removed based on very low or high sequencing coverage (<5X or >200X), deviation from the genomic average GC content, and blastN searches against the nt database^36^ along suspect contig lengths. Candidate regions that failed to collapse into a single haplotype due to high heterozygosity were identified using Purge Haplotigs (v1.1.2)^37^ with the minimum alignment coverage set to 80%; any contigs flagged by Purge Haplotigs as redundant were manually inspected prior to removal from haploid genome assemblies.

Manually curated long-read assemblies were corrected at the single nucleotide level using long-reads with Nanopolish (v0.9.0) (https://github.com/jts/nanopolish) and high-accuracy Illumina reads with multiple iterations of Pilon (v1.2.3.)^38^ until the number of identified base-pair changes in each iteration stabilized. After sequence polishing, organelle genomes (mitochondrion and plastid) were identified using expected GC content, length and confirmed with blastN homology searches against the nt database^36^; organelle genomes were removed from the final genome assemblies and considered separately.

### Ploidy, GC content, telomere, repeat, rDNA and genome synteny assessment

Ploidy was assessed using nQuire^39^ and ploidyNGS^40^ with Illumina short reads mapped to the genome assembly via Bowtie 2 (v2.3.1)^41^. To assess base composition heterogeneity within a genome, GC content was calculated across each contig using a sliding window approach in 10 kbp windows with 1 kbp increments and plotted using a custom script in R. Telomeres were identified by scanning contig ends for tandem repeats involving permutations of the TTAGGG telomeric repeat. Repeats in each genome were identified and soft-masked using RepeatMasker (v4.1.3) (http://www.repeatmasker.org).

Ribosomal DNA (rDNA) regions were identified using a combination of blastN^36^ with known rDNA genes from pelagophytes as queries, RNAmmer-1.2^42^, and Rfam (v14)^43^ searches followed by manual curation. rDNA regions within each genome were aligned with Sibelia (v 3.0.7)^44^ and visualized with Circos^45^. Whole genome alignments were generated using MUMmer (v4.0)^46^ with nucmer based alignments requiring 90% identity, a 2,500 bp minimum alignment length, and an increased minimum exact match length of 50 bp. Resulting whole genome alignments between *Ac. anophagefferens* strains were assessed using dnadiff and plotted with mummerplot in MUMmer (v4.0)^46^.

### Gene Prediction and Orthogroup Inference

Transcriptomes were assembled using Trinity^47^ in genome-guided mode, where HISAT2 (v2.2.1)^48^ was used to map RNA-seq reads to the genome. Gene models for protein-coding genes were predicted by combining BRAKER2 (v2.1.5)^49^ and PASA (v.2.3.3) gene model predictions into consensus gene structures using EVidenceModeler (v.2.1.0)^50^ with the *ab initio* based BRAKER2 predictions and transcriptome based PASA predictions weighted 1:10. Protein-coding genes were also predicted separately using Funannotate (v1.8.15) (https://github.com/nextgenusfs/funannotate/) with GeneMark-ET (v.4.72)^51^. Genes predicted to encode proteins <90 amino acids were removed from all gene sets.

The completeness of predicted protein-coding gene sets was assessed using BUSCO (v3.0, database eukaryota.obd9)^31^ and the similarity of gene models predicted by EVidenceModeler and Funannotate was assessed using ParsEval (v0.16.0)^52^. BUSCO completeness of gene prediction sets combined with visual assessment of gene-model structure using RNA-seq reads aligned to the genome was used to select the best gene prediction dataset. Using the ParsEval output, gene loci determined to be unique to Funannotate predicted gene sets were added to those from EVidenceModeler to create more complete protein-coding gene sets for all pelagophytes; the exception was for *P. calceolata* CCMP1756 where unique EVidenceModeler gene loci were added to the Funannotate gene sets. Many gene models were manually curated to fix incorrectly predicted intron splice sites and gene boundaries where RNA-seq data supported these changes. Various intron characteristics of predicted protein-coding genes were determined using a custom python script. To assess the similarity of *Ac. anophagefferens* strains and pelagophyte species at the protein-coding gene level, the percent identity of all best-reciprocal blast hits using blastP for all pairwise comparisons of strain/species protein-coding gene sets were determined. An e-value cut-off of 0 was used for identifying intra-species best-reciprocal blast hits, while 1e-30 was used for inter-species comparisons.

Orthologous groups shared between pelagophytes were inferred using both Broccoli (v1.1)^53^ and OrthoFinder (v2.5.5)^54^ using predicted protein-coding gene sets in two diatom species (*Thalassiosira pseudonana* and *Phaeodactylum tricornutum*) as an outgroup. Genes unassigned to orthogroups in *Ac. anophagefferens* strains (singletons) were re-assessed using blastx and tblastx searches^36^ against custom databases of predicted protein-coding gene sets and genomes in other strains, respectively; singletons with hits >85% identity and >70% gene coverage threshold were added to the ‘best-hits’ orthogroup or to a new orthogroup. OrthoFinder orthogroups were selected for further refinement and analysis because a higher proportion of genes were assigned to orthogroups, and pangenome estimates within *Ac. anophagefferens* were more conservative. Using a custom python script, genes determined to be core pangenome genes using Broccoli but not OrthoFinder were identified; the orthogroup assignment of these genes were re-assessed using sequence similarity and gene neighborhood synteny between the various strains.

Single-copy orthogroups with an ortholog present in all pelagophyte and diatom outgroup species were used to infer phylogenetic relationships between the *Ac. anophagefferens* strains and pelagophyte species. Each orthogroup was aligned using MAFFT-linsi (v7.474)^55^ and trimmed with BMGE (v1.1)^56^ prior to concatenation into a multi-gene alignment. A maximum-likelihood (ML) tree was inferred using IQTREE (v2.3.0)^57^ under the model Q.pfam+F+I+R4 (as selected using the MFP model test using BIC criteria^58^) with 1,000 ultra-fast bootstrap (UFBoot2) replicates^59^. Single gene trees for each of the single copy orthogroup alignments were inferred in IQTREE under the model LG4X and 1,000 UFBoot2 replicates. Single gene trees were used to assess the gene concordance factor (gCF)^57^ for individual gene trees to the previously inferred multi-gene tree in IQTREE. Alternative topologies with differing *Ac. anophagefferens* strain relationships were assessed using likelihood and AU tests^60^ in IQTREE.

### Functional Annotation/Enrichment

Predicted protein-coding genes were functionally annotated using interproscan-5.47-82^61^ with default settings and eggNOG-mapper (v2.1.4-2)^62^. Gene ontology (GO) terms identified by interproscan were used to determine significantly enriched biological processes and molecular functions between protein-coding gene sets of species and strains using GOEnrichment (https://github.com/DanFaria/GOEnrichment) with a p-value cut off of <0.05. GO terms associated with species or strain-specific genes/orthogroups, or candidate LGTs were visualized using REVIGO (v1.8.1)^63^. Signal peptides were predicted using SignalP (v6)^64^.

### Phylogenetic Pipeline for LGT identification

All predicted protein-coding genes were queried against both the NCBI nr and MMETSP transcriptome^65^) databases using DIAMOND with the ‘more-sensitive’ option^66^; the top 1,000 hits with an e-value < 1e-10 were retrieved for each gene. Genes with >10 hits, where at least three of the predicted homologs were bacteria, archaea and/or viruses were considered further. Within this subset of protein-coding genes, DIAMOND hits were compared for each gene within an orthogroup and combined non-redundantly if >50% of hits were identical. Where <50% hit overlap was found within genes of an orthogroup, the orthogroup and gene models were inspected and manually refined prior to tree building.

Homologs were aligned using MAFFT (v7.474)^55^ with default settings. Divergent sites in the alignment were trimmed with BMGE (v1.1)^56^ using the BLOSUM30 scoring matrix and an entropy parameter of 1; all sites and sequences with >50% gap rates were removed. Taxonomic redundancy of individual alignments was reduced at the genus (eukaryotes, except for stramenopiles for whom all OTUs were retained) or phyla level (bacteria/archaea) using custom scripts based on the TreeTuner pipeline^67^ and utilizing FastTree (v2.1.11)^68^. Remaining OTUs were realigned and trimmed as before, and the final alignment was used to infer a ML phylogeny using IQTREE (v2.3.0)^57^ under the LG4X model with 1,000 UFBoot2 replicates^59^.

To assess phylogenies for recent LGTs from prokaryotes or viruses into pelagophytes, tree topologies were evaluated based on the position of pelagophyte sequences in relation to other eukaryote and prokaryote OTUs using similar criteria as in Eme *et al.*^20^. The topology of all single-gene trees were manually examined and candidate LGTs were considered based on the following criteria: (i) candidate LGTs in pelagophytes could not branch sister to homologs in any other non-pelagophyte stramenopile group, (ii) other stramenopile sequences had to be separated from any prokaryote-pelagophyte containing clade by at least three bipartitions with >80% UFBoot2 support, and (iii) any eukaryote sequences intervening between pelagophyte and prokaryote OTUs could not include more than two major eukaryotic groups (e.g., haptophytes or alveolates). Additionally, the presence of more than two major eukaryotic groups neighboring the prokaryote-pelagophyte containing clade, and the proportion of prokaryotic to eukaryotic sequences within the prokaryote-pelagophyte containing clade, as well as within three and five well-supported bipartitions (>80% UFBoot2 support) of the clade was considered. Genomic context of candidate LGTs was assessed for neighboring eukaryotic genes and long-read sequencing support for these regions of genome assemblies. Intron presence and evidence of gene expression was also evaluated.

## RESULTS

### Pelagophyte genomes are structurally diverse and genetically heterogeneous

The genomes of seven pelagophyte strains/species were sequenced and assembled primarily using long-read sequence data (*Ac. Anophagefferens* CCMP1707, 1708, 1850, 1984 and 3368; *Pelagomonas calceolata* CCMP1756; and *Aureoumbra lagunensis* CCMP1510; Table 1). All assemblies are highly contiguous (L50 = 3-29 contigs, N50 = 0.53-5.4 Mbp) and have similar BUSCO-based levels of completeness (83.5-86.1% complete against the eukaryota.obd9 dataset; Extended Data Figure 1). The average GC content of these genomes varies dramatically, between 39.4% in *Au. lagunensis* and 62.9%-69.9% in *P. calceolata* and *Ac. anophagefferens*, respectively. The genomes of all *Ac. anophagefferens* strains are predicted to be diploid, while *P. calceolata* and *Au. lagunensis* appear to be haploid (Extended Data Figure 2).

**Table 1.** Summary of genome assembly statistics for seven pelagophyte genomes. Genome statistics shown are based on haploid versions of long-read sequencing-based genome assemblies. BUSCO completeness is based on comparison of predicted protein-coding genes to the eukaryotic dataset (v3; n=303), where BUSCO presence includes single-copy, duplicated, and fragmented BUSCO predicted genes (see also Extended Data Figure 1).

|  | *Ac. anophagefferens* |  |  |  |  | *P. calceolata* | *Au. lagunensis* |
| --- | --- | --- | --- | --- | --- | --- | --- |
|  | CCMP<br>1984 | CCMP<br>1850 | CCMP<br>3368 | CCMP<br>1707 | CCMP<br>1708 | CCMP<br>1756 | CCMP<br>1510 |
| Contigs | 143 | 209 | 190 | 233 | 233 | 6 | 32 |
| Genome<br>size<br>(Mbp) | 52.76 | 53.85 | 52.17 | 52.97 | 52.86 | 32.30 | 41.32 |
| N50<br>(Mbp) | 0.983 | 0.620 | 0.742 | 0.574 | 0.529 | 5.422 | 3.465 |
| L50 | 10 | 22 | 17 | 27 | 28 | 3 | 5 |
| Coverage | 46× | 79× | 94× | 58× | 90× | 119× | 121× |
| GC% | 69.89 | 69.49 | 69.95 | 69.83 | 69.81 | 62.93 | 39.46 |
| Ploidy | diploid | diploid | diploid | diploid | diploid | haploid | haploid |
| Repeats | 8.1 Mbp | 8.2 Mbp | 5.7 Mbp | 6.2 Mbp | 5.9 Mbp | 1.9 Mbp | 9.1 Mbp |
| Protein<br>Coding<br>Genes | 25,737 | 26,943 | 25,956 | 24,125 | 24,557 | 17,704 | 17,841 |
| BUSCO<br>present | 92.7% | 92.1% | 91.5% | 91.8% | 92.4% | 91.1% | 88.9% |

Haploid genome assemblies of the five *Ac. anophagefferens* strains range from 52.17-53.85 Mbp across 143-233 contigs (Table 1). This is comparable to, but much more contiguous than, the original CCMP1984 reference genome^11^ (11-16 contigs have telomeres on one or both ends), and generally more contiguous and less artifactually duplicated than recently published *Ac. anophagefferens* assemblies for CCMP1984, 1850 and 1707^12^. The genome of *P. calceolata* CCMP1756 (32.30 Mbp) is fully resolved, consisting of six telomere-to-telomere chromosomal sequences, each 4-6 Mbp in length – consistent with the genome structure of *P. calceolata* RCC100^13^. Putative centromere regions of ∼200-300 kbp were identified in all six *P. calceolata* chromosomes that have a lower-than-genome-average GC content (∼51%) and low levels of sequence conservation (Extended Data Figure 3). The genome of *Au. lagunensis* (41.32 Mbp) assembled in 32 contigs, with five contigs having telomeres on one or both ends.

The amount of repeat content identified in *P. calceolata* was much lower than the other pelagophyte genomes, consisting of 1.9 Mbp (∼6% of its genome) of mostly unclassified repeat elements (Extended Data Figure 4, Supplementary Information Table 1). In contrast, ∼22% of the *Au. lagunensis* genome is repetitive (9.11 Mbp), including an expansion of LTR retrotransposable elements comprising 6.2% of its genome. Overall, *Au. lagunensis* has ∼800 more annotated LTRs than any other pelagophyte, including a 2-4× increase in LTR Copia and Gypsy elements. Annotated repeat content across *Ac. anophagefferens* genomes varied from 5.7-8.2 Mbp (11.2-15.4% of the genome). Some strain specific differences were observed, including an increase in LINEs in CCMP1850 (213 vs. 0-63 in other strains) and at least 300 additional LTRs in CCMP1850, suggesting that this strain has undergone recent repeat expansion.

The number of predicted protein-coding genes in the pelagophyte genomes sequenced herein ranges from ∼17,800 in *P. calceolata* and *Au. lagunensis* to ∼24,000-27,000 in the *Ac. anophagefferens* strains. Despite these differences, inferred levels of BUSCO completeness across all genomes were similar (Table 1; Extended Data Figure 1); 22 BUSCO genes were not identified in any strain/species, suggesting that they are absent or highly divergent in pelagophytes. *Ac. anophagefferens* strains have 3.5-9× more introns in their protein-coding genes than the other pelagophytes (0.433-0.917 introns per gene compared to 0.186 in *P. calceolata* and 0.135 in *Au. lagunensis*; Supplementary Information Table 2)). Intron sizes were similar in all three species (103-177 bp average). Notably, a large fraction of introns in *Ac. anophagefferens* (∼14%-17%) and *P. calceolata* (∼14%) were predicted to use an alternative GC-AG splice signal; this is much higher than in other eukaryotes (e.g., 0.3-2% in plants and metazoans^69^ and 4% in the red alga *Cyanidioschyzon merolae*^70^), and is likely attributable to their high genome GC content. In contrast, in *Au. lagunensis*, where the average genome GC content is much lower, an alternative GC-AG splice signal was predicted in 0.32% of introns.

### Aureococcus anophagefferens has a pangenome

To establish a foundation for a fine-scale, intra-species comparative genomic analysis, we inferred the degree of relatedness between sequenced *Ac. anophagefferens* strains via rDNA, organelle and whole genome similarity, as well as the overall identity between orthologous protein-coding genes. 18S rDNA copies in *Ac. anophagefferens* were found to be 98.9-100% identical within and between strains (including intron-containing and intron-less versions); intra- and inter-strain similarities are similar (see Supplementary Information). The plastid genomes of the strains are 99.9% identical across their entire length, while mitochondrial genomes are 99.4-100% identical^71^. Furthermore, complete *Ac. anophagefferens* nuclear genomes align to one another at >97% average nucleotide identity across 89-97% of their lengths (Extended Data Figure 5A-D), and pairwise ortholog similarity between all protein-coding genes in *Ac. anophagefferens* strains showed an average of ∼97% amino acid identity (Extended Data Figure 5E). By contrast, average pairwise ortholog similarity is 58.5% between *Ac. anophagefferens* and *P. calceolata*, and only 48.5% between *Ac. anophagefferens* and *Au. lagunensis.* Altogether, these data strongly suggest that the five strains of *Ac. anophagefferens* are closely related members of the same species.

Specific relationships between *Ac. anophagefferens* strains were inferred using a multi-gene tree approach, including 1,328 single-copy genes found in all five strains and the other pelagophyte species. The resulting relationships between strains were maximally supported (Figure 1 schematic); the gene concordance factor (gCF) of individual gene trees to the multi-gene tree, however, was only 17.4%, suggesting conflicting and/or limited signal in the individual single gene trees. An AU-test of a number of alternative strain relationships determined that they were all significantly worse than the maximum likelihood topology at a confidence interval of >99.9%.

**Figure 1.**
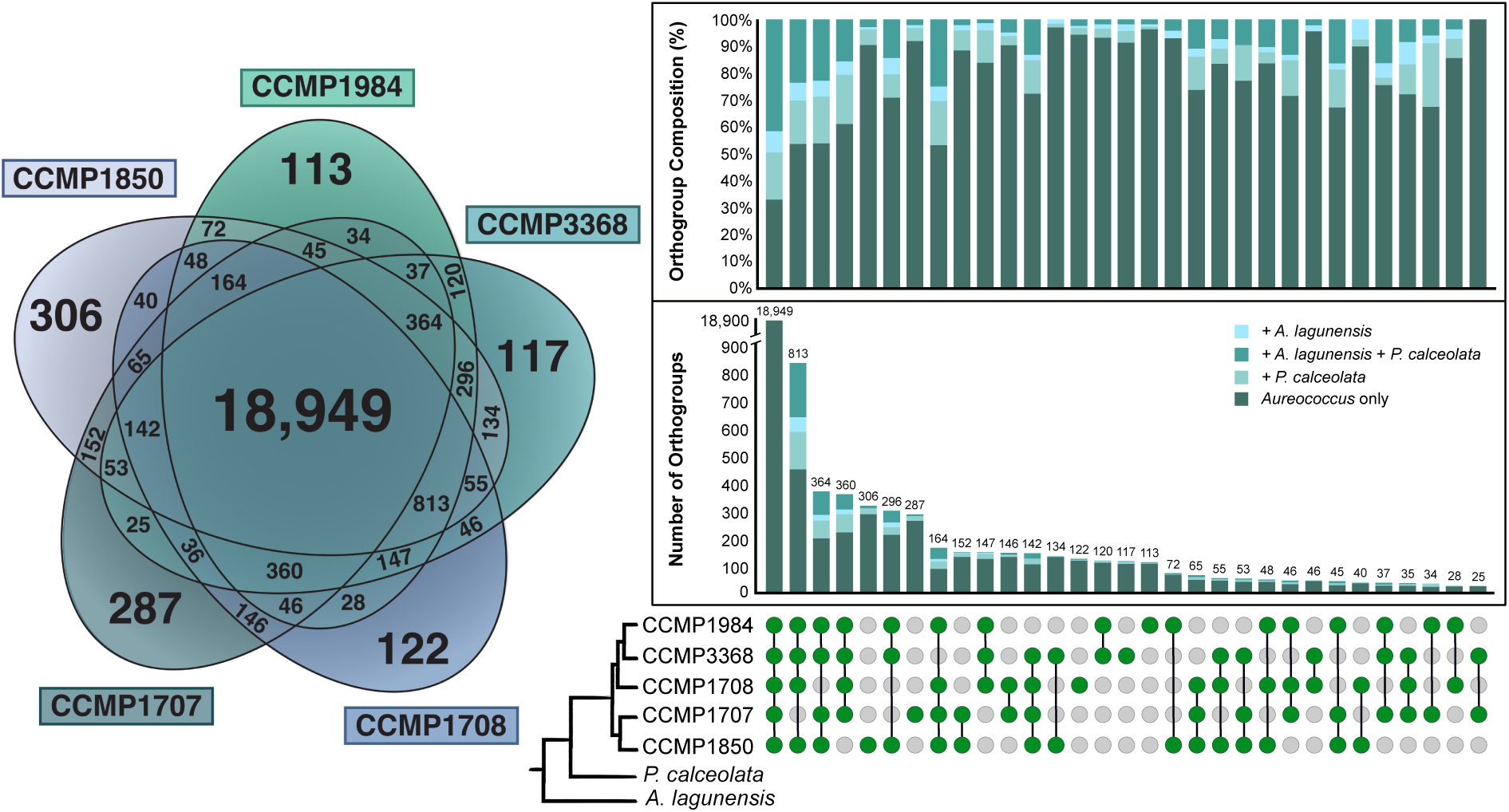
An extensive pangenome in *Aureococcus anophagefferens.* The total pangenome is estimated at 23,356 orthogroups (including singletons). The Venn-diagram (left) shows the number of orthogroups shared among all five analyzed strains of *Ac. anophagefferens* (core genome; 81.1% of orthogroups), as well as those found in select strains or only one strain (accessory genome; 18.8% of orthogroups). The column graph (right) shows the number of orthogroups/singletons present in each pangenome sub-category ordered from largest to smallest. Pangenome sub-categories are mapped onto a schematic phylogeny representative of *Ac. anophagefferens* strain relationships inferred using 1,328 single-copy core orthogroups found across all pelagophyte genomes (green = present in strain, grey = absent in strain). The number and proportion of orthogroups/singletons with orthologs in *P. calceolata* CCMP1756 and/or *Au. lagunensis* CCMP1510 for each pangenome sub-category is shown within a given column and above the column graph, respectively.

Using the protein-coding gene sets for five strains, a pangenome of *Ac. anophagefferens* was estimated to contain 23,356 orthogroups and singletons (Figure 1). 18,949 orthogroups (81.1%) are part of the core genome, i.e., common to all strains. The remaining 4,407 orthogroups/singletons (18.9%) were identified in one to four strains of *Ac. anophagefferens*, comprising the accessory genome. Nearly half of the accessory genome orthogroups are present in four of five strains (∼42%) and/or have homologs in either *P. calceolata* or *Au. lagunensis* (∼26%) (Figure 2), suggesting that a large proportion of the accessory genome may be due to differential gene loss. Nonetheless, the remaining half of the accessory genome was identified in a smaller subset of *Ac. anophagefferens* strains (between one and three), including 4% of orthogroups/singletons that were strain specific. This suggests that other mechanisms such as recent LGT, gene duplication, and *de novo* gene origin played a role in intra-species accessory genome formation (see below).

**Figure 2.**
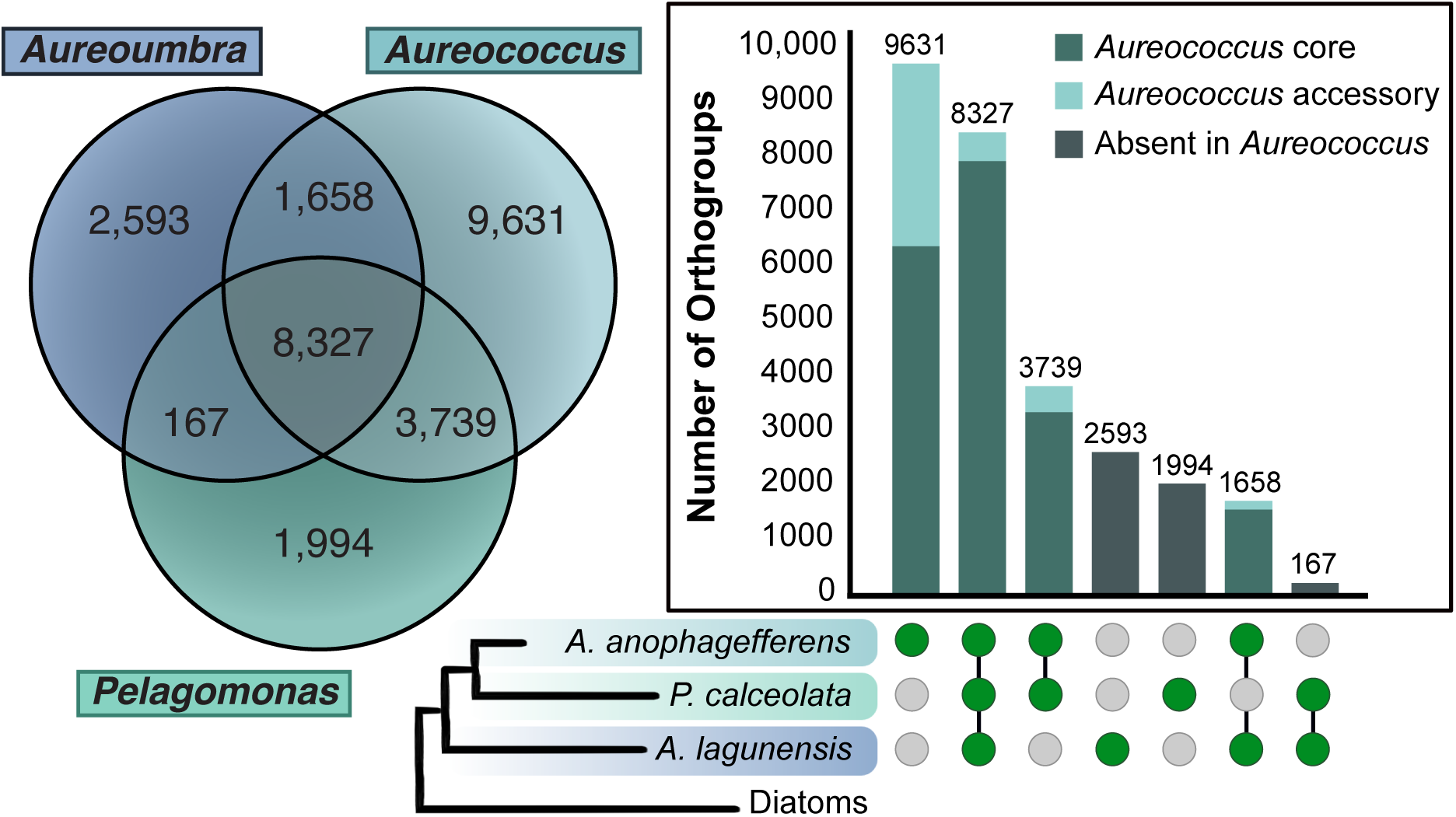
Genes shared amongst pelagophyte algae. The Venn-diagram (left) indicates the number of orthogroups/singletons shared between or unique to *Aureococcus anophagefferens* (pangenome), *Pelagomonas calceolata* CCMP1756, and *Aureoumbra lagunensis* CCMP1510. The column graph (right) shows the number of orthogroups/singletons shared or unique to combinations of the pelagophyte algae. Pelagophyte shared gene categories are mapped onto a schematic phylogeny representative of known species relationships (green = present in species, grey = absent in species). When *Ac. anophagefferens* is present, the number of orthogroups that are part of the core or accessory genome in *Ac. anophagefferens* is shown.

Approximately two-thirds of the orthogroups in the *Ac. anophagefferens* pangenome could be assigned to an orthologous group in the EggNOG database (Figure 3A). While >70% of accessory orthogroups were found to have at least low levels of gene expression (>1 TPM), only 48.4% were assigned to an EggNOG orthogroup, with ∼1/3 of these having unknown functions (KOG/COG category ‘S’). While the strain-specific subset of accessory genes showed similarly robust evidence for expression (∼2/3 of such genes), they were even more divergent; only ∼1/3 were assigned to an EggNOG orthogroup and/or interpro annotation, and only ∼15% had >1 homolog in a non-pelagophyte species in the ‘nr’ database. Altogether, this indicates that a large fraction of accessory orthogroups, especially the strain-specific genes, are highly divergent from known sequences or are entirely novel.

**Figure 3.**
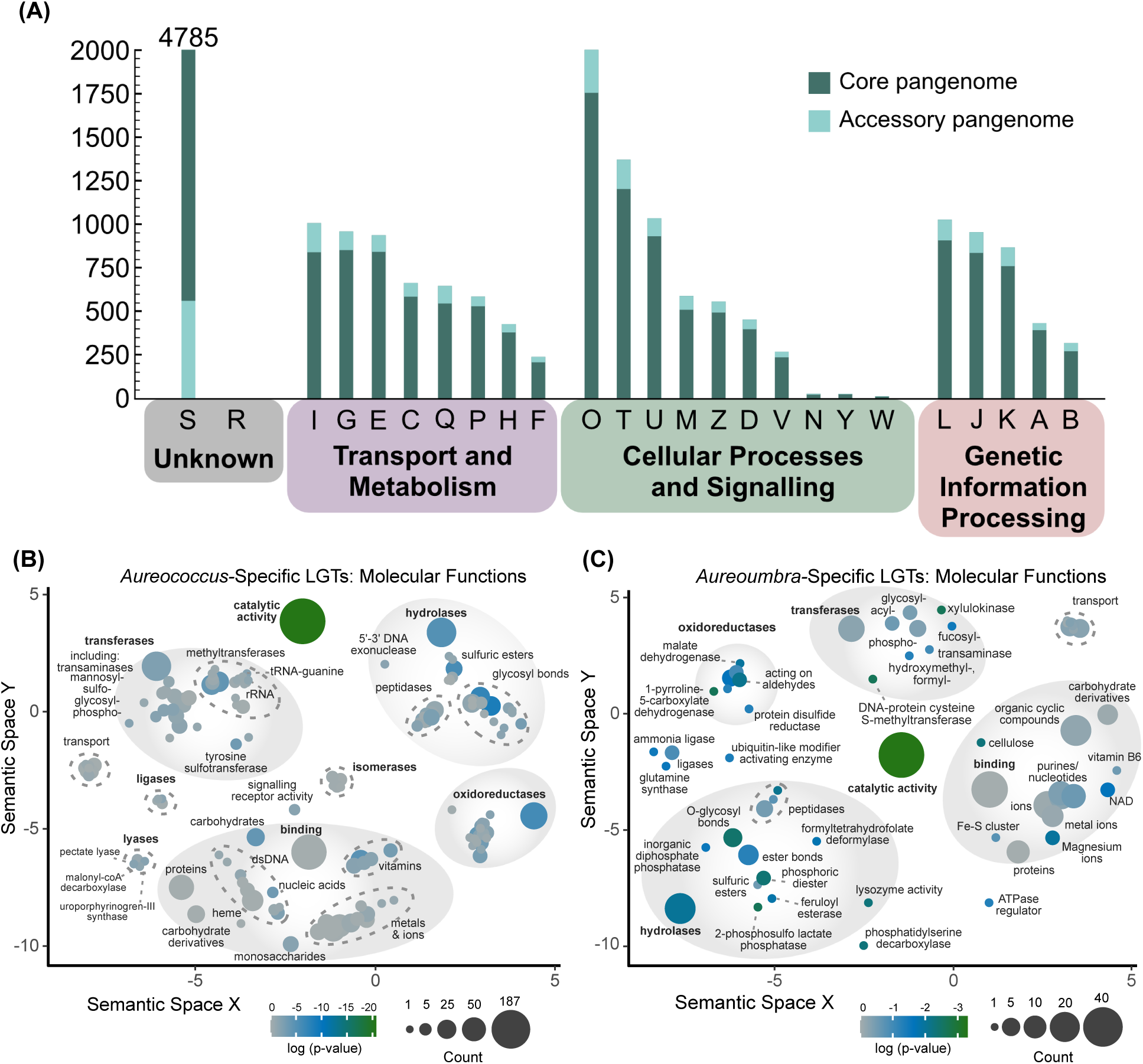
Functional categories of genes in the (A) *Ac. anophagefferens* pangenome, and in (B) *Ac. anophagefferens-*specific or (C) *Au. lagunensis*-specific LGTs. **(A)** The number of orthogroups/singletons assigned to each KOG/COG category within the *Ac. anophagefferens* pangenome (core genome = dark, accessory genome = light). When >2,000 orthogroups were present within a category, their number is indicated above a column. **(B/C)** REVIGO based scatter plots of GO terms associated with molecular functions of *Ac. anophagefferens-*specific **(B)** and *Au. lagunensis*-specific **(C)** recent LGTs. The number of orthogroups with a GO term and its degree of significant enrichment (log p-value) is reflected by the bubble size and colour, respectively. Labeling has been summarized for clarity; complete terms are available in Supplementary Table 3 and 4.

*Ac. anophagefferens* accessory orthogroups with EggNOG orthology were found to be associated with a variety of KOG/COG functional categories (Figures 3A). The most abundant functions are broadly related to either transport and metabolism (32.3%, including: lipids (‘I’; 8.7%), carbohydrates (‘G’; 5.7%), secondary metabolites (‘Q’; 5.4%) and amino acids (‘E’; 5.2%)), post-translational modifications (‘O’; 13.3%), or signal transduction mechanisms (‘T’; 8.9%). Overall, most of the set of accessory genes are associated with gene ontology (GO) terms related to metabolic processes (305, 70%), transmembrane transport (61, 31.6%) and/or biological regulation (36, 8%) (Extended Data Figure 6, Supplementary Information Tables 5 and 6). GO term enrichment analysis determined that compared to the full pangenome, the accessory gene-set is most significantly enriched (p<0.05) in biological processes related to fatty acid metabolism (13 orthogroups), macromolecule (41) and general biosynthetic processes (66), and translation (31). Some of the most numerous metabolic process related terms that were not enriched include those involved in protein modifications (63) (e.g., phosphorylation (32), ubiquitination (12), and glycosylation (4)), as well as the metabolism of organonitrogen-containing and nucleobase-containing compounds (152 and 93 orthogroups, respectively). Transport related orthogroups in the accessory genome involved many for monoatomic and metal ions (24), nitrogen-containing compounds (12), proteins (7) and sulfates (1).

Accessory orthogroups associated with molecular functions were largely related to catalytic activities (500, 46.5%) and binding (686, 63.8%); others were associated with transmembrane transport (52, 4.8%) or transcription regulator activity (20, 1.9%). Catalytic activity was found to encompass a variety of transferases (202), hydrolases (137), oxidoreductases (93), isomerases (25), lyases (11) and ligases (11) (see Supplementary Information). Approximately one-third of orthogroups associated with binding functions were predicted to bind ions (229) including metal ions (115), while another ∼1/3 were predicted to bind nucleic acid/nucleosides (223) and ∼1/6 carbohydrate derivatives (106). The most significantly enriched molecular functions within the accessory genome are phosphopantetheine (19) and vitamin binding (45), acyltransferase (69), carbon-sulfur lyase (5), and peroxidase (7) activity (Extended Data Figure 6, Supplementary Information Table 6).

Using a phylogenetic pipeline, we identified 113 recent LGTs of prokaryote/viral ancestry in the accessory genome of *Ac. anophagefferens,* 2.6% overall (Figure 4, Supplementary Information File 1). While this was not a sizeable portion of the accessory genome, several noteworthy functional terms were associated with these transferred genes, suggesting that they underpin metabolic differences between *Ac. anophagefferens* strains (Extended Data Figure 7, Supplementary Information Tables 7-8, Supplementary Information). This includes lipid metabolism (e.g., ceramide metabolism), xylan catabolism (e.g., an alpha-glucuronidase and a pectate lyase), aldehyde lyase activity, ADP catabolism, *de novo* IMP biosynthesis, an arsenate transmembrane transporter involved in arsenic detoxification, and DNA repair in stress response. A small number of strain-specific orthogroups/singletons (17/945) were predicted to be recent LGTs from prokaryotes/viruses, including a D5 family helicase/primase in CCMP1850 (Supplementary Information). A striking example is a ∼15 kbp low-GC region of *Ac. anophagefferens* virus (AaV) origin found uniquely in the CCMP1850 genome. This region contains a series of ORFs with predicted DUF285 domains – which have been observed to be in flux between the host and virus^72^ (Extended Data Figure 8, Supplementary Information).

**Figure 4.**
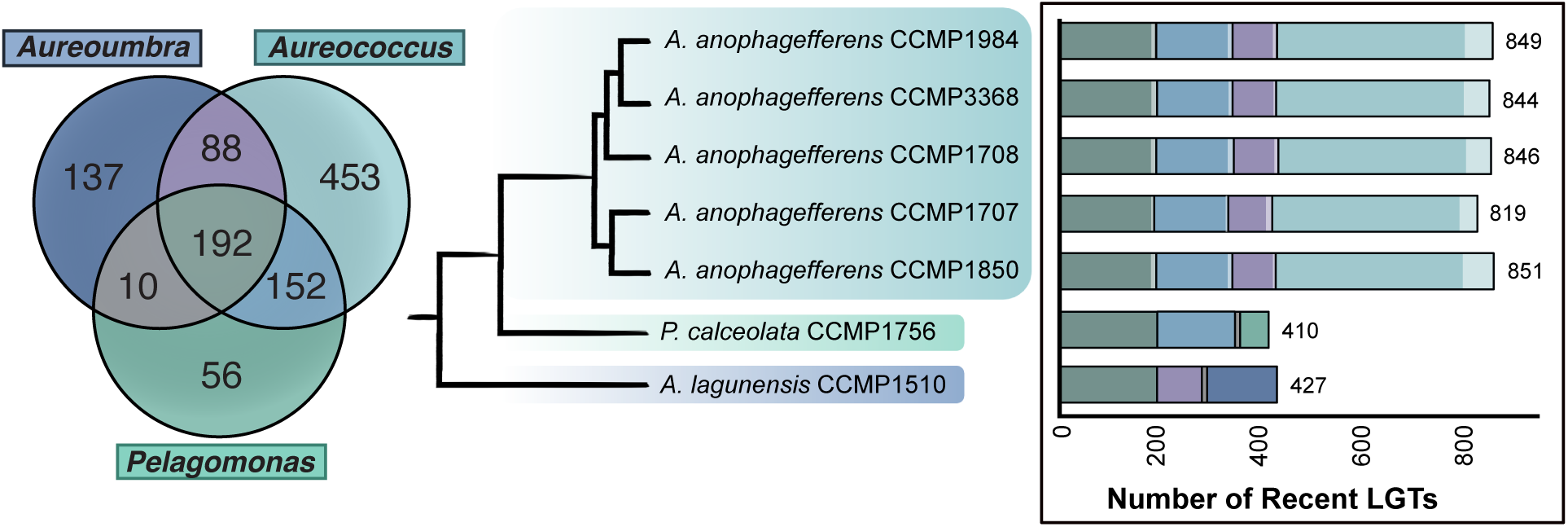
Recent lateral gene transfers (LGTs) in pelagophyte algae of prokaryotic/viral origin. The Venn-diagram (left) indicates the number of orthogroups/singletons inferred to be ‘recent’ LGTs from prokaryotes/viruses in various combinations of pelagophyte species. The column graph (right) shows the number of ‘recent’ LGTs in each species and specific strains within the *Ac. anophagefferens* pangenome. Within each column, the colours correspond to the same gene-sharing category as in the Venn-diagram, while the shade of the colour indicates whether it is part of the core (dark) or accessory (light) genome in each *Ac. anophagefferens* strain. The schematic phylogeny shown is as in Figure 1.

### *Ac. anophagefferens* has many ecologically relevant orthogroups absent in other pelagophytes

29.6% of identified orthogroups were found to be shared between all three pelagophyte species (8,327 of 28,108 orthogroups in total) (Figure 2). Unsurprisingly, more unique orthogroups are shared between the more closely related *Ac. anophagefferens* and *P. calceolata* (3,739) than between these algae and *Au. lagunensis*. While *P. calceolata* and *Au. lagunensis* only share 167 orthogroups to the exclusion of *Ac. anophagefferens*, *Ac. anophagefferens* and *Au. lagunensis* share 1,658 (∼10× more) where *P. calceolata* was absent, suggesting a common retention of genes potentially related to bloom formation and persistence in these HAB causing species. While all species had many species-specific orthogroups, the impact was particularly pronounced in *Ac. anophagefferens*, where over 40% of *Ac. anophagefferens-*containing orthogroups were specific to the species, ∼1/3 of which were in the accessory genome (see Supplementary Information).

73% of orthogroups specific to *Ac. anophagefferens* show at least low levels of gene expression (TPM >1); this number is 79% and 90% for *P. calceolata*-specific and *Au. lagunensis*-specific orthogroups, respectively. Despite evidence for active transcription, only ∼60% of *Ac. anophagefferens*-specific orthogroups have EggNOG orthogroups or interpro based annotations assigned to them (30% in *P. calceolata* and 45% in *Au. lagunensis*). Ignoring species-specific orthogroups aligned with EggNOG orthogroups of unknown function (∼25% in each species), the most abundant KOG/COG terms in *Ac. anophagefferens-*specific orthogroups were found to be related to post translational modifications (‘O’; 575, 10.6%), signal transduction (‘T’; 421, 8%), or transport/metabolism of lipids (‘I’; 394, 7.3%), carbohydrates (‘G’; 304, 5.6%) and amino acids (‘E’; 272, 5.1%). Similarly, in *P. calceolata* and *Au. lagunensis*, species-specific orthogroups were most frequently categorized as being involved in post translational modifications (73/12.9% and 101/10.1%, respectively) and signal transduction (42/7.4% and 68/6.8%) as well as the transport/metabolism of carbohydrates (41/7.3% and 76/7.6%) and amino acids (31/5.5% and 56/5.6%). Additionally, *P. calceolata*-specific orthogroups were abundant in energy production/conversion (‘C’; 31/5.5%), while *Au. lagunensis* were abundant in replication, recombination, and repair (‘L’; 56/5.6%) (Extended Data Figure 9A, Supplementary Information). Overall, these data speak to many pelagophyte species-specific differences in gene content related to post-translational modifications and signal transduction, as well as carbohydrate, lipid and amino acid transport/metabolism – some of which can be attributed to prokaryote-derived genes.

### LGT has impacted pelagophyte genomes

1,077 pelagophyte orthogroups/singletons were inferred to be the product of recent LGT events from prokaryotes/viruses (i.e., absent in other stramenopiles and limitedly distributed across other algal groups) (Figure 4; all trees are available in Supplementary Information File 1). The proportion of orthogroups in any given species attributed to recent LGT is similar, ranging from 3.5% in *P. calceolata* to 3.8-4.0% in *Ac. anophagefferens* and *Au. lagunensis*, respectively. The impact of recent LGTs on species-specific orthogroups varied: 2.8% in *P. calceolata*, 4.7% in *Ac. anophagefferens*, and 5.3% in *Au. lagunensis*. While other mechanisms have no doubt contributed to protein-coding gene differences between examined pelagophyte species, over 60% of the inferred recent LGTs are species-specific, including over 50% of those in *Ac. anophagefferens*, suggesting that LGT has contributed to pelagophyte biology over recent evolutionary timescales.

In *Ac. anophagefferens*, 368 of 453 recent, species-specific LGTs reside in the pangenome core (81%); they are bona fide bacterial genes integrated into the genomes of all five strains and are not present in the other two pelagophyte species. Within the accessory genome (discussed above), the overall contribution of LGT to a given strain was highly similar (3.9 +/− 0.1% of orthogroups). 63/113 accessory genome LGTs (55.8%) were found in 4/5 strains, likely resulting from ancestral LGT events and subsequent differential gene loss within specific *Ac. anophagefferens* strains. As noted, a small fraction of strain-specific genes within *Ac. anophagefferens* (17) were attributed to recent LGT. Around 80% of predicted LGTs in *Ac. anophagefferens* and ∼95% of LGTs in *P. calceolata* and *Au. lagunensis* were found to be actively transcribed (TPM>1), suggesting they are functional.

Predicted recent prokaryotic LGTs within pelagophytes are associated with a wide variety of KOG/COG functional categories, most frequently with carbohydrate, secondary metabolite, amino acid or lipid transport and metabolism, as well as post-translational modifications (Extended Data Figure 9B). In *Ac. anophagefferens*, species-specific LGTs make up 40-50% of such transfers in most KOG/COG categories related to transport and metabolism as well as post-translational modifications, suggesting that recent LGTs have played an outsized role in this organism’s metabolic capacities and ability to regulate protein activity, localization and interactions. Recent LGTs within each pelagophyte species were most significantly enriched in GO terms associated with metabolic processes, including the metabolism of organo-nitrogen containing compounds, small molecules, carbohydrates and general biosynthesis, as well as molecular functions related to catalytic activities, including numerous transferases, hydrolases, oxidoreductases and lyases (Supplementary information). While most associated GO terms were not significantly enriched in species-specific LGTs (Supplemental Information Tables 3-4, 7-12), many of their predicted functions could have a key role in algal-bloom dynamics and pelagophyte abundance in the open ocean (Figure 3B-C, Extended Data Figure 10, Supplementary Information). This includes a thymidine kinase in *P. calceolata* (Figure 5A); a bacterial-type biotin transporter (BioY) in *P. calceolata* (Figure 5B); a xylulokinase (XylB) in *Au. lagunensis* (Figure 5C); a predicted alpha-glucuronidase (AguA2) associated with xylan catabolism in *Ac. anophagefferens* (Figure 5D); a polygalacturonase gene (Pgu1) associated with pectin degradation in *Ac. anophagefferens* (Figure 5E); and a tellurite-resistance/dicarboxylate transporter (TehA) in *Ac. anophagefferens* (Figure 5F) (see Supplemental Information).

**Figure 5.**
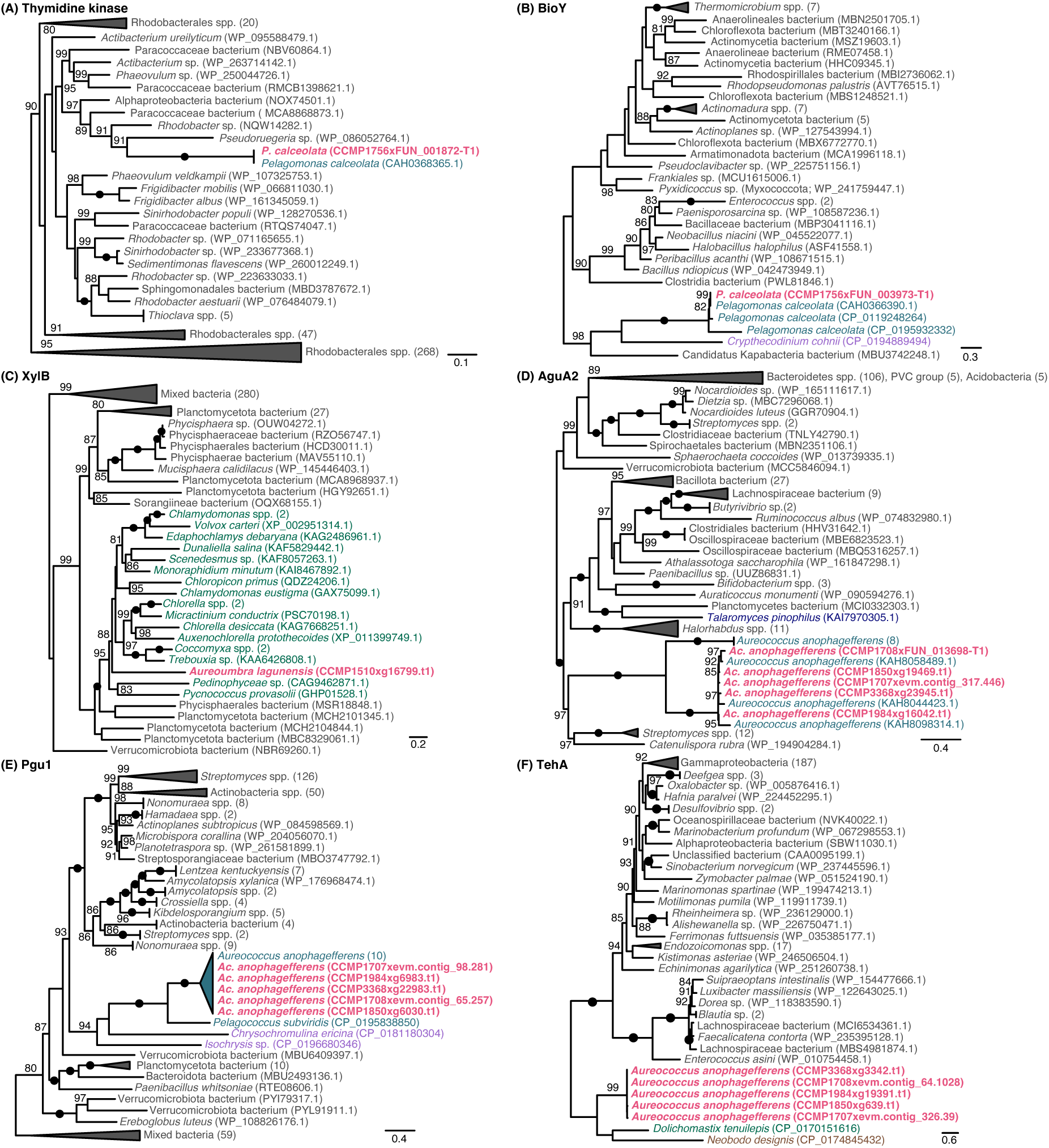
Examples of recent LGTs in pelagophyte algae of prokaryote/viral origin. (A) a thymidine kinase in *P. calceolata,* (B) a bacterial-type biotin transporter (BioY) in *P. calceolata,* (C) a xylulokinase (XlyB) in *Au. lagunensis,* and a (D) alpha-glucuronidase (AguA2), (E) polygalacturonase gene (Pgu1) and (F) tellurite-resistance transporter (TehA) in *Ac. anophagefferens.* Phylogenies were inferred using the best-fitting model (Methods) and are midpoint rooted. Only support values for nodes > 80% (UFBoot2) are shown; black circles indicate 100% support. OTUs are colored according to their taxonomic group as follows: Bacteria=grey, Viridiplantae=green, Pelagophyceae=teal, Haptophyceae/Alveolata=purple, Fungi=blue, Excavata=brown. Sequences derived from data generated in this study are shown bold and in pink.

Where have the pelagophyte-specific LGTs come from? Putative donors map to a wide range of bacterial lineages, including members of the Gammaproteobacteria, Alphaproteobacteria, Deltaproteobacteria, Firmicutes, Actinobacteria, and FCB-group. In some instances, the amino acid identity between the predicted donor and the pelagophyte gene/protein is >70% (and thus break the so-called ‘70% rule’^73^) (e.g., a thymidine kinase in *P. calceolata*, Fig. 5A, and a T4-type lysozyme in *Au. lagunensis;* see Supplemental Information). A handful of recent LGT trees have viral/phage homologs present (most noticeably in strain-specific genes); however, the directionality of transfer in these cases is typically unclear. Notably, although we have not specifically searched for it, eukaryote-to-eukaryote gene transfer appears to have play an important role in the evolution of at least some of the bacterial genes identified in our LGT detection pipeline, as many of our trees contain OTUs from dinoflagellates and/or haptophyte species that branch together with pelagophytes (e.g., Figure 5B/C/E, Supplemental Information). Reliably inferring the timing and order of these LGTs from bacteria to eukaryotes and between eukaryotes is at present not possible.

## DISCUSSION

We have shown that the genomes of extremely closely related strains of the HAB-forming pelagophyte species *Ac. Anophagefferens* can differ by almost two Mbp and have substantially different gene sets linked to their ecological success. Together with results gleaned from genomic investigations of other pelagophytes^12–14^, haptophyte and chrysophyte algae^8–10^, and other eukaryotes such as amoebae^74^ and fungi (e.g., ref. ^6^), our data underscore the significance of the pangenome concept for eukaryotes^7,19^. Fundamental questions about the processes underlying intra-species genetic variation across the eukaryotic tree nevertheless remain.

Differential gene loss and/or duplication of core genes followed by divergence / differential retention of paralogs appears to have played a role in shaping the accessory genome of *Ac. anophagefferens*, as in other eukaryotes (see ref. ^7^ and references therein). A significant number of accessory genes in the strains examined herein have no obvious homology to known protein-coding genes in functional or sequence databases, suggesting that they are either poorly characterized, highly divergent, or very recently evolved. While *de novo* gene birth was initially thought to be unlikely, this is not the case (see refs.^75,76^ and references therein). Determining what fraction of orphan genes within the pelagophyte accessory genome has evolved *de novo* from non-coding regions will require a detailed synteny-based analysis of our newly sequenced genomes, but we should remain open to the possibility of gene evolution ‘from scratch’ as a mechanism contributing to niche adaptation.

While accessory genomes in bacteria have been largely attributed to LGT^77^, evolutionary studies of eukaryotic pangenomes suggest that the impact of foreign gene acquisition is minimal (e.g., <0.1% of accessory genes in haptophytes^8,9^). Intra-species gene content variation in *Ac. anophagefferens* does not appear to be largely driven by recent prokaryotic LGTs (∼2.6% of accessory genes), but recent gene transfers have nevertheless contributed to a variety of processes and functions that are likely adaptative, as seen in other algae. This includes bacterium-derived accessory genes involved in lipid metabolism (e.g., ceramide metabolism and polyketide synthases), xylan catabolism (e.g., an alpha-glucuronidase), pectate and aldehyde lyase activity, *de novo* IMP biosynthesis, DNA repair and stress response related processes, and arsenic detoxification. Some accessory genes were found to have close homologs in phages and dsDNA viruses; the viral contribution to strain-specific genes was proportionately higher than the rest of the accessory genome, and we found evidence for both individual protein-coding genes and larger genomic regions unique to CCMP1850 that are clearly derived from AaV, the virus known to infect *Ac. anophagefferens*^72^ (Extended Data Figure 8). This is interesting given the differential susceptibilities of such strains to viral infection in the lab^78^. Viruses thus appear to have played an ongoing, albeit limited, role in the evolution of individual strains of *Ac. anophagefferens*.

Over half of the ∼1,100 LGTs identified within the Pelagophyceae are species-specific, including many genes with functional predictions relevant to HAB proliferation and niche adaptation. Whereas some LGTs provided unique functions to a given pelagophyte species (e.g., xylose metabolism, pectin degradation, nucleotide salvage and biosynthesis pathways), others were associated with similar functions/domains predicted within more ancestral genes (e.g., arsenic detoxification, DNA alkylation damage repair, repair of oxidatively damaged proteins and DNA, DNA mismatch repair); these LGTs, as well as those that were duplicated after being acquired, could provide increased metabolic efficiency and flexibility in specific environmental conditions and/or have evolved more specialized or new roles within these algae. Overall, the proportion of orthogroups attributed to recent LGT in each species were similar, ranging from ∼3.5% in *P. calceolata* to ∼4.0% in both *Ac. anophagefferens* and *Au. lagunensis.* This is greater than the ∼1% average contribution of LGT to eukaryotic genes suggested by Van Etten and Bhattacharya^18^ and that reported in previous stramenopile algal genomes (1.09%-2.13%^79,80^). Two of these LGTs – a thymidine kinase in *P. calceolata* and a T4-type lysozyme in *Au. lagunensis* – share >70% amino acid sequence identity with their closest bacterial homologs (72% and 79%, respectively), thereby breaking the proposed ‘70% rule’ for LGT in eukaryotes^73^.

While the majority of LGTs were acquired from diverse bacterial sources directly that plausibly coexist in their environment (with minor contributions from archaea or viruses), approximately one-third of predicted recent LGTs appear connected to additional gene transfers of bacterial genes between eukaryote species. Within these more complex gene transfer scenarios, pelagophyte homologs were often closely related to subsets of dinoflagellate and/or haptophyte species, and occasionally to some green algae. Altogether, this strongly suggests that eukaryote-to-eukaryote LGT is contributing to the spread of bacterial genes to algae. Studies on the species composition of some *Ac. anophagefferens* HAB locations have identified the presence of a variety of dinoflagellate and haptophyte species, including some associated with HAB formation^81^. Their co-occurrence with pelagophytes suggests LGT between these distinct eukaryotic lineages is plausible.

The dynamics of HABs in coastal ecosystems are complex^15,82^. In addition to viral predation, HAB-forming species interact with other algae, protists, and photosynthetic and heterotrophic prokaryotes, and experience fluctuating nutrient and light levels, osmotic and oxidative stress, and the presence of xenobiotic compounds in anthropogenically modified environments such as those inhabited by the pelagophytes *Ac. anophagefferens* and *Au. lagunensis*. Recent LGTs appear to help these species cope with various HAB related environmental stresses and increase their metabolic flexibility, allowing them to access organic compounds and more effectively rely on heterotrophic metabolism. For example, we identified LGT-derived genes associated with biotin transport and synthesis in *P. calceolata* and *Au. lagunensis,* respectively, and enzymes related to carbohydrate, phosphate and nitrogen-containing compounds in all examined species. In *Ac. anophagefferens*, the 368 predicted LGTs in its core genome that were not identified in the other pelagophytes (Fig. 4) are dominated by orthogroups with metabolism- and transport-associated functions, suggesting that foreign gene acquisition by ancestral strains facilitated the rise to prominence of today’s HAB-forming strains. A greater proportion of pelagophyte LGTs encode proteins that are secreted than do vertically inherited genes; such proteins presumably facilitate host-environment interactions, nutrient acquisition, or act as toxins. Overall, our study clearly shows that LGT has continuously and consistently contributed to the evolution of these three pelagophyte species, including having a limited impact on strain-specific genes within *Ac. anophagefferens,* and provides further support for the notion that LGT contributes meaningfully to eukaryote genome biology and evolution.

## Acknowledgements

This research was supported by an NSERC Discovery Grant (RGPIN-2019-05058) awarded to JMA. We thank M. Dlutek, B. Curtis, G. Filloramo and J. Jerlström-Hultqvist for extensive technical and bioinformatic support. SJS was support by graduate scholarships from NSERC (CGS-D) and the Killam Trusts. ML and CM were supported by NSERC USRA summer scholarships.

## Data Availability

BioProject IDs for pelagophyte genome sequence and assemblies are as follows: PRJNA488217 (CCMP1984), PRJNA690688 (CCMP3368), PRJNA690542 (CCMP1850), PRJNA690695 (CCMP1707), PRJNA690699 (CCMP1708), PRJNA690709 (CCMP1510), PRJNA690716 (CCMP1756). Raw sequence datasets are available in the SRA repository under accession numbers: SRR13386523 (CCMP1984), SRR13386499 (CCMP3368), SRR13386516 (CCMP1707), SRR13386522 (CCMP1708), SRR13378193 (CCMP1850), SRR13386728 (CCMP1510), and SRR13386796 (CCMP1756).

---

**Extended Data Figure 1.**
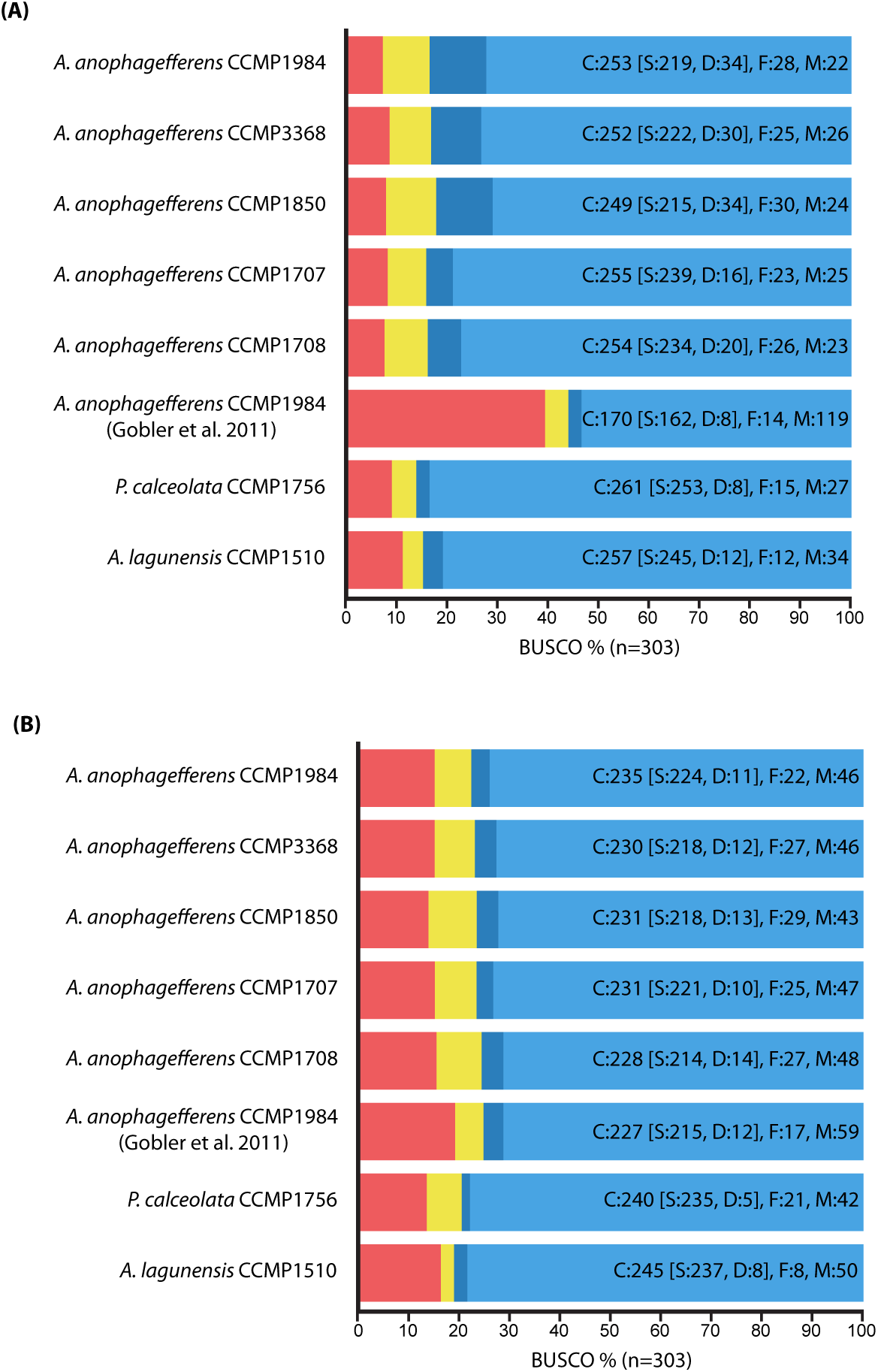
Benchmarking Universal Singly Copy Orthologs (BUSCO) summaries for newly sequenced pelagophyte (A) protein-coding gene predictions and (B) genomes. BUSCO assessment was performed against the eukaryotic dataset (v3; eukaryota_odb9; n=303). BUSCO assessment of the original *Aureococcus anophagefferens* CCMP1984 genome (Gobler et al., 2011) is shown for reference. C=complete, S=single copy, D=duplicated, F=fragmented, M=missing.

**Extended Data Figure 2.**
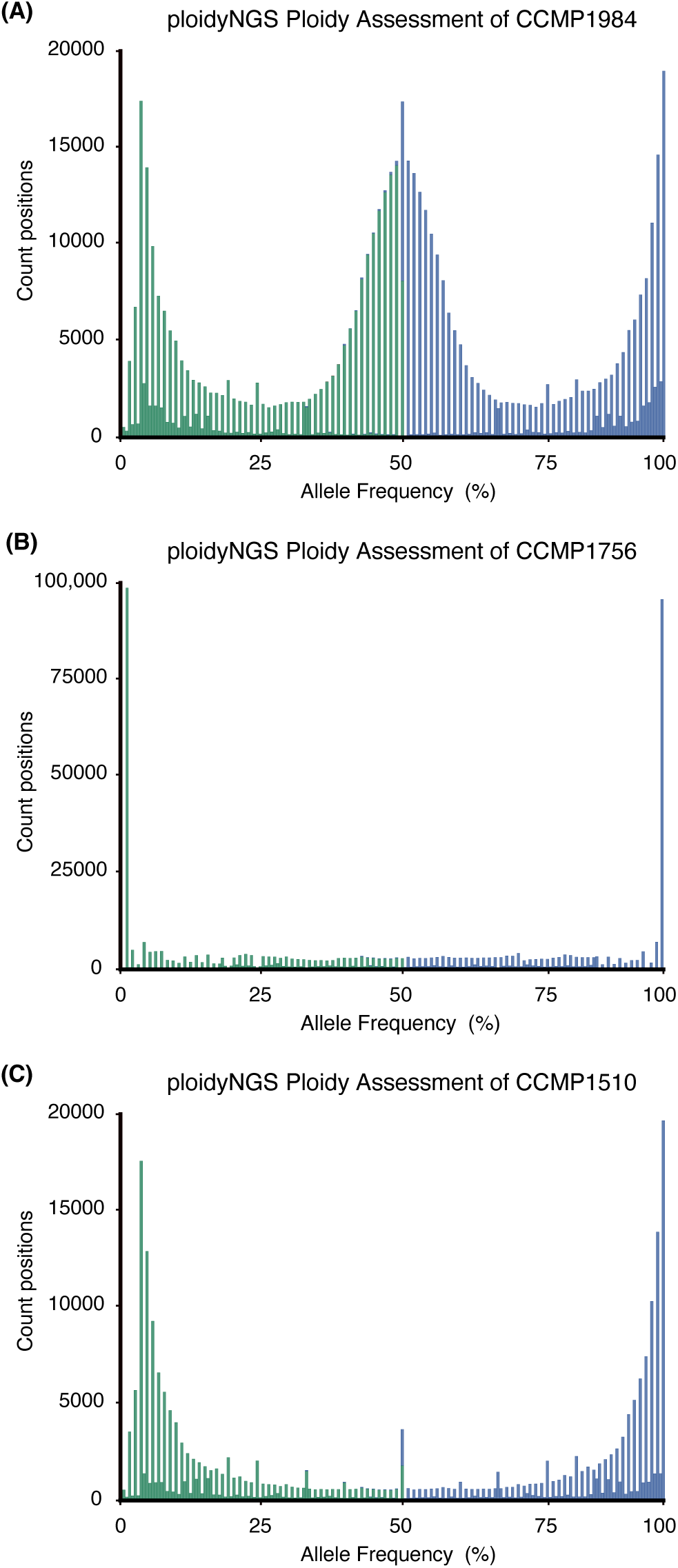
Ploidy assessment of all pelagophyte genomes based on ploidyNGS output. (A) *Aureococcus anophagefferens* CCMP1984, (B) *Pelagomonas calceolata* CCMP1756 and (C) *Aureoumbra lagunensis* CCMP1510. For simplicity, only one strain of *Ac. anophagefferens* is shown, but is reflective of results in all five strains and results from analysis using nQuire.

**Extended Data Figure 3.**
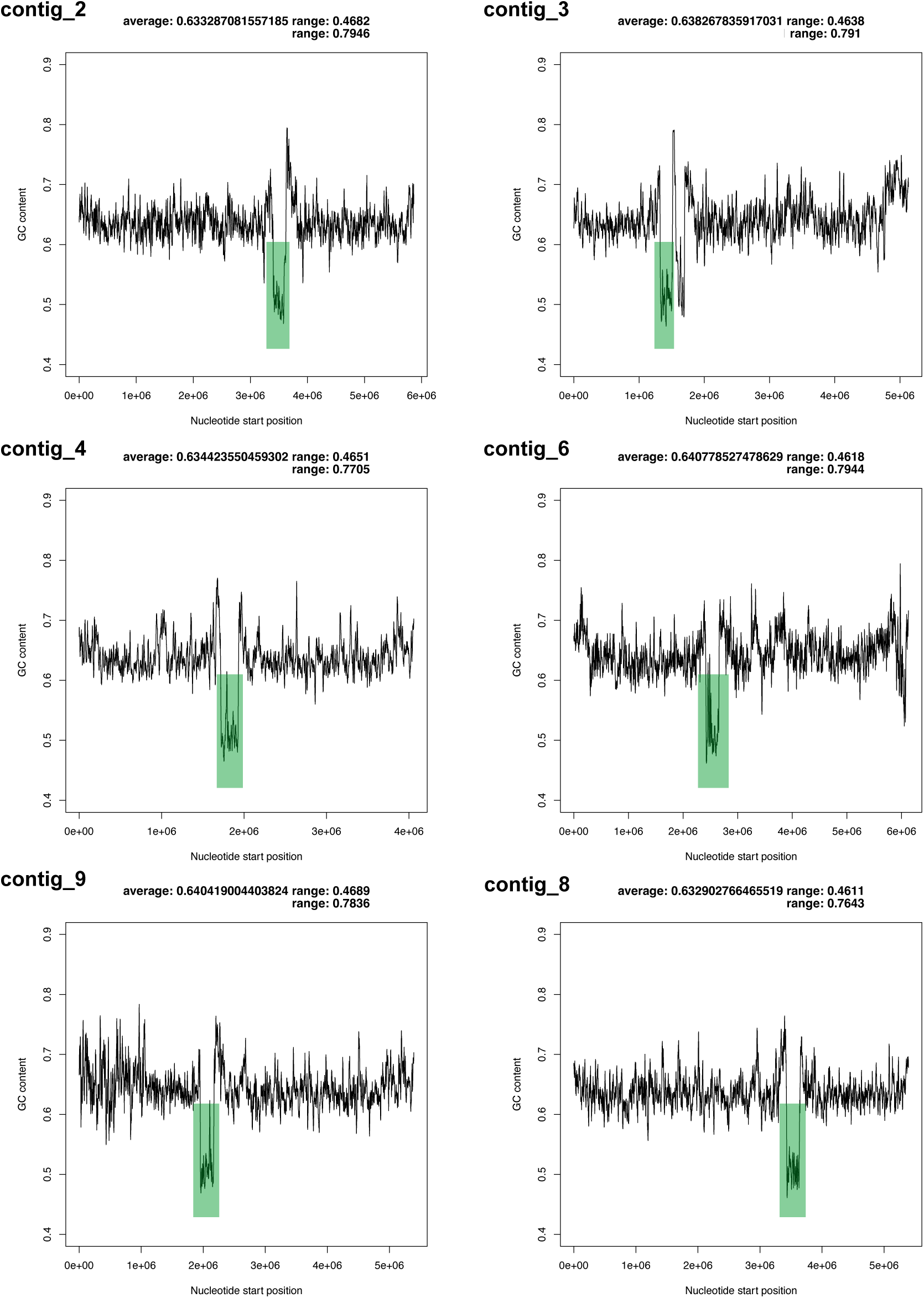
GC content sliding window analysis of all six fully resolved *Pelagomonas calceolata* CCMP1756 contigs. Average GC content across each contig is shown in sliding windows of 10 kbp. The overall average and range of GC content across each contig is shown. Putative centromere regions in each contig are highlighted by green boxes.

**Extended Data Figure 4.**
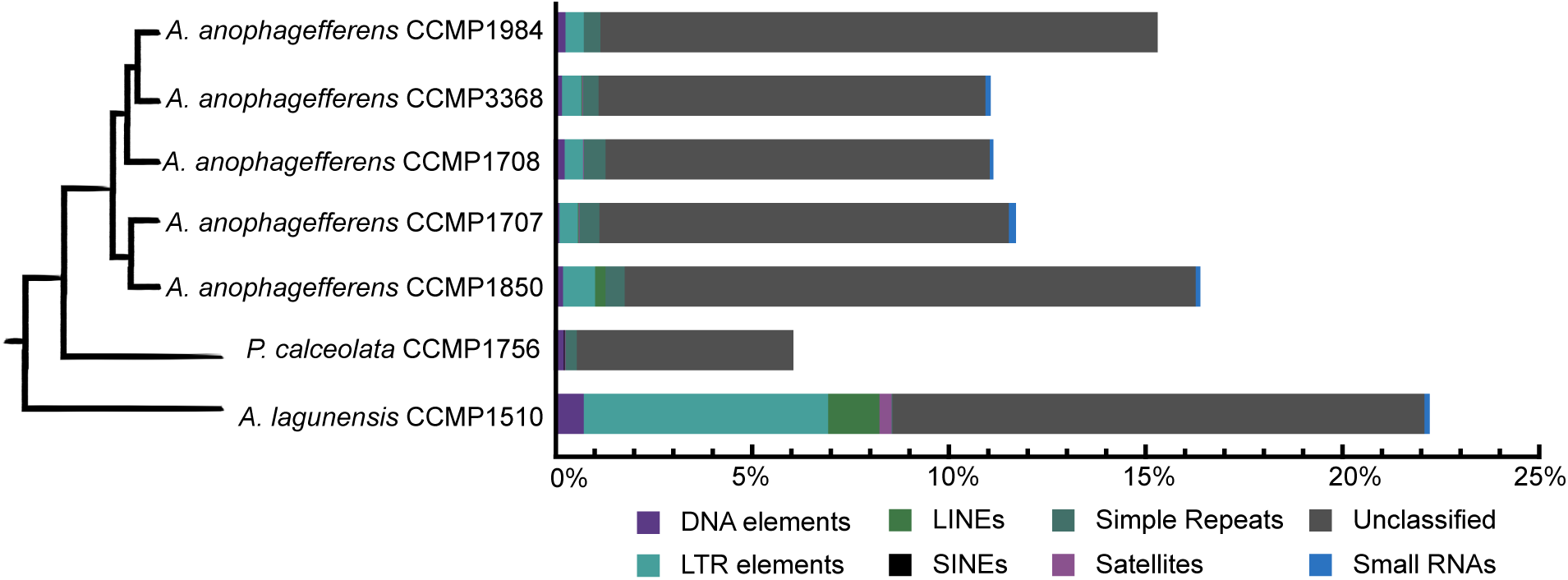
Genome repeats identified in all pelagophyte genomes. Repeat content is shown as a percentage of the total genome assembly for each pelagophyte. Different repeat classifications as identified by RepeatMasker (v4.1.3) are shown by corresponding colours within each column. The schematic phylogeny shown is as inferred using multi-gene phylogenies. For a more detailed breakdown of repeat content, see Supplementary Information Table 1.

**Extended Data Figure 5.**
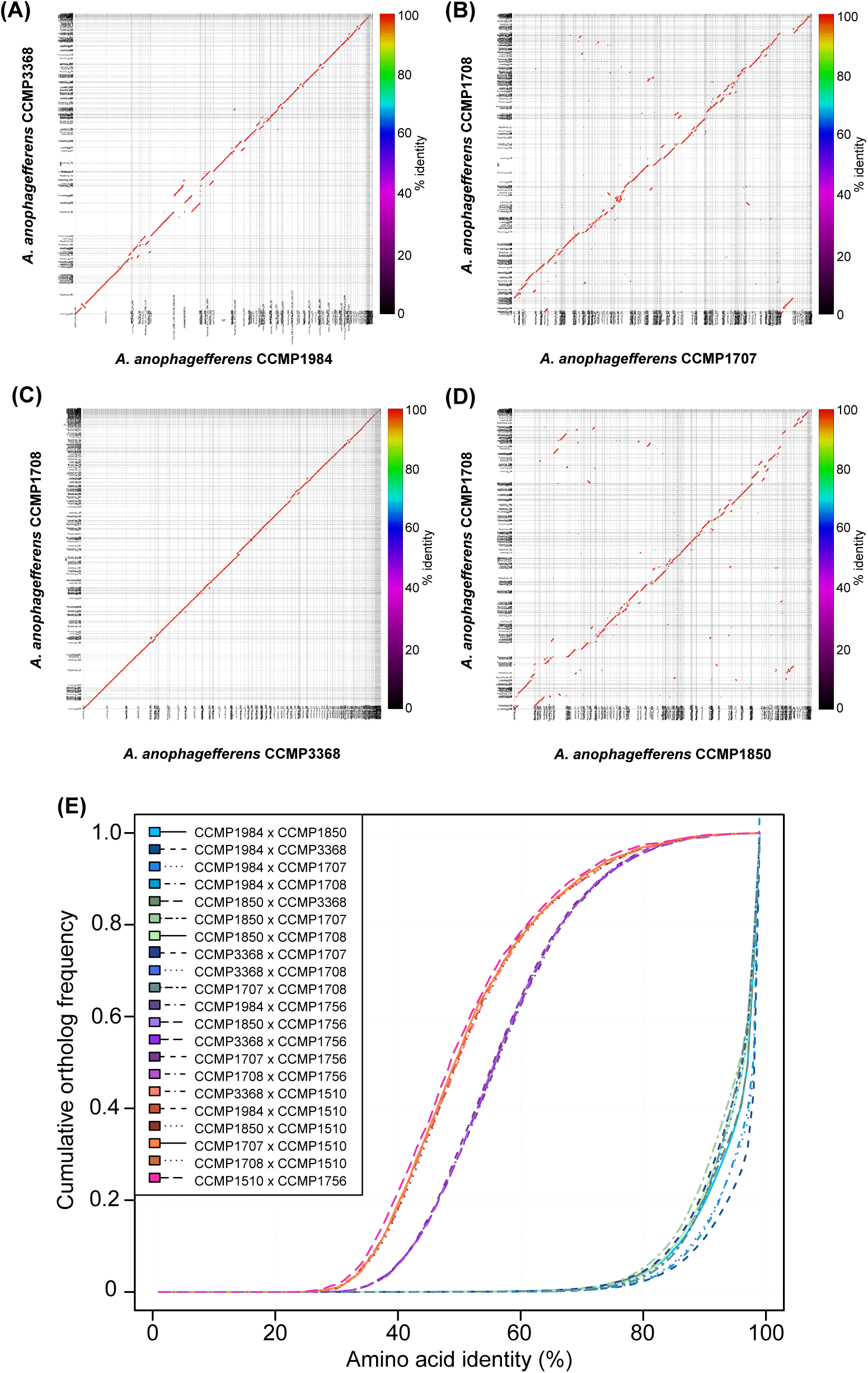
Genome similarity based on whole genome alignments and protein-coding gene identity between strains of *Aureococcus anophagefferens* and other pelagophyte species. **(A-D)** Mummer plots of pairwise, whole genome nucleotide-based alignments of haploid genome assemblies for select *Ac. anophagefferens* strains. Colours of matching regions indicate percent identity of the nucleotide alignment, where red indicates >97% identity. **(A)** *Ac. anophagefferens* CCMP1984 vs. *Ac. anophagefferens* CCMP3368, **(B)** *Ac. anophagefferens* CCMP1707 vs. *Ac. anophagefferens* CCMP1708, **(C)** *Ac. anophagefferens* CCMP3368 vs. *Ac. anophagefferens* CCMP1708, **(D)** *Ac. anophagefferens* CCMP1850 vs. *Ac. anophagefferens* CCMP1708, **(E)** Pairwise ortholog similarity between *Ac. anophagefferens* strains and pelagophyte species based on percent identity of reciprocal best blast hits of protein-coding genes. Comparisons between *Ac. anophagefferens* strains are shown in blue/green colours, while purple indicates comparisons between *Ac. anophagefferens* and *Pelagomonas calceolata*, and orange or pink indicates comparisons between *Ac. anophagefferens* or *P. calceolata* with *Aureoumbra lagunensis*, respectively.

**Extended Data Figure 6.**
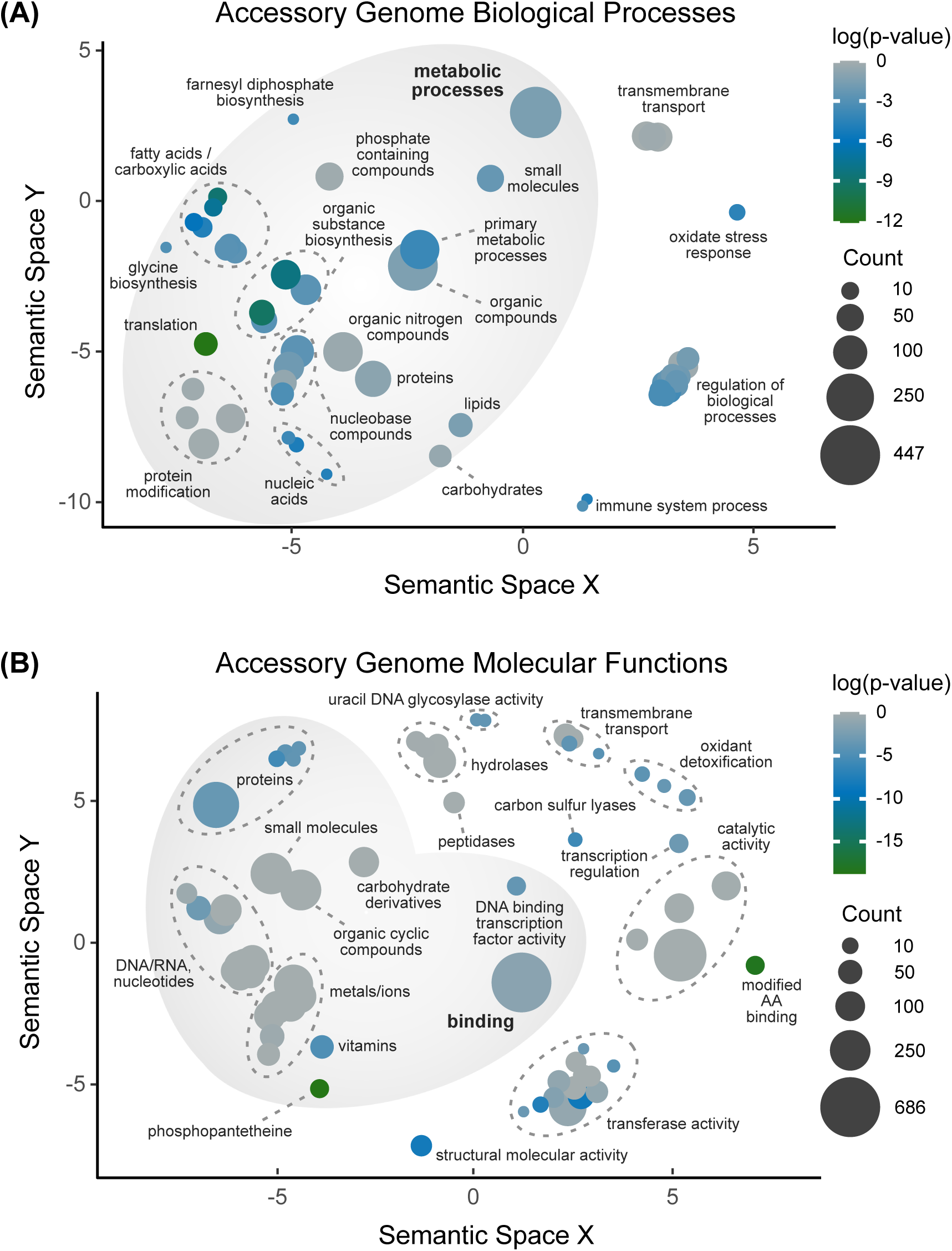
REVIGO based scatter plots of GO terms associated with the *Aureococcus anophagefferens* accessory genome. GO terms associated with the entire accessory genome that are related to **(A)** biological processes, and **(B)** molecular functions. The number of accessory orthogroups with a GO term and its degree of statistically significant enrichment (log p-value) is reflected by the bubble size and colour, respectively. Labeling has been summarized for clarity; complete terms are available in Supplementary Table 5-6.

**Extended Data Figure 7.**
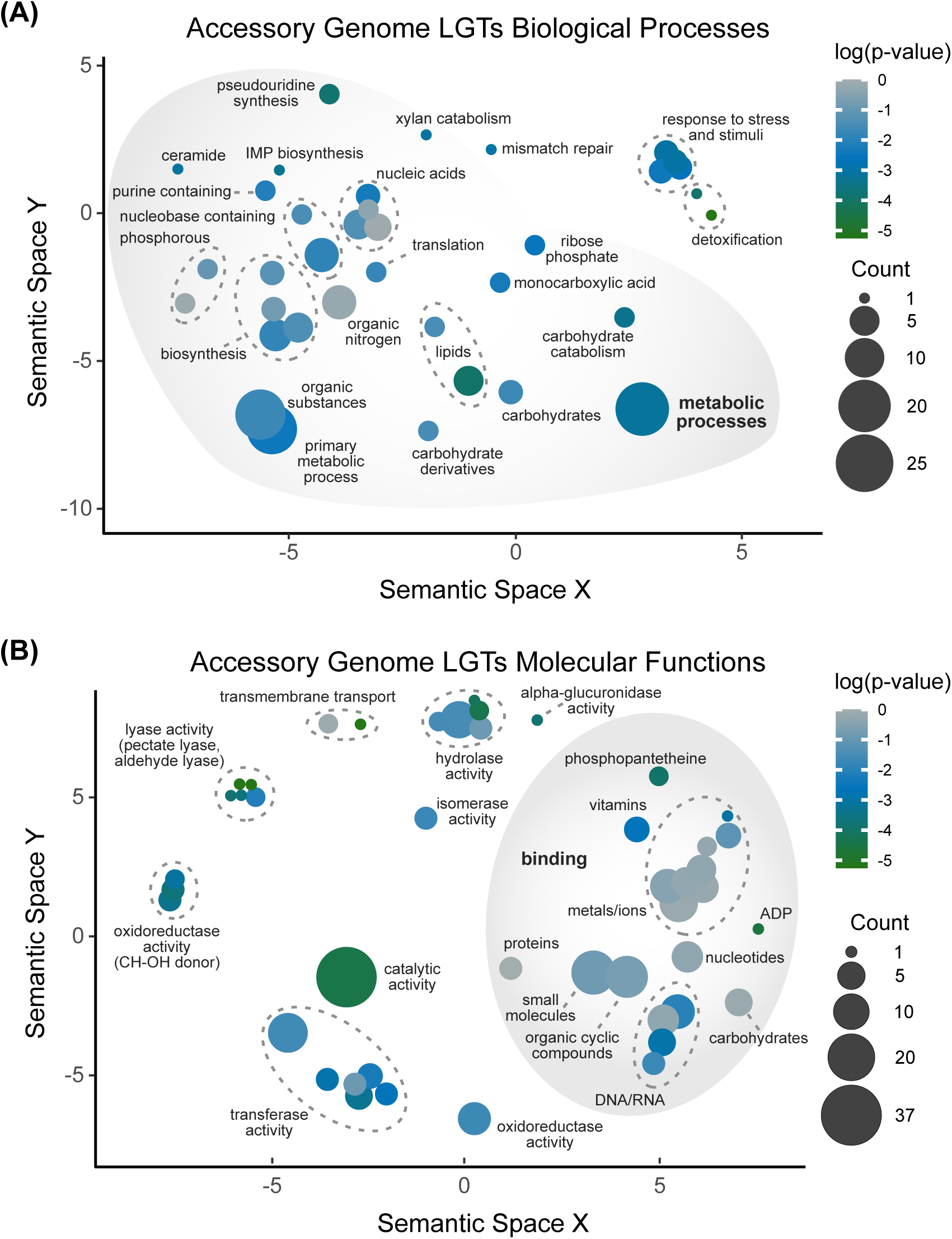
REVIGO based scatter plots of GO terms associated with recent LGTs the *Aureococcus anophagefferens* accessory genome. **(A)** GO terms associated with the recent LGTs within accessory pangenome that are related to biological processes, and **(B)** GO terms associated with recent LGTs within the accessory pangenome that are related to molecular functions. The number of LGTs with a GO term and its degree of statistically significant enrichment (log p-value) is reflected by the bubble size and colour, respectively. Labeling has been summarized for clarity; complete terms are available in Supplementary Table 7-8

**Extended Data Figure 8.**
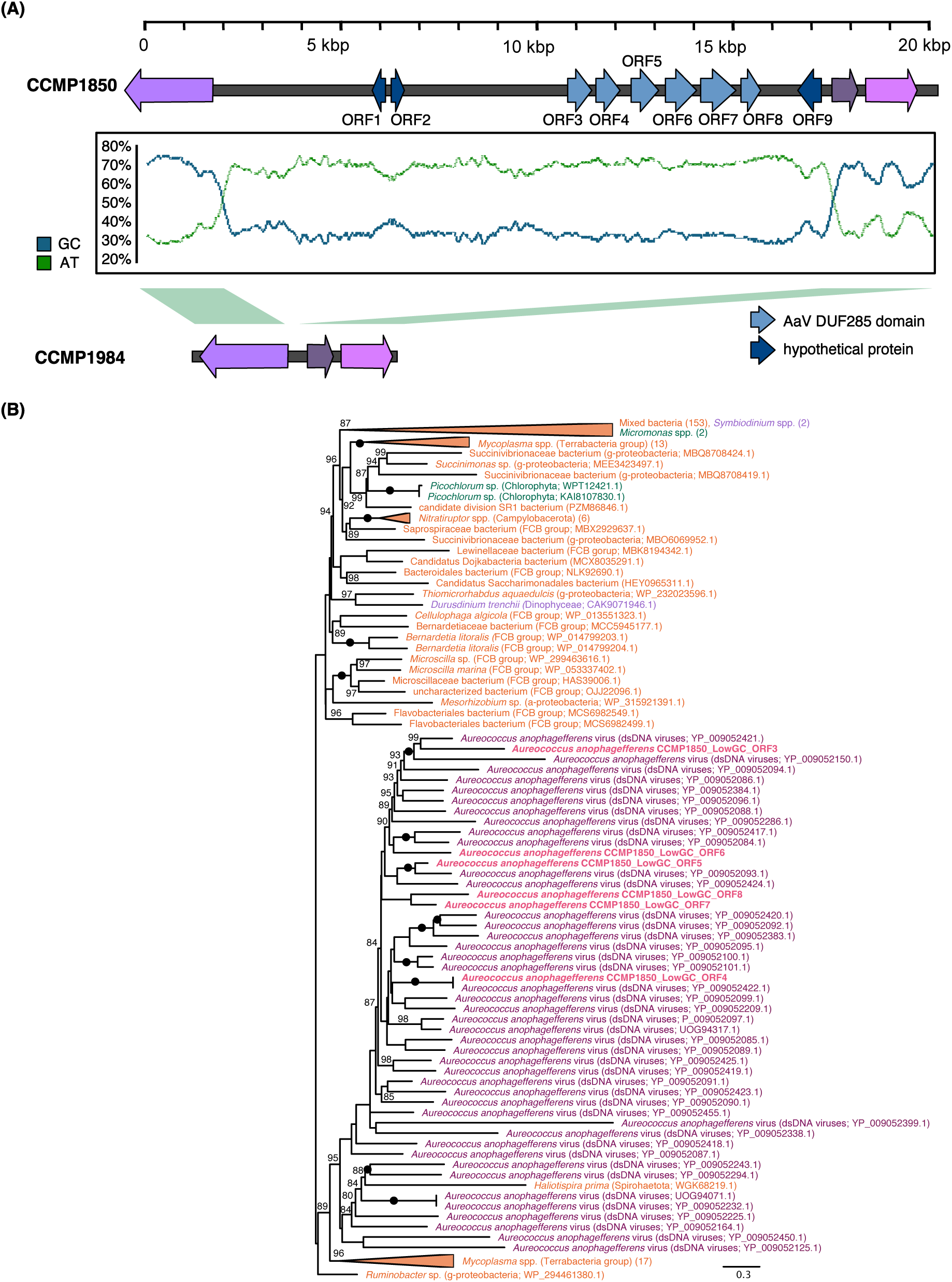
A unique low GC-content region in *Aureococcus anophagefferens* CCMP1850 of viral origin. **(A)** A schematic representation of a low GC content, unique region in the genome of CCMP1850 (top) and it syntenic region in other *Ac. anophagefferens* strains (below). Protein-coding genes predicted in this region are shown to scale along the contig map with block arrows. Purple/pink blocks indicate genes with corresponding homologs in other *Ac. anophagefferens* strains, with different shades representing different orthologs. Blue blocks indicate predicted ORFs (>100 AA) within the region unique to CCMP1850, with dark blue representing hypothetical proteins and light blue specifically encoding DUF285 domains. The middle plot shows the GC (blue) vs AT (green) content mapped across the CCMP1850 contig in a sliding window with increments of 500 bp **(B)** A maximum likelihood phylogeny of ORFs containing the DUF285 domain (leucine rich repeat surface protein) and all homologs in nr with an e-value <1e-5. The phylogeny was inferred using 269 OTUs and 328 sites under the model WAG+R8 (as determined to best fit the data according to BIC) with 5,000 UFBoot2 replicates and rooted in midpoint. Only UFBoot2 support values > 80% are shown; a black circle indicates a node was maximally supported. Sequences are colored according to major eukaryotic group they are affiliated with as follows: Alveolata = purple, Viridiplantae = green, Bacteria = orange and viruses/phages = dark purple. Sequences derived from data generated here are shown bold and in pink. The scale bar represents 0.3 substitutions per site.

**Extended Data Figure 9.**
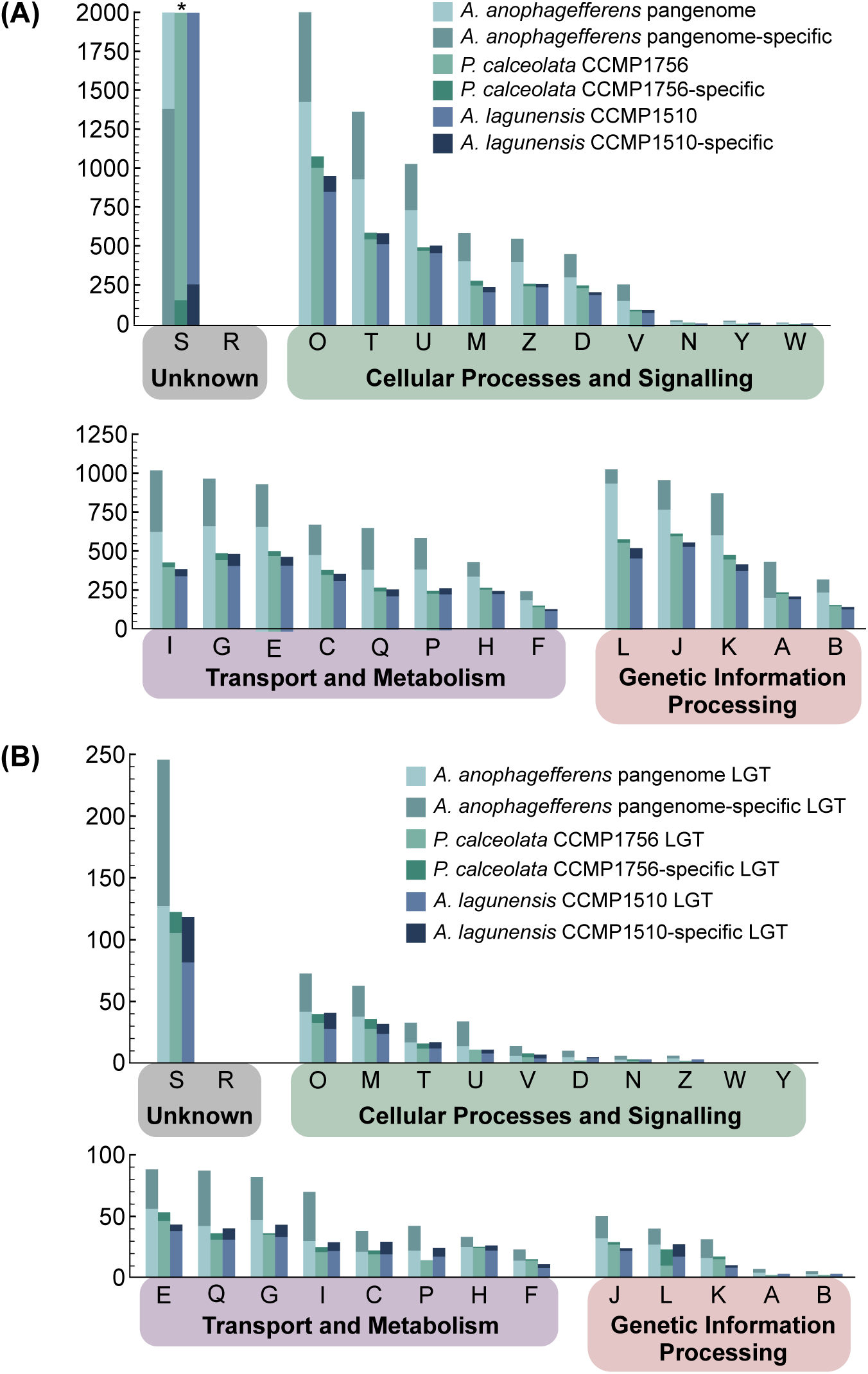
KOG/COG categories of orthogroups in pelagophyte genomes (A) and in predicted recent LGTs in each species (B). Within a column, the proportion of species-specific genes/orthogroups term is indicated within a column (darker shading). The ‘*’ in panel (A) indicates bars have been cut off at a maximum amount for readability. The actual counts for category ‘S’ are as follows: *Ac. anophagefferens*: 4785, *P. calceolata*: 2419, and *Au. lagunensis:* 2270.

**Extended Data Figure 10.**
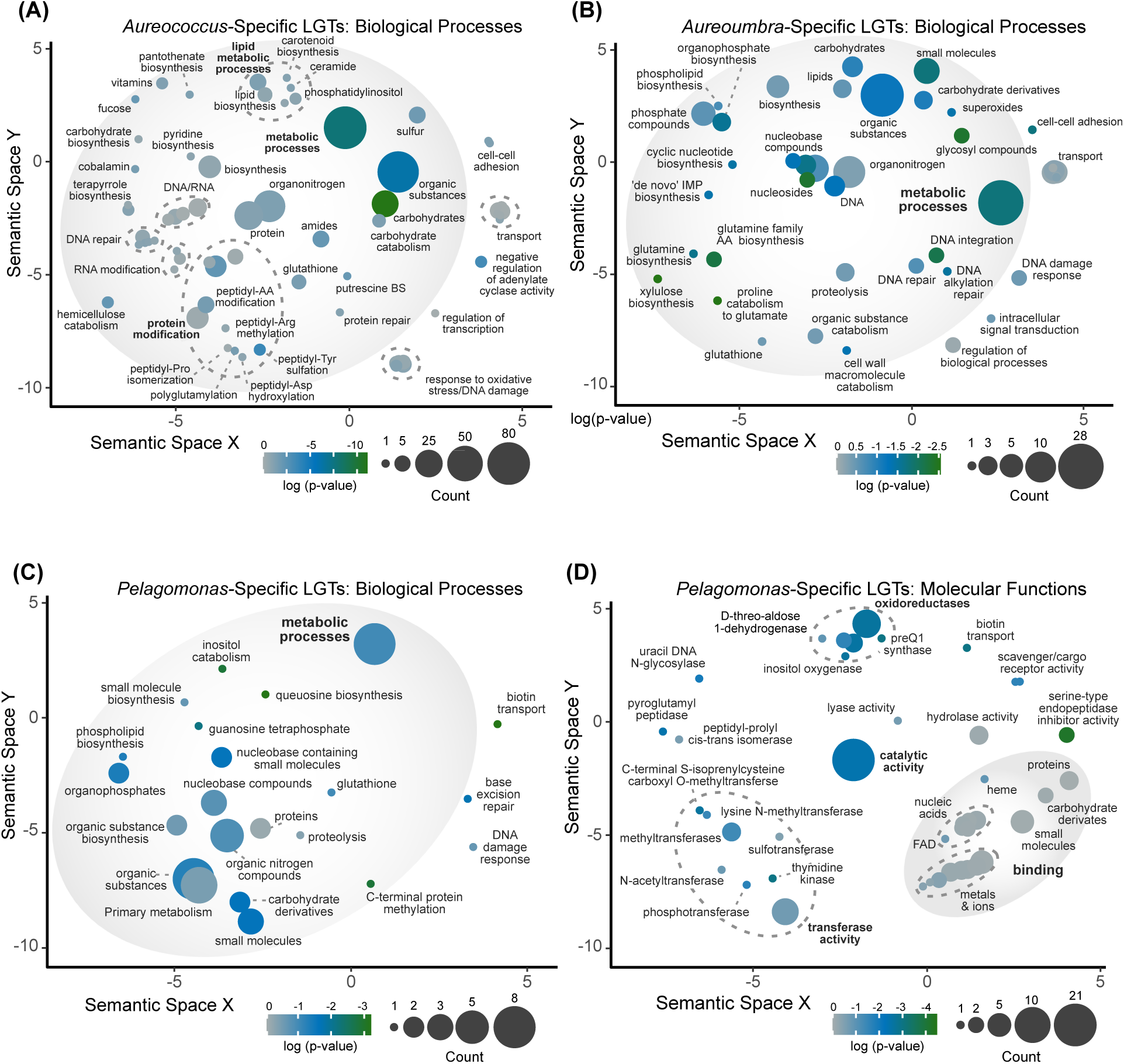
Additional REVIGO based scatter plots of GO terms associated with species-specific LGTs within the pelagophytes. GO terms associated with biological processes in (**A)** *Ac. anophagefferens*-specific LGTs (see also Figure 3), **(B)** *Au. lagunensis*-specific LGTs (see also Figure 3), and **(C)** biological processes associated with *P. calceolata-* specific LGTs, as well as **(D)** molecular functions associated with *P. calceolata-*specific LGTs. The number of LGTs with a GO term and its degree of statistically significant enrichment (log p-value) is reflected by the bubble size and colour, respectively. Labeling has been summarized for clarity; complete terms are available in Supplementary Tables 9-12.

