## Supplementary Information for "Pangenome biology and evolution in harmful algal-bloom-forming pelagophyte algae"

**The relationship between *Ac. anophagefferens* strains and pelagophyte species**

Previous studies of *Ac. anophagefferens* blooms have suggested there is little genetic variability in select marker genes between strains, suggesting that cryptic species do not exist within and between bloom sites[1,2]. Here, 18S rDNA copies in *Ac. anophagefferens* were found to have 98.9-100% identity within and between strains, with intra- and inter-strain similarity being similar. The exception was one or two divergent 18S rDNA copies in most strains that show only 92.5-96% identity to all other 18S rDNA copies; this lower identity is on par with the identity between *Ac. anophagefferens* and *Au. lagunensis* 18S genes (93-94%) and lower than the average identity to 18S genes in *P. calceolata* (97-98%). Although concerted evolution is thought to reduce polymorphisms in rDNA copies within a genome such that copies are largely homogenous[3], intragenomic variation in rDNA sequences have been reported for microbial eukaryotes including ciliates[4–6], foraminifera[7] and dinoflagellates[8], and some metazoans (e.g., refs. [9,10]).

*Ac. anophagefferens* strains CCMP1984, 1708 and 3368 have 11 rDNA copies located mostly near the end of one telomere containing contig; CCMP1850 and 1707 have 12 or 13 copies split across a few smaller contigs. Strong synteny was observed in the rDNA region of *Ac. anophagefferens strains*, apart from a ~0.6 Mbp chromosomal inversion involving the two most internal rDNA copies between CCMP1984 and the other strains. Notably, 18S rDNA in syntenic positions between strains did not necessarily have the highest identity or even 18S structural form to one another. In *P. calceolata*, two nearly identical copies of rDNA (99.8% identity) were found located in a subtelomeric region within ~10 kbp of one another. Five rDNA regions were identified near the ends of separate contigs in *Au. lagunensis*, with the 18S rDNA showing 98.8-99.3% pairwise identity and all containing a 420 bp intron previously predicted to be self-splicing[11]. A notable strain-level polymorphism in the 18S rDNA copies of *Ac. anophagefferens* is the presence of both intron-containing (903-975 bp intron) and intron-lacking versions, both of which were supported by RNA-seq and long-read sequencing data. Due to the intron boundaries predicted at a GC-AG splice site and lack of clear secondary structure similar to Group I or Group II intron elements, it seems likely a spliceosomal intron. An apparently unrelated 18S rDNA intron was identified in all copies of the 18S rDNA in *Au. lagunensis* that had no significant sequence similarity to the rDNA introns found in *Ac. anophagefferens*, that was previously determined to be a Group I intron with high similar to those in the 18S rDNA in red algae and present in *Au. lagunensis* due to LGT[11].

There was not enough signal within the 18S rDNA alone to infer the relationships between *Ac. anophagefferens* strains, as expected due to the gene's high sequence conservation. A maximum-likelihood multi-gene phylogeny of >1,300 individual protein-coding genes appeared to resolve the intra-species relationships, with all nodes maximally supported. Interestingly, strains isolated from either Great South Bay or West Neck Bay bloom sites did not branch together (although these bloom locations are only ~100 km apart). CCMP1984 and CCMP3368, which were isolated in consecutive years from the same bloom location, were found to branch together. Outside of this, no geographic or temporal patterns in strain relationships were observed.

Of note is the fact that while bootstrap values maximally support strain relationships in the maximum likelihood tree, gCF values were surprisingly low (<20%). This could be due to limited signal in the single gene trees attributable to high similarity between the strains' gene copies and/or conflicting signals in the single gene tree dataset due to processes like incomplete lineage sorting and recombination having occurred. While a handful of alternative topologies were observed frequently in the single gene trees, none of the alternative topologies were considered likely on the basis of AU tests and were rejected. Most topologies of the individual gene trees supported a divide between CCMP1707 and 1850 and the other strains. Further support for this relationship comes from mitochondrial genome identity[12] overall whole genome alignments between the strains, and from the 2-3X increase of strain-specific genes in the clade containing CCMP1850 and 1707 compared to the clade containing CCMP strains 1984, 3368 and 1708. One interesting phenotypic difference between CCMP1707 and 1850 compared to the other strains is that they were found to be unsusceptible or only partially susceptible to lysis by AaV, while both CCMP1984 and 1707 were highly susceptible to infection[13]. Closely comparing the protein-coding genes of susceptible versus non-susceptible strains has the potential to provide key insight into resistance to AaV infection and how *Ac. anophagefferens* HABs persist. A close comparison between the sister strains CCMP1707 and 1708 who were isolated from the same HAB yet show differing viral susceptibilities may be the most enlightening.

**The pangenome of *Ac. anophagefferens***

Based on the five strains of *Ac. anophagefferens* included here, a species pangenome was inferred that consists of 23,356 orthogroups, 18,949 (81.1%) that are core and 4,407 that are accessory and found in 1-4 of the strains. The addition of each strain to the pangenome assessment in stepwise fashion resulted in the addition of 200-300 new orthogroups (after the initial pairwise comparison where >800 unique orthogroups were found), incrementally increasing the proportion of the accessory pangenome by ~4%. This suggests that the pangenome is still 'open' and that inclusion of data from more strains will continually decrease the size of the core genome and increase the overall pangenome size. A few other studies have explored gene content variation between some *Ac. anophagefferens* strains[14–16]. On the basis of presence-absence of KEGG identifiers, Gann *et al.*[16] identified ~90% of protein-coding genes with KEGG annotations to be found in common between four *Ac. anophagefferens* strains. Studies involving transcriptomes from CCMP1850 under varying culture conditions found that only ~95% of its transcripts aligned to the CCMP1984 reference genome, suggesting that fine-scale variation exists between at least these two strains[15]). Similar levels of variation were estimated between CCMP1984 and 1850 here, who shared ~95.5% of their respective orthogroups. Similar transcriptome mapping approaches found that only ~85% of transcriptome reads derived from a Chinese strain of *Ac. anophagefferens* mapped to the reference genome of CCMP1984[14]. This is lower than the proportion of predicted protein-coding genes that overlap between any pair of strains considered here, indicating that the diversity within *Ac. anophagefferens* genomes is not fully represented by comparisons of the geographically similar strains available in culture collections.

### *Predicted functions of strain-specific genes*

Within the *Ac. anophagefferens* pangenome, 4.05% of orthogroups/singletons were found to be strain-specific, with numbers ranging from 113 in CCMP1984 to 306 in CCMP1850. A small number of strain-specific genes/orthogroups (17/945) were determined to be recent LGT events from prokaryotes/viruses. Generally, 1-3 singletons in a strain were attributed to recent LGT, with the exception of CCMP1850 where five recent LGTs were inferred. Overall, four strain-specific genes had best blast hits to viruses/virophages including: (i) a homing endonuclease in CCMP1707 (SINGLETON0007987) that had best blast hits to *Pithovirus* (33% identity) and branched with various viruses, phages and bacteria in a poorly resolved phylogenetic tree; (ii) a cytosine specific DNA methylase in CCMP1708 (N0_HOG0020680_2) that had best blast hits to *Erythorocytic necrosis* virus (57% identity) and was phylogenetic resolved within a clade of a large number of other dsDNA viruses and some bacteria; and (iii) a D5 N-terminal like domain containing gene in CCMP1850 (SINGLETON_559) that had best blast hits to Yellowstone lake virophages (48% identity) and branched specifically with an unclassified virus in corresponding phylogenies. Although none of these predicted recent strain-specific LGTs with strong viral affinities were found to be expressed under normal culture conditions, their presence in single strains suggests viruses – and not just *Ac. anophagefferens* specific virus – have continually and recently integrated genetic material into these algae.

Recent strain-specific LGTs not involving viruses included a variety of predicted functions and bacterial donor lineages. In CCMP1984, recent LGTs included a beta-ketoacyl synthase gene likely acquired from gamma-proteobacteria (45% identity) (SINGLETON_649), as well an aldo-keto reductase (SINGLETON_592) that interestingly branched within a clade comprised of a handful of bilaterian and red-algal aldo-keto reductases that was sister to planctomycete bacteria (39% identity). In CCMP1708, strain-specific LGTs included a predicted SET domain associated with protein lysine methyltransferases (SINGLETON_314) acquired from xanthomonadales (31% identity), and a low molecular weight phosphatase that branched with *P. calceolata* homologs and *Phycisphaerales* bacterium (43% identity) (N0_HOG0024117). One additional recent LGT was identified in CCMP1707 (SINGLETON_46) that has close homologs in a green alga (*Pyramimonas parkeae*), two chlorarachniophyte algae, and an uncharacterized metallo-hydrolase in actinobacteria (35% identity); the specific branching pattern observed is consistent with at least one eukaryote-to-eukaryote LGT between *Ac. anophagefferens* and *P. parkeae* after an initial bacterial-to-eukaryotic LGT, while the presence in some chlorarachniophyte species can be explained by EGT related to their green-algal derived plastid. Strain-specific recent LGTs in CCMP1850 were more common. These included: (i) a calcium ion binding protein (N0_HOG0024262) that has unclear bacterial origin but best blast hits to cyanobacteria (49% identity); (ii) a DSD1-family PLP dependent enzyme likely acquired from a Chloroflexota bacterium (55% identity) (SINGLETON_402); (iii) an HRDC-domain containing ATP dependent helicase with homology to RecQ annotated genes in gamma-proteobacteria (50% identity) (SINGLETON_389) that have involvement in UV-induced DNA repair (Fig. 3.8B); and (iv) an N-acetylglucosamine/arylsulfatase domain (N0_HOG0014446_2) that is a strain specific duplication of a core LGT from FCB-group bacteria (30% identity) (data not shown). This duplicated LGT shares 98% identity with the core LGT at the amino acid level but resides in a non-syntenic region of the genome.

In general, strain-specific genes not attributed to recent LGT were divergent from other orthogroups with similar annotations and/or were patchily distributed amongst related algae; phylogenetic patterns for most of these genes were unclear though sometimes indicative of more ancient prokaryotic LGTs that have been differentially duplicated or lost within select *Ac. anophagefferens* strains. Overall, strain-specific genes with associated GO terms were found to be involved in a variety of biological processes and molecular functions, but are largely associated with binding (65%) and/or catalytic activities (49%), mostly related to transferases (19%), hydrolases (14%) or oxidoreductases (9%). Some examples include: a molybdenum cofactor sulfurase (MOSC domain) in CCMP1708 (N0_HOG0020786), a phosphoglycerate mutase in CCMP3368 (SINGELTON_708), an ammonium transmembrane transporter in CCMP1707 (SINGLETON_61)  and an ethylene response element containing DNA-binding transcription factors in CCMP1850 (SINGLETON_525).

### *Predicted functions of Ac. anophagefferens accessory genes of LGT origin*

Within the accessory genome, 113s LGTs were identified of recent prokaryotic/viral origin (including those that were strains-specific and discussed above). 98/113 accessory genome LGTs had an associated KOG/COG term (Extended Data 9B); over a quarter of these, however, were assigned to 'unknown functions' ('S'). Approximately half of characterized KOG/COG terms were predicted within transport and metabolism functional categories, with those related to secondary metabolites ('Q'; 11), carbohydrates ('G', 10) and lipids ('I', 10) being the most abundant. Most other accessory genome LGTs were assigned to post-translational modification ('O', 7), signal transduction ('T', 4) or translation ('J', 7) KOG/COGs, the latter including mostly genes involved in translation regulation. Recent LGTs within the *Ac. anophagefferens* accessory genome were significantly enriched in GO terms for biological processes related to metabolism (19 orthogroups) (Extended Data Figure 7A). This includes those involved in: (i) lipid metabolism (5), including significant enrichment in ceramide metabolic processes; (ii) carbohydrate metabolism (3), which includes significantly enriched carbohydrate catabolism related terms (2) related to glycolytic processes or Xylan catabolism; (iii) protein metabolism

(4); and (iv) metabolism of nucleobase containing compounds (5), including *de novo* IMP biosynthesis. Other significantly enriched biological processes within the accessory genome include response to DNA damage (2), sulfur transport and arsenic detoxification.

GO terms related to molecular functions of recent LGTs within the accessory genome were most significantly enriched and abundant in catalytic activities (37, 66%) (Extended Data Figure 7B). This broadly included several predicted transferases (13), hydrolases (10), oxidoreductases (9), isomerases (1) and lyases (2). Recent LGTs within the accessory genome were significantly enriched in specific types of methyltransferases. This included a rRNA methyltransferase (YqxC) acquired from alpha/beta-proteobacteria (51% identity) that was absent in CCMP1707, a methyltransferase type 11 gene found only in CCMP1707 and 1708 (N0_HOG0024049), and two CheR-type MCP methyltransferases missing in either CCMP1707 (N0_HOG0021883) or 1850 (N0_HOG0017575). A specific bacterial donor lineage was unable to be pinpointed for the methyltransferase type-11 orthogroup as branch support was low; *Ac. anophagefferens* homologs had best-blast hits to beta-proteobacteria (35% identity) and appeared to be a divergent strain-select duplication of a core genome LGT (N0_HOG0019944) (50% identity, >50% blast hit overlap). Notably, 18 other core genome orthogroups were predicted as methyltransferase type-11 genes, including 13 specific to *Ac. anophagefferens* amongst the pelagophytes that were all predicted to be recent LGTs from a variety of bacteria, suggesting that this type of gene may be acquired by LGT at a high frequency and/or prone to duplication post-acquisition. Although the two CheR-type MCP methyltransferases both appear to be most closely related to gamma-proteobacteria (~34% identity), they had largely non-overlapping top-blast hits (70/500) and align to one another with only 26% identity across 40% of their lengths, suggesting that they could be a result of separate LGTs, or they are divergent paralogs. Due to overlapping blast hits and sequence identity, one of the accessory CheR-type MCP methyltransferases appeared to be a strain-select duplication of a recent LGT within the core genome (N0_HOG0008561) that was retained in all strains except CCMP1707.

Also significantly enriched within GO terms for recent LGTs in the accessory genome were specific hydrolases associated with alpha-glucuronidase and alkaline ceramidase activity. The alpha-glucuronidase (N0_HOG0024163) appeared to be a duplication of a core genome recent LGT specific to *Ac. anophagefferens* (N0_ HOG0011328). Phylogenetic analyses of the alkaline ceramidase showed the *Ac. anophagefferens* homologs embedded within mostly mixed proteobacteria, but that first branched sister to the haptophyte *G. huxleyi*, a pattern suggestive of eukaryote-to-eukaryote LGT (N0_HOG0024019). Two other accessory genome oxidoreductases predicted to be recent prokaryotic LGTs were associated with antioxidant activity – both of which were expressed in all strains and annotated as alkyl hydroperoxide reductases. The genes within these two orthogroups share ~45% amino acid identity across 30% of the gene; however, they had largely non-overlapping blast hits suggesting that they are likely separate LGT events rather than derived from gene duplication. Phylogenetic analyses showed that one peroxiredoxin was likely acquired from delta-proteobacteria (N0_HOG0020729; absent in CCMP1707), while the other had unclear origins, branching with some alpha-proteobacteria, rokubacteria and actinomycetes (N0_HOG0022030; absent in CCMP1707 and 1708).

Other accessory genome LGTs with significantly enriched GO terms included a pectate lyase (discussed below), and an aldehyde lyase specifically annotated as a class-I fructose-bisphosphate aldolase (Fba1) (N0_ HOG0011034) that branched within a clade of Gammaproteobacteria and Actinobacteria (60% identity) and had a close homolog in *P. calceolata*, suggesting it was present in a common ancestor of the Pelagomonadales and secondarily lost in CCMP3368. Transporters in the accessory genome associated with recent LGT includes an arsenate ion transmembrane transporter (N0_HOG0015620; absent in CCMP1850) discussed below and a sulfate permease (SUL1) (N0_HOG0006690) that was likely ancestrally present in the Pelagomonadales and differentially duplicated and lost in select strains. Phylogenetic analyses of the SUL1-containing accessory orthogroup suggests the gene was initially acquired from a Bacteroidetes species (40% identity) and has been transferred between the Pelagomonadales and select dinoflagellate species (*Polarella glacialis* and *Symbiodinium* spp.) via eukaryote-to-eukaryote LGT.

Many of the accessory genome recent LGTs were also associated with binding related GO terms (25, 45%) (Extended Data Figure 7). This included significantly enriched GO terms for phosphopantetheine binding and a mismatched DNA binding related to DNA damage response. One recent accessory genome LGT associated with DNA binding was also associated with hydrolase and transcription factor activity and predicted to encode a homing-endonuclease domain with a NUMOD4 endonuclease DNA-binding motif and AP2/ERF DNA binding domain (N0_HOG0021814; absent in CCMP3368 and 1707). Notably, this was the only predicted NUMOD4 endonuclease within the pelagophytes, a domain that is commonly found in bacteria and viruses[17], but uncommonly identified in eukaryotes outside of metazoa/metamonads (based on annotations in the interpro database). Phylogenetic analyses showed that *Ac. anophagefferens* branched specifically within gamma-proteobacteria (45% identity) and had homologs in a handful of phages and dsDNA viruses (but not AaV).

### *Summary of the functional significance of the accessory genome*

Accessory genes within the *Ac. anophagefferens* pangenome inferred here were predicted to be involved in a wide range of metabolic and cellular processes. Some of the most numerous and significantly enriched were categories related to the metabolism of carbohydrates and lipids or nucleobase-containing and organic nitrogen compounds. Some metabolic process related genes were associated with detoxification of intracellular amines, formaldehyde, and sulfur, and thus could play important roles in differential responses to dealing with potentially toxic metabolites related to heterotrophic metabolism. Others are predicted to be involved in degrading metabolites that are key components of the extracellular matrix and could be related to differential maintenance of the extracellular polymeric substance (EPS) in some strains, which has been speculated to have a link to toxicity in pelagophytes[18]. Two particularly interesting lyase genes were identified within the accessory genome that were acquired in recent LGT events. This included an additional pectate lyase gene in some strains that may result in differing abilities to degrade pectin in plant material in their environment (notably, other pelagophyte species did not encode a pectate lyase gene at all) and a DMSP lyase associated with tolerance of osmotic and oxidative stress[19] and the production of chemical deterrents for grazers and pathogens. Many HAB-forming algae like dinoflagellates and haptophytes are major producers of DMSP[20] – the gene in *Aureococcus* showed high sequence similarity to *Karenia brevis* and other dinoflagellates as well as alpha-proteobacteria, suggesting its presence in *Ac. anophagefferens* is due to a eukaryote-eukaryote LGT of a bacterial gene.

A variety of post-translational modifications were significantly enriched within the accessory genome and several genes involved in transcription regulation and translational initiation, suggesting an important role of accessory genes in fine-tuning metabolic responses to changing environmental conditions via gene expression and post-translational control within individual strains. This was exemplified by the number of predicted transferases encoded in the accessory genome, including several ubiquitin-like protein transferases and strain-specific ubiquitin ligases that have roles in modulating enzyme activity, as well as protein kinases involved in activation/inactivation of enzymes and signal transduction pathways and acyl/methyltransferases that have roles in gene regulation. The abundance of sulfotransferases and glycosyltransferases found to vary between strains could be linked to differences in their extracellular-matrix composition, cell signalling, sulfur metabolism and/or detoxification of sulfur within the cell, and overall glycolipid formation and cell-cell interactions.

Other accessory genes were linked to oxidative stress response (e.g., peroxidases) and DNA repair (e.g., uracil DNA glycosylases, RecQ enzymes, Hiran domain), suggesting that strains may have different abilities to deal with oxidative stress and associated DNA damage. A handful of accessory genes were related to defense related functions and transmembrane transport. This includes differences in abilities amongst strains to transport arsenic from the cell, a heavy metal that is highly abundant in anthropogenically modified environments where it typically forms HABs and can be toxic. Two arsenic transmembrane transporters were identified within the pangenome – a core orthogroup of ancestral origin and a clearly distinct accessory orthogroup acquired by recent LGT that has been subsequently lost in CCMP1850. While this accessory gene does not provide a novel function to strains, it could result in differences in their abilities to detoxify arsenic. Other transporters for a variety of ions, metal ions, nitrogen containing compounds and sulfates were predicted within the accessory genome. This suggests differences between strains in relation to organic nitrogen import exists, which could result in key differences in their reliance on particular nitrogen containing compounds for heterotrophic metabolism. As *Ac. anophagefferens* was previously determined to encode a substantially larger suite of proteins that utilize metal ions as cofactors compared to other algae[21], the presence of a number of metal ion binding proteins in the accessory genome suggests that strains have differing requirements for various metals.

Defense related genes within the accessory genome included genes related to the degradation/inactivation of beta-lactam and chloramphenicol antibiotics. Antibiotics such as beta-lactams are commonly used and present in anthropogenically modified environments, such as those where *Ac. anophagefferens* forms HABs, from wastewater[22]. Both classes of antibiotic degrading enzymes were present in abundance within the core genome, with a large number being identified in *Ac. anophagefferens* specifically or predicted to be the results of recent LGTs (discussed further below). Differential presence between strains within the accessory genome suggests that strains may have different abilities to detoxify these antibiotics and/or to use them as precursors for secondary metabolite biosynthesis. Also related to defense was a MACPF domain containing gene present in select strains that had a patchy distribution amongst eukaryotes and likely rooted in a bacterial origin. MACPF genes within eukaryotes are associated with cytolytic toxins (e.g., ref. [23]), host cell invasion or are known to form pores in the membrane of virus infected cells (e.g., refs. [24,25]). The presence of MACPF was previously noted in the CCMP1984 reference genome[21]). While the exact role of the MACPF gene within select *Ac. anophagefferens* strains is currently unclear and would need to be experimentally investigated, it could have significant impact on bloom dynamics and/or impact co-occurring algae or bivalves within its environment. Variation in its presence between the strains investigated here suggests that it could have an impact on strain-select niche differentiation as the gene could be related to toxin production or controlling levels of virus infected cells within blooming populations. Altogether, these observations hint at diverse roles of accessory genes within the *Ac. anophagefferens* pangenome, many of which could provide functional advantages specific to the environment and conditions it experiences in HABs (e.g., oxidative stress, heavy metals, reliance on heterotrophic metabolism, toxin production) resulting in strain-specific niche adaptations.

### *A unique AaV region in CCMP1850*

Genome-wide sliding window GC content analysis identified a large ~15 kbp region with substantially lower than average GC content uniquely within *Ac. anophagefferens* CCMP1850 (Extended Data Fig. 8), i.e., ~30% like that of AaV[26] compared to ~70% GC like the rest of the genome. The genes and genomic regions surrounding this uniquely low-GC region in 1850 were found to be syntenic with the other four strains, indicating that this was a unique and recent insertion in the 1850 genome; long sequence reads spanned the transitions between the low- and high-GC regions, providing support for its integration. While other core/accessory genes found in the neighboring region were expressed, no RNAseq reads mapped to the low-GC region. Nevertheless, nine ORFs encoding >100 consecutive amino acids were predicted in this region. Six of the predicted ORFs (ORF3-ORF8) had conserved domain hits to a bacterial DUF285 domain (a predicted leucine-rich repeat surface protein often associated with lipoproteins with antigen functions) and had top blast hits to several DUF285 domains in AaV (64-68% identity).

The other three ORFs had no obvious conserved domains but two had top blast hits to a non-DUF285 domain hypothetical protein in AaV (ORF1, 59% identity) or a hypothetical protein in gamma-proteobacteria (ORF2, 58% identity).

Overall, ~43% of the low-GC region in the CCMP1850 genome could be aligned to the AaV genome with nearly 82% identity. Although this region is clearly derived from AaV, it is not obviously colinear with it. Instead, it appears to be limited to regions encoding the DUF285 domain, which are found in two larger regions within the AaV genome, comprising 11.3% of its sequence, which are themselves thought to have been originally acquired by AaV from a bacterium[26]. Other loci in the *Ac. anophagefferens* genome have previously been noted to encode copies of the DUF285 domain[26], but none have been identified within a larger viral-derived region such as that presented here for CCMP1850. Phylogenetic analysis of the DUF285 containing ORFs within the low-GC region and their homologs in the nr database show that all CCMP1850 ORFs branch within AaV sequences with 95 UFBoot2 support. This in turn branches sister to several *Mycoplasma* spp. homologs (89 UFBoot2 support) and then a variety of bacterial species, suggesting that this domain in AaV may have originally been acquired from *Mycoplasma.* Sequences from the different DUF285 encoding ORFs in CCMP1850 do not necessarily branch sister to one another. In fact, they branch highly supported as sister to at least five different AaV DUF285 homologs, suggestive of being acquired as a larger region of the AaV genome rather than one DUF285 gene that has duplicated. One gene flanking this low-GC region (N0_HOG0015793, absent in CCMP1708) is a fatty acid desaturase predicted to be a recent LGT from proteobacteria and appears to have been partially duplicated in CCMP1984 and 3368 after integration; this fatty acid desaturase did not have detectable homologs in viruses, suggesting that it was not also acquired from AaV. Its presence adjacent to this recent integration in CCMP1850 could suggest that this genomic region has been more prone to LGT.

**Functional enrichment of species-specific genes in pelagophytes**

Overall, 73% of *Ac. anophagefferens*-specific, 79% of *P. calceolata*-specific and 90% of *Au. lagunensis*-specific orthogroups showed at least low levels of gene expression (TPM >1). Despite evidence for active transcription, only ~60%, 30% and 45% of the *Ac. anophagefferens*-, *P. calceolata*- and *Au. lagunensis*-specific orthogroups had EggNOG orthogroups or interpro based annotations assigned to them. Ignoring orthogroups that matched to EggNOG orthogroups with unknown functions (~25% in each species), the most abundant KOG/COG terms in *Ac. anophagefferens*-specific orthogroups were related to post-translational modifications ('O'; 575, 10.6%), signal transduction ('T'; 421, 8%), or transport/metabolism of lipids ('I'; 394, 7.3%), carbohydrates ('G'; 304, 5.6%) and amino acids ('E'; 272, 5.1%). Similarly, in *P. calceolata* and *Au. lagunensis,* species-specific orthogroups were most frequently categorized as being involved in post-translational modifications (73/12.9% and 101/10.1%, respectively) and signal transduction (42/7.4% and 68/6.8%, respectively), as well as the transport/metabolism of carbohydrates (41/7.3% and 76/7.6%, respectively) and amino acids (31/5.5% and 56/5.6%, respectively). Additionally, *P. calceolata*-specific orthogroups were abundant in energy production/conversion ('C'; 31/5.5%), while orthogroups specific to *Au. lagunensis* were abundant in replication, recombination, and repair ('L'; 56/5.6%) (Fig. 3.18). Overall, this indicates that there are many differences between the pelagophyte species in terms of their coding capacity associated with post-translational modifications and signal transduction, as well as carbohydrate, lipid and amino acid transport/metabolism – some of which can be attributed to recent prokaryotic LGT (discussed below).

### *Ac. anophagefferens-specific genes*

*Ac. anophagefferens*-specific orthogroups were most significantly enriched in GO terms for biological processes related to transmembrane transport (238 orthogroups), including transport of monoatomic ions (138), metal ions (47), and xenobiotics (3). GO terms related to carbohydrate metabolism (110), including arabinan/arabinose metabolism (5), and protein metabolism (328), including protein modifications (230), were also amongst the most significantly enriched biological processes in orthogroups specific to *Ac. anophagefferens*. Protein modification related terms include a wide variety of significantly enriched post-translational modifications (41) (phosphorylation, glycosylation, methylation, alkylation, hydroxylation, and carboxylation) and peptidyl amino-acid modifications (54) (aspartic acid hydroxylation (7), arginine methylation (12), glutamate carboxylation (2) and peptidyl-prolyl isomerization (24)). Other significantly enriched metabolic processes include those related to the metabolism of amides (31) and fatty acids (16). GO terms that were highly abundant but not significantly enriched include those associated with the metabolism of phosphorous (145), organonitrogen compounds (399), nucleobase-containing compounds (105) and those related generally to biosynthetic processes (104).

In terms of GO terms related to molecular functions, *Ac. anophagefferens*-specific orthogroups were most significantly enriched in orthogroups involved in monoatomic ion channels (128) and passive transmembrane transport (134), specific hydrolases for sulfuric esters (47) and O-glycosyl compounds (83), sulfotransferases (85), carbohydrate binding (87) (mostly monosaccharides (77)) and various categories of oxidoreductases (334). While general transmembrane transport (226) was significantly enriched overall, transporters associated with specific metabolites/molecules were not. Some of the most numerous transporters in orthogroups specific to *Ac. anophagefferens* include those for chloride ions (13), lipids (5), xenobiotics (4), ammonium (4), carbohydrate derivatives (3), and metal ions (56) (e.g., calcium (7), magnesium (3), sodium (3) and potassium (31)). Over 60% of *Ac. anophagefferens*-specific orthogroups with a predicted GO term were related to binding (1724), including many significantly enriched specific binding related terms. Ion binding (665) was particularly highly abundant, with ~60% specifically associated with metal ion binding (398); this included significantly enriched terms for binding of transition metal ions (173) and iron ions (89), and highly abundant terns for binding of calcium (166) and zinc ions (73). Protein binding related terms were numerous and significantly enriched (928), with ~1/3 associated with catalytic activities acting on proteins and a wide variety of other molecular functions. Carbohydrate binding (87), and specifically monosaccharide binding (77) GO terms were both significantly enriched, as was amide (27) and modified amino acid binding overall (26). Also notable was the significant enrichment of orthogroups involved in vitamin binding (111), including those specifically involved in L-ascorbic acid binding (75), which has roles as a cofactor for several enzymes and in protection from oxidative stress via neutralization of ROS.

Nearly half of *Ac. anophagefferens*-specific orthogroups with predicted molecular function GO terms were associated with a catalytic activity (1408). Oxidoreductases (334) were found to be significantly enriched and numerous in general. This included a variety of different oxidoreductase classes, including those acting on paired donors with incorporation/reduction of molecular oxygen (91). Hydrolases (412) were even more abundant, including several significantly enriched subtypes that were mainly sulfate ester hydrolases (47) and hydrolases that act on O-glycosyl compounds (93), including more specifically enriched enzyme classes such as alpha-glucuronidases (4), alpha-L-arabinofuranosidases (5) and beta-N-acetyl-hexosaminidases (7). Within predicted peptidases (95), carboxypeptidases (15) were significantly enriched and largely included serine-type (11), cysteine-type (32) and metallopeptidases (18). The most abundant enzymes within *Ac. anophagefferens*-specific orthogroups were transferases (517). Significant enriched classes of transferases include acyltransferases (138), sulfotransferases (85) (e.g., aryl sulfotransferases (20)) and arginine-specific methyltransferases (12). Some more numerous, but not significantly enriched, transferases identified were phospho- (127), methyl- (91), glycosyl- (53), and ubiquitin-transferases (48).

### *P. calceolata-specific genes*

*P. calceolata*-specific orthogroups were largely associated with GO terms for biological processes related to metabolism (84), transport (25) or biological regulation (14). Compared to pelagophytes overall, *P. calceolata*-specific orthogroups were most significantly enriched in protein metabolism (45) and related protein modifications (36), organonitrogen compound metabolism (60), regulation of metabolic processes (14), inorganic ion transmembrane transport (8) and guanosine tetraphosphate metabolism (2). Orthogroups associated with protein modifications included a number of more specific modifications that were significantly enriched such as protein phosphorylation (15), ubiquitination (7), peptidyl-proline isomerization (4), peptidyl-histidine modification to diphthamide (1), C-terminal amino acid methylation (1) and peptidyl-amino acid modifications overall (8). Several other GO terms related to metabolic processes were significantly enriched, such as inositol catabolism (1) and those associated with nucleobase-containing small molecules (11); these including purine metabolism (7), and more specifically IMP (2) and guanosine tetraphosphate (2) metabolism, guanine catabolism (1), and queuosine biosynthesis (1). While not significantly enriched, a number of *P. calceolata*-specific orthogroups were related to metabolism of phosphate-containing compounds (24) or carbohydrate derivates (8).

GO terms associated with regulation of metabolic processes mostly included orthogroups involved in regulation of DNA-templated transcription (11) and were significantly enriched in the regulation of phenylpropanoid metabolism (1) and the negative regulation of cytokinin-activated signaling pathways (1). Some *P. calceolata*-specific orthogroups were predicted to be involved in response to stimuli (6), including significantly enriched GO terms for response to high light intensity (1) and to DNA damage (3). Several specific transport related GO terms were found to be significantly enriched. These included those involved in the transport of inorganic ions (6) such as phosphate (2) and ammonium (2), as well as transporters for sulfur-containing compounds (2), including biotin (1). Additionally, one significantly enriched orthogroup unique to *P. calceolata* was associated with peroxisome fission and specifically annotated as peroxisomal biogenesis factor 11 (PEX11).

In terms of GO terms related to molecular functions, *P. calceolata*-specific orthogroups were most significantly enriched in orthogroups involved in enzyme inhibitor activity (3) (mostly serine-type endopeptidase inhibitors (2)), zinc ion binding (18), amino-acyltransferase activity (11) and formyltetrahydrofolate deformylase activity (2). In general, most orthogroups were associated with binding (223), catalytic activity (150) or transmembrane transport (24). While not significantly enriched, some of the more abundant GO terms involved in binding were specifically related to binding proteins (118) or ions (90), including a variety of metals (53), nucleotides (36), and carbohydrate derivatives (32). Transmembrane transport (24) related terms include both passive transport (10) and significantly enriched secondary active transport (5) mechanisms. More specific enriched transporters include those for inorganic anions (4), phosphate (2) and biotin (1). Around 40% of *P. calceolata*-specific orthogroups were associated with catalytic activity. The most numerous class of enzymes identified in *P. calceolata*-specific orthogroups were transferases (64). These include significantly enriched GO terms specific to aminoacyl transferases (11), thymidine kinases (1) and other protein kinases (14), carboxyl-O-methyltransferases (2) and other methyltransferases (11), and ubiquitin-protein transferases (10). A number of orthogroups were also associated with hydrolase activity (31), including significantly enriched serine-type endopeptidases (2) and formyltetrahydrofolate deformylases (2). Some specific classes of oxidoreductases (33) were also significantly enriched, mainly a preQ1 synthase associated with queuosine biosynthesis. Other specific classes of enzymes that were significantly enriched included a methylglyoxal synthase related to lyase activity (3), and a mannose-6-phosphate isomerase related to isomerases (10).

### *Au. lagunensis-specific genes*

Orthogroups specific to *Au. lagunensis* were mostly associated with GO terms for biological processes related to metabolic processes (194), transport (55) or biological regulation

(41). The most significantly enriched biological processes were related to DNA metabolism (28) and integration (16), signal transduction (23) (including intracellular (14) and phosphorelay signal transduction (8) as well as cell surface receptor signaling pathways (5)), biological regulation (41) and sulfate assimilation by phosphoadenylyl-sulfate reductases (2). Although not significantly enriched, some abundant terms included those related to general metabolic processes for organonitrogen compounds (102), proteins (83) and related protein modifications (57), nucleobase-containing (49) and phosphate-containing (44) compounds. Transport related terms (55) were significantly enriched in chloride (3), ammonium (3), chromate (1) and lactate (1) transport. Other abundant transport related GO terms included those for nitrogen compounds (11). Within cellular processes, telomere maintenance and organization (3) and intracellular monoatomic cation homeostasis (2) GO terms were significantly enriched.

Protein modification terms were largely associated with phosphorylation (28), peptidyl-amino acid modifications specific for proline (4), aspartate (2) or histidine (10), protein ubiquitination (9) and deubiquitination (2), and significantly enriched in glycosylation (10). Some specific processes related to amino acid metabolism (6) were significantly enriched, mainly asparagine biosynthesis (2), D-amino acid metabolism (1) and proline catabolism to glutamate (1). Several processes related to carbohydrate (21) and carbohydrate derivative (7) metabolism were also significantly enriched; these include polysaccharide metabolism (5), beta-glucan metabolism (3), 1-3-beta-D-glucan biosynthesis (2), peptidoglycan catabolism (1) and glycosaminoglycan catabolism (1), as well as substituted mannose (1) and xylulose (1) metabolic processes. Other significantly enriched metabolic processes included ectoine biosynthesis (1). Also significantly enriched in species-specific orthogroups in *Au. lagunensis* were processes related to stress response (18), which included a number of orthogroups predicted to be involved in a variety of DNA repair processes (12) or were peroxidases related to oxidative stress (3).

The most significantly enriched GO terms associated with molecular functions in *Au. lagunensis*-specific orthogroups were sialytransferases (3), phosphoadenylyl-sulfate reductases (2), malic dehydrogenases (2), phosphatidylinositol binding proteins (8), GDP-mannose transmembrane transporters (2), 1,3- beta-D-glucan synthases (2) and peptidoglycan muralytic lysozymes (2). Most orthogroups were associated with binding (360), catalytic activity (310) or transmembrane transport (45). Significantly enriched binding related GO terms included both phosphatidylinositol (8) and sequence specific DNA binding (8). Other binding related GO terms amongst the most abundant in *Au. lagunensis*-specific orthogroups include those related to binding proteins (161), ions (142), nucleotides (85) (largely specifically purines (77)) and carbohydrate derivatives (70). A variety of transmembrane transport related GO terms were significantly enriched, including potassium (3), ammonium (3), GDP-mannose (2), lactate (1), and chromate (1) transporters as well as chloride channel activity (5).

As in the other pelagophyte species-specific orthogroups, a wide variety of functions related to catalytic activity was identified in *Au. lagunensis*. The most numerous were transferases (106), which included glycosyltransferases (24), phosphotransferases (30), acyltransferases (29), sulfotransferases (14) and ubiquitin-protein transferases (12). Many specific enzyme activities were significantly enriched within subclasses of transferases. These include sialytransferases (3), 1,3-beta-D-glucan synthases (2) and other glycosyltransferases (22); a formate c-acetyltransferase and a diaminobutyrate acetyltransferase; protein histidine kinases (2), phosphorelay kinases (2) and a D-xylulokinase (1); and alkylglycerone-phosphate synthase activity (1). While many *Au. lagunensis*-specific orthogroups were predicted to be hydrolases (101), no specific subclasses were significantly enriched. Some of the most abundant hydrolases were related to those acting on ester bonds (20), acid anhydrides (17) or O-glycosyl compounds (13), or were predicted to be peptidases (30). Orthogroups associated with oxidoreductase activity (59) were significantly enriched in specific GO terms for malate dehydrogenase (2), 1-pyrroline-5-carboxylate dehydrogenase (1), ferredoxin hydrogenase (1) and phosphoadenylyl-sulfate reductase activities (2). A handful of lyases (7) and isomerases (9) were additionally identified, none of which were significantly enriched.

### *Overall trends in species-specific genes*

The functional annotations associated with species-specific genes suggests that these pelagophyte species have diverse metabolic capabilities. A number of transmembrane transporters were specific to individual pelagophyte species, suggesting that they have different abilities in terms of what metabolites they uptake and use for metabolic processes. *Ac. anophagefferens*-specific orthogroups were most significantly enriched in transmembrane transport, including a number of species-specific metal ion transporters, that together with a large number of species-specific metal-binding proteins reflects its unique requirement for metal cofactor use in a variety of catalytic enzymes[21]. Also significantly enriched were *Ac.*

*anophagefferens*-specific xenobiotic transmembrane transporters likely involved in the detoxification of chemicals such as drugs, pesticides or pollutants present in anthropogenically modified environments and can be directly related to providing a competitive advantage in the environment for which they for HABs. *P. calceolata* on the other hand was significantly enriched in transmembrane transporters including those for sulfur compounds and a bacterial-type biotin transporter, which likely represents a novel way for *P. calceolata* to uptake biotin from its environment more effectively. *Au. lagunensis*-specific genes were significantly enriched in GDP-mannose transmembrane transporters and specific transporters for chloride, ammonium and chromate, the latter of which could be directly related to detoxification of chromates present in the environment from industrial pollution. Species-specific nitrogen compound transporters were identified in *Ac. anophagefferens* and *Au. lagunensis* (3-4 unique transporters for ammonium/urea and 1 unique gene classified as a ureide permease in both species), as were a large number of species-specific genes related to organic nitrogen compound metabolism (399 in *Ac. anophagefferens* and 102 in *Au. lagunensis*). Together this could reflect substantial differences in uptake and utilization of different organic nitrogen compounds for heterotrophic metabolism.

Outside of transmembrane transport associated genes, species-specific orthogroups were largely associated with catalytic activities that are likely linked to their unique metabolic capacities. In *Ac. anophagefferens,* carbohydrate, proteins and fatty acid metabolic processes (including many linked to polyketide synthases) were significantly enriched, including a large number of enzymes predicted to be involved in post-translational modifications and peptidyl amino-acid modifications. Protein modification related GO terms were also significantly enriched in *P. calceolata*-specific orthogroups, suggesting that these species have different suites of post-translational modifications used to regulate protein function and adapt to environmental conditions. *P. calceolata*-specific orthogroups were significantly enriched in some specific metabolic processes related to purine metabolism, including a gene involved in the biosynthesis of queuosine nucleosides that suggests it is uniquely able to synthesize these nucleotides *de novo* (discussed further below).

Also related to purine metabolism were two species-specific formyltetrahydrofolate deformylases (which are genes related to IMP biosynthesis) and a guanine deaminase involved in utilizing guanine as a nitrogen source via the production of xanthine and ammonia; both of these appear to be functionally redundant with a number of genes found in common with at least *Ac. anophagefferens,* suggesting that rather than providing a unique function to *P. calceolata* they could result in an increased capacity for both purine biosynthesis and nitrogen metabolism via the degradation of guanine. Completely unique to *P. calceolata* was a methylglyoxal synthase that has been linked to metabolic flexibility under stress conditions (mainly phosphate starvation) in bacteria[27]; this gene likely has a more ancient bacterial origin from cyanobacteria as it was also identified in some select diatoms and a green alga. In *Au. lagunensis,* species-specific genes were significantly enriched in signal transduction, suggesting this species has unique mechanisms to adapt and response to their environment and/or could be related to interactions with other organisms and specific niche specializations. DNA integration terms significantly enriched within *Au. lagunensis*-specific genes appeared to be largely associated with expansions of transposons within the species. Some other significantly enriched species-specific genes within *Au. lagunensis* included a phosphoadenylyl-sulfate reductases involved in sulfate assimilation crucial for further synthesis of sulfur-containing compounds, and a variety of carbohydrate metabolism terms related to beta-glucan metabolism, peptidoglycan and glycosaminoglycan catabolism and xylulose metabolic processes, suggesting it has a range of unique carbohydrate metabolism capabilities.

Pelagophyte species encoded a large number of orthogroups involved in dealing with oxidative stress and detoxification of ROS and other metabolites. Overall, *Ac. anophagefferens* had ~1.5X more orthogroups associated with the 'antioxidant activity' GO term (52) than *P. calceolata* (34) or *Au. lagunensis* (32). This included an increase in species-specific orthogroups annotated as heme peroxidases, hydroperoxide reductases, glutaredoxins, thioredoxins, and a variety of superoxide dismutase enzymes (Ni, Cu/Zn), suggesting that it has a greater capacity to deal with oxidative stress compared to other pelagophyte species. The glutathione S-transferase family of proteins, which can also play a role in detoxifying ROS, xenobiotics and other metabolic byproducts, were also highly expanded within *Ac. anophagefferens,* who encoded nearly double the number of orthogroups with these predicted domains.

**The functional impact of LGT in pelagophytes**

The vast majority of candidate recent LGTs within pelagophytes were found to have transcriptome-based evidence for expression (>80%) and, by, extension, functionality. Predicted

LGTs across pelagophytes were associated with a wide variety of COG functional categories, including most frequently carbohydrate, secondary metabolite, amino acid or lipid transport and metabolism, as well as post-translational modifications (Extended Data Fig. 9B). In *Ac. anophagefferens*, species-specific LGTs made up 40-50% of LGTs identified in most COG categories related to transport and metabolism as well as post-translational modification and signal transduction, suggesting that recent LGTs have a large role in *Ac. anophagefferens'* unique metabolic capabilities and ability to regulate protein activity in response to changing environmental conditions. Species-specific LGTs within the other species contributed proportionality less to most COG categories, except for DNA replication and repair. Within *Ac. anophagefferens*, most recent LGTs were present within the core genome (~87%) and the overall contribution of LGT to a given strain was highly similar (3.9 +/- 0.1% of orthogroups). There was some evidence for LGT continuously contributing to the evolution of individual *Ac. anophagefferens* strains; the majority of recent LGTs, though, were presumably present in the genome of at least their common ancestor and differentially retained and/or duplicated throughout their evolution, suggesting they played a role in helping the ancestor of these *Ac. anophagefferens* strains come to dominance. While being cautious to not over-interpret the data, the increased proportion of recent LGT-derived genes within both HAB forming pelagophyte species is worth noting. This might suggest the acquisition of new bacterial genes via species-specific LGTs and differential retention of LGTs ancestral to the Pelagophyceae are at least in part related to their association with specific HAB environments and have contributed to their ecological success.

Recent prokaryotic/viral LGTs identified unique to *Ac. anophagefferens* amongst pelagophytes (i.e., *Ac. anophagefferens*-specific orthogroups) were most significantly enriched in GO terms for biological processes related to protein metabolism (29) and related protein modifications (14), peptidyl-amino acid modifications (6) and proteolysis (13), carbohydrate metabolism (24) and metabolic processes related to organic substances in general (66) (Extended Data Fig. 10A). Over half of *Ac. anophagefferens*-specific LGTs associated with primary metabolism were related to metabolic processes of organonitrogen compounds (38) (e.g., putrescine (1) and tetrapyrrole (1) biosynthesis). While most other biological processes were not significantly enriched, many were abundant and/or potentially interesting in relation to *Ac. anophagefferens* HAB dynamics such as those connected to transmembrane transport (5) and response to either DNA damage (3) or oxidative stress (2).Molecular function related GO terms predicted for *Ac. anophagefferens*-specific LGTs were largely associated with catalytic activities (186/239, 78%) (Fig. 3B) and included a wide variety of hydrolases (56), oxidoreductases (47), and transferases (58), as well as a handful of lyases (3), isomerases (4) and ligases (2). Also highly abundant were GO terms related to binding (97); these included significantly enriched terms associated with binding metal ions (29) (e.g., transition metals (15)), vitamins (18) and carbohydrates (13). While not enriched, many orthogroups were associated with binding of ions (46), proteins (38) or nucleotides (23) in general.

*Au. lagunensis*-specific recent LGTs were mostly associated with biological process GO terms for metabolic processes (28 orthogroups), including significantly enriched in those related to glycosyl compounds (2), xylulose (1) and glutamine (2) biosynthesis, and DNA integration (2) (Extended Data Fig. 10B). Other LGTs were predicted with GO terms for transport (5) and biological regulation (2). Molecular function-related GO terms associated with *Au. lagunensis*-specific LGTs were most significantly enriched for catalytic activities (40), including significantly enriched in specific genes for xylulokinase, 1-pyrroline-5-carboxylate dehydrogenase, DNA- protein cysteine S-methyltransferase, 2-phosphosulfolactate phosphatase and lysozyme activities. (Fig. 3C). GO terms for transmembrane transport (4) and binding (23) (e.g., significantly enriched cellulose binding) were also predicted.

Overall, *P. calceolata*-specific orthogroups attributed to recent LGT were associated with biological processes related to either metabolism (6 orthogroups) or transport (1) (Extended Data Fig. 10C), and molecular functions related to catalytic activities (19), transmembrane transport (1) or binding (9) (Extended Data Fig. 10D). The most significantly enriched GO terms for biological processes were related to genes involved in biotin transport, queuosine biosynthesis, inositol catabolism, guanine tetraphosphate metabolism and C-terminal protein amino acid methylation. *P. calceolata-* specific recent LGTs were significantly enriched in molecular function associated GO terms for a number of catalytic activities, which included a variety of oxidoreductases (7), transferases (7), and some specific hydrolases (3) associated with peptidase activity. The most significantly enriched molecular functions were related to preQ1 synthase, inositol oxygenase, lysine N- methyltransferase, serine-type endopeptidase inhibitor and biotin transport activities.

**Species-specific LGTs contribute to differential metabolic flexibility**

Due to the ecological and economic impacts of pelagophyte HABs, the characteristics of their blooms have been extensively studied to understand factors contributing to their establishment, proliferation and decline (e.g., see ref. [28] and references therein; [18,29]). Both pelagophyte species known to form HABs (*Ac. anophagefferens* and *Au. lagunensis*) thrive under low concentrations of dissolved inorganic matter and high concentrations of dissolved organic matter (particularly nitrogen and carbon) and are able to outcompete co-occurring species for nutrients/access unique nutrient sources during intense shading throughout blooms that requires them to heavily rely on heterotrophic metabolism (e.g., see refs. [16,21]). *Ac. anophagefferens* blooms in particular are notably preceded by diatom blooms that, together with agricultural run-off and wastewater, create a nutrient loading scenario whereby abundant organic matter creates optimal conditions for *Ac. anophagefferens* blooms given their gene complement[21]. Dealing with nutrient related stress resulting from limited availability of various compounds is closely linked to being able to fine-tune their metabolism in response to fluctuating nutrient and light conditions and encoding unique and/or duplicated genes that increases their metabolic flexibility – numerous recent LGTs within these pelagophytes were frequently assigned to KOG/COG functional categories related to carbohydrate, secondary metabolite, amino acid or lipid transport and metabolism as well as with post-translational modifications and thus are associated with such metabolic flexibility. Some recent LGTs encode apparently novel functions (i.e., those new to the organism(s)), while others are predicted to encode domains similar to other ancestral genes, thus potentially increasing metabolic flexibility and/or efficiency under specific conditions.

### *LGTs related the metabolism of nitrogen-containing compounds*

In both *Ac. anophagefferens* and *Au. lagunensis*, GO terms associate with LGTs were significantly enriched in organonitrogen-containing compounds, suggesting that recently transferred genes has contributed to differences in nitrogen metabolism between these HAB-forming species. A handful of species-specific LGTs in *Au. lagunensis* are related to amino-acid degradation, including the catabolism of proline and aromatic amino acids (e.g., under oxygen limiting conditions by an alpha-proteobacterial derived indolepyruvate ferredoxin oxidoreductase (SINGLETON_2810) involved in tryptophan and other aromatic amino acid catabolism) that could contribute to broader cellular energy processes and provide alternative carbon and nitrogen sources. Also specific to *A. lagunensis* was a highly expressed (165 TPM) glutamine synthase type III gene (N0_HOG000039112) recently acquired from either alpha/beta-proteobacteria (60% identity) that was distinct from another glutamine synthase type III enzyme found in all pelagophytes (<30% identity); its presence could allow for increased synthesis of glutamine from ammonia in *Au. lagunensis*, supporting a variety of other biosynthetic processes for which glutamine acts as a nitrogen source. On the other hand, *Ac. anophagefferens*-specific LGTs included a bacterial L-aspartate dehydrogenase (N0_HOG0020033) likely acquired from alphaproteobacteria (~60% identity) and involving eukaryote-to-eukaryote gene transfer to/from haptophytes. As no other aspartate dehydrogenase was annotated within the pelagophytes and are relatively rare in eukaryotes, its presence suggests that this LGT provides an alternative and unique way for *Ac. anophagefferens* to metabolize aspartate compared to most other algae, contributing to its metabolic flexibility.

A unique complement of LGT-derived serine-type and metallo-peptidases in *Ac. anophagefferens* (8 orthogroups; significantly enriched) could be involved in a higher capacity for protein turnover in response to nutrient availability and/or related to nitrogen scavenging from extracellular protein degradation. Previous experimental work has suggested that *Ac. anophagefferens* can use protein present in its environment as a source of nitrogen via hydrolysis of peptides at its cell surface[30]. As a number of these peptidases acquired by recent LGT were predicted to function extracellularly (e.g., two subtilisin-related S8 peptidases, an S9 peptidase an M14 type A carboxypeptidase), they could provide at least part of the enzymatic explanation for this observation. A variety of different classes of peptidases without signal peptides were identified in each set of species-specific LGTs, suggesting they each have varying abilities to cleave distinct peptide bonds (S8/S9/M14/M16/M17 in *Ac. anophagefferens*, C15 in *P. calceolata* and S49/M41 in *Au. lagunensis*), contributing to proteolysis and regulation of protein activity within the cell.

The highly expressed C15 family peptidase identified to a LGT present only in *P. calceolata* was annotated as a bacterial type pyroglutamyl peptidase I) (SINGLETON_1063). Phylogenetic analyses of this gene showed the *P. calceolata* homolog branching within a highly supported clade including other *P. calceolata* strains, another closely related Pelagomonadales species (*Pelagococcus subviridis*) and a green alga (*Tetraselmis astigmatica*) prior to being embedded within bacterial sequences mostly derived from proteobacteria (52% identity) – a topology suggestive of eukaryote-to-eukaryote LGT of an initially acquired bacterial gene. This specific family of peptidases is involved in removing pyroglutamic acid (a proteogenic amino acid derived from cyclization of glutamine) at N-terminal positions of proteins; having an additional version may allow for greater efficiency in protein degradation/turnover and a more efficient response to changes in its environment. In *P. calceolata*-specific LGTs, the GO term for 'serine-type endopeptidase inhibitor activity' was significantly enriched.

One particularly interesting gene originating from LGT was a moderately expressed (4-11 TPM) D-alanine-D-alanine ligase (DdIA) in *Ac. anophagefferens* (N0_HOG0018385) likely acquired from deltaproteobacteria (37% identity) directly or indirectly via the green alga *Cymbomonas tetramitiformis* (48% identity). DdIA genes are generally thought to be uncommon in eukaryotes; they are typically associated with bacteria, where they form D-alanine dipeptides, key precursors for peptidoglycan synthesis. Due to its presence in bacterial cell walls, D-alanine is amongst the most abundant D-amino acids in the ocean and components of dissolved organic nitrogen[31,32]. Plastids that retain a peptidoglycan layer are generally rare in algae and have been observed in glaucophytes (e.g., ref. [33]) and some land plants (e.g., ref. [34]); at least some of the protein-coding genes responsible have been identified in their nuclear genomes[35,36] as well as the genomes of some green algae like *C. tetramitiformis* (e.g., refs. [37,38]). The only gene present specifically related to peptidoglycan synthesis in *Ac. anophagefferens* was DdIA, suggesting that instead of resulting in the synthesis of a peptidoglycan like layer, the enzyme probably functions in an alternative role. D-alanine and D- alanine dipeptides in some bacteria have been associated with cell signalling related to biofilm dissolution as well as modification of cell wall surface proteins related to antibiotic resistance[39]; D-alanine dipeptides could play a similar role and be related to regulation/modifications of the EPS layer of *Ac. anophagefferens* – however, the exact function is unclear.

A number of transporters that were specifically amino acid permeases were encoded in the *Ac. anophagefferens* genome, suggesting it is likely able to transport D-alanine into its cell from the environment. A gene for alanine racemase that converts the L and D enantiomers was also found within the core genome of *Ac. anophagefferens* that had homologs in *P. calceolata* and diatoms, ultimately appearing to be derived from a more ancient bacterial LGT within the Diatomista. A previous study determined that D-alanine in diatoms was mainly derived from its own synthesis via this alanine racemase rather than from bacteria in its environment[40]. Whether D-alanine within *Ac. anophagefferens* is largely derived from its own synthesis or is partly acquired from bacterial degradation in its environment remains to be determined, as does the precise function and incorporation of D-alanine within the cell. Interestingly, D-alanine is known to be highly abundant in certain bivalves, including species of hard clams that have been impacted by *Ac. anophagefferens* blooms, where they accumulate D- alanine and rely heavily on it as an osmoprotectant during environmental stress[41,42]. Whether the use of D-alanine in both *Ac. anophagefferens* and bivalves is a coincidence or potentially related to its impact on the species is unknown. Although speculative, by forming D-alanine dipeptides, *Ac. anophagefferens* could make both D and L- alanine less bioavailable to bivalves, limiting their ability to regulate their internal salt balance – a process that is critical for its survival in this environment.

Some non-species specific LGTs within the pelagophytes were also related to metabolic flexibility. One example of an LGT shared by all pelagophyte species was an AroE enzyme that functions in the shikimate pathway (N0_HOG00223711 and N0_HOG0016006; recently acquired from a proteobacterial species (34-39% identity)), which links carbohydrate and amino acid metabolism and is associated with the production of numerous secondary metabolites (e.g., reviewed in ref. [43]). The other six enzymes within the shikimate pathway (DAPH synthase II and AroB/D/K/A/C) were found to be present in all pelagophytes and are all closely related to homologs in other stramenopile algae, branching in monophyletic clades that were sister to land plants/green algae and/or bacteria in their respective phylogenetic trees. AroE genes in diatoms are encoded as an AroD/E fusion gene closely affiliated with fungi and plants[44]. All pelagophyte species were found to encode a similar fusion. The additional AroE gene within pelagophytes had no close homologs in other algae, clearly branching within proteobacteria. Some genes within the shikimate pathway in fungi, plants, oomycetes and dinoflagellates have been attributed to both prokaryote-to-eukaryote and eukaryote-to-eukaryote LGT events and to EGT[44,45]. The presence of a second copy of the AroE gene in pelagophytes acquired by a recent LGT is intriguing and could facilitate greater efficiency of the shikimate pathway within pelagophytes, an important process for generating aromatic amino acids and potentially other secondary metabolites like phenolics.

### *LGTs related to the degradation of plant-tissue*

Some recent LGTs specific to either of the HAB-forming pelagophyte species were predicted to be secreted from the cell and related to the degradation of plant-based material. *Au. lagunensis* encoded a whole suite of genes that could allow for the unique utilization of xylose directly as an organic carbon source via the XI pathway[46]. Within this potential pathway, a non-LGT derived xylose isomerase (N0_HOG0000737) would first convert xylose to xylulose, which in turn would be phosphorylated by an LGT-derived xylulokinase (XylB) (N0_HOG0023691), producing xylulose-5-P that can enter the pentose phosphate pathway. XylB enzymes are not commonly identified in eukaryotes outside of green algae; instead, they are typically found in bacteria where they are involved in xylose metabolism. Phylogenetic analyses of the XylB gene in *Au. lagunensis* supports an initial LGT from a *Planctomycetes* bacterium (46%) into chlorophyte algae followed by a eukaryote-eukaryote LGT into *Au. lagunensis* (Fig. 5C).

Two other enzymes recently acquired by LGT in *Au. lagunensis* were predicted to contribute to xylose generation via plant tissue degradation. This includes a feruloyl esterase B/C/D (SINGLETON_3123) and xylosidase (YicI; SINGLETON_4196) that together function in degrading plant tissue by removing feruloyl esters linked to xylan molecules, making them accessible to xylosidases that can then generate xylose from xylan[47,48], a product that could feed directly into XI pathway, contributing to utilization of xylose directly as a carbon source. Phylogenetic analysis of the YicI gene showed the *Au. lagunensis* homolog branching in a maximally supported monophyletic group with some dinoflagellates and a single haptophyte species that then branched sister to a clade of mostly CFB-group bacteria. While the overall phylogeny is suggestive of eukaryote-to-eukaryote LGT of an initially acquired bacterial gene into eukaryotes, the specific directionality of transfer between these algae is unclear. Phylogenetic analysis of the feruloyl esterase also was indicative of secondary eukaryote-to-eukaryote transfer of an originally acquired bacterial gene between *Symbiodinium* spp., *E. huxleyi* and *Au. lagunensis*.

Notably, XylB and xylosidase genes are occasionally genomically linked in operons in bacterial species such as firmicutes[49]. While there was some overlap in the closest-related bacterial species to both genes identified in *Au. lagunensis*, the genes were located on different contigs and ultimately branched with different subsets of bacterial species suggesting that while it remains a possibility, the two genes were likely not acquired together in a single unit. All of these genes were predicted to be secreted, supporting an extracellular function in plant tissue degradation specifically linked to xylans present in the hemicellulose component of plant cell walls and other components of plant tissue. While a xylose-specific transporter (as found in some bacterial species) was not identified, xylose uptake into *Au. lagunensis* could occur by one of the many non-specific MFS transporters or ABC transporters encoded within the genome as has been observed in some bacteria and archaeal species[46]. Two other recent LGTs specific to *Au. lagunensis* were annotated as cellulases and could contribute to an enhanced capability for cellulose degradation; these were likely acquired from either gamma-proteobacteria (47% identity) (N0_HOG0022260) or CFB-group bacteria (52% identity) (SINGLETON_4245).

*Ac. anophagefferens*, on the other hand, has many genes that suggest it is capable of degrading pectin, glucans/starches and components of hemicellulose and could allow it to largely rely on plant material as an organic carbon source. A large complement of simple and complex carbohydrate-degrading enzymes has been previously identified within *Ac. anophagefferens*[16,21]. Those identified here included a number of recent LGTs predicted to be secreted from the cell, supporting their function extracellularly in plant cell wall degradation. This included a polygalacturonase (Pgu1) with a pectin lyase fold (N0_HOG0008316) and a pectate lyase (PelB) that together convert pectin present in plant cell walls into pectate and further degrade it. Phylogenetic analyses of the Pgu1 gene in *Ac. anophagefferens* found it to branch with a large group of mostly high-GC gram-positive actinobacteria (32% identity) and the haptophyte *Chrysochromulina ericina*, suggestive of eukaryote-to-eukaryote LGT of an initially acquired bacterial gene (Fig. 5E). While a homolog to Pgu1 was not identified in *P. calceolata*, one was identified in the closely related species *Pelagococcus subviridis* that branched with *Ac. anophagefferens* sequences in the phylogeny; this suggests that this LGT occurred more ancestrally within the Pelagomonadales and has been subsequently lost in *P. calceolata*. Several other orthogroups were identified across pelagophytes with a pectin lyase fold, including two other recent LGTs found in all pelagophytes that were divergent from both the *Ac. anophagefferens*-specific LGT discussed above and from one another (25% identity and only 20-25% gene coverage). While it remains possible that these pectin lyase-containing orthogroups are the result of substantially diverged duplications of a single LGT, phylogenetic evidence strongly suggests these were likely acquired as three separate gene transfers (data not shown).

Two PelB genes were identified specifically in *Au. anophagefferens* – one present in the core genome not obviously derived from LGT (N0_HOG0019014) and a second present in the accessory genome (N0_HOG0020712) acquired recently from within actinobacteria and specifically branched sister to a clade of *Nocardiopsis* spp. (42% identity). The accessory PelB gene was highly expressed (>150 TPM) and predicted to encode a signal peptide, suggesting its protein product functions extracellularly. An additional pectate lyase gene in some strains may result in differing abilities to degrade pectin in plant material in their environment. Notably, neither orthogroup was identified in other pelagophytes, nor were other specific pectate lyase domains, suggesting a unique role in *Ac. anophagefferens.* Pectate lyases, and particularly PelB genes, have been identified in a variety of plant-pathogenic and other bacteria, fungi[50–52] and in oomycetes (e.g., refs. [44,53]) where they are typically implicated in strategies for plant invasion and degradation. Some fungal homologs appeared more distantly related to the gene identified in *Ac. anophagefferens,* but a separate origin was strongly supported by phylogenetic analyses. While it is more likely these pectin metabolism genes are predominantly working to degrade decaying plant material in *Ac. anophagefferens*' environment due to plant defenses for microbial grazing that would have to be overcome, it is possible that they are acting to some extent on live seagrass, contributing to their demise.

Other than pectin/pectate degradation, an alpha-L- fucosidase (N0_HOG0011319) acquired via LGT uniquely present in *Ac. anophagefferens* predicted to also be secreted suggests the organism can access fucose present in glycans, glycoproteins and glycolipids of plant cell walls. Phylogenetic analyses resolved the *Ac. anophagefferens* homologs as directly sister to a handful of bacteria, albeit with low support. While the origin of this gene appears ultimately to be from bacteria, the patchy presence of some dinoflagellates and haptophytes nearby suggests that eukaryote-to-eukaryote LGT could be involved. Additionally present was a predicted YihQ-like alpha-glucosidase in *Ac. anophagefferens* (N0_HOG0020068) that branched directly with spirochaetes with high support, with some other eukaryote homologs from Viridiplantae present nearby. YihQ-like alpha-glucosidases are not commonly found in eukaryotes outside of Viridiplantae; as they are involved in the metabolism of complex carbohydrates via the breakdown of alpha-glucosidic bonds in polysaccharides/complex sugars, their presence likely expands *Ac. anophagefferens*' ability to rely on different carbohydrate sources for heterotrophic metabolism during HABs. Two orthogroups derived from a single LGT from Actinobacteria (39% identity) were predicted with alpha-glucuronidase activity and associated with xylan catabolism (N0_HOG0024163 and N0_HOG0011328) (AguA2) (Fig. 5D) were detected in *Ac.*

*anophagefferens* specifically that could be related to degradation of the hemicellulose component of plant cell walls. Other pelagophytes did not encode a predicted alpha-glucuronidase, nor were any annotated outside of opisthokonts in the interpro database, suggesting that this enzyme provides *Ac. anophagefferens* with a unique functional capability to break down plant material as a part of their nutrient acquisition strategies.

Both *Ac. anophagefferens* and *Au. lagunensis* blooms are known to substantially impact sea grass beds, the destruction of which is largely attributed to severe light attenuation caused by their bloom densities. Based on the presence of many genes likely involved in degrading various components of dead/decaying plant material (largely including those linked to recent LGT discussed above), both HAB-forming species could be capable of relying on different components of plants for nutrients during periods of increased reliance on heterotrophic metabolism. This could act as an important positive feedback loop similar to that proposed for other aspects of these HABs (e.g., ref. [54]), where dead/decaying plant material allows these algae to proliferate during bloom conditions, which causes increased shading and further destruction of sea-grass beds, in turn providing further substrate for organic nutrients accessible to these algae.

### *LGTs related to cofactor/vitamin transport and biosynthesis*

Some species-specific recent LGTs were related to cofactor/vitamin biosynthesis. The discovery of different recent pelagophyte LGTs related to biotin, an important cofactor in many metabolic processes, suggests some species have acquired genes that serve to increase biotin within their cell using different approaches – increased transport or synthesis. In *P. calceolata* a high-affinity bacterial-type biotin transmembrane transporter (BioY) was identified (SINGLETON_1119), suggesting that it has an additional and unique biotin transport system to uptake and outcompete other algae for biotin, which is often limited in some parts of the open ocean[55,56]. Phylogenetic analyses suggested that the BioY gene in *P. calceolata* is derived from a recent LGT from within Firmicutes or Actinobacteria (35% identity) either directly or indirectly via eukaryote-eukaryote LGT from a dinoflagellate, as a close homolog was identified in *Crypthecodinium cohnii* (Fig. 5B). The BioY-encoding gene was expressed under normal culture conditions (72 TPM) and had homologs in independently sequenced strains of *P. calceolata* (ref. [57] and MMETSP derived data), suggesting that it has a functional role within this alga and is not the result of contamination. This biotin transport system in bacteria sometimes involves two other genes to form a tripartite complex of BioMNY proteins. Many firmicutes, however, only encode *bioY*, which has been shown to encode functional biotin transporters on their own[58]. As this type of biotin transport system is not typically found in eukaryotes[58] and was not identified within the other pelagophytes, it likely represents a novel way for *P. calceolata* to uptake biotin from its environment more effectively. *Au. lagunensis*, on the other hand, harbors an LGT related to biotin biosynthesis (BioA) that was found to branch separately from BioA homologs in other eukaryotes (including a BioA-encoding orthogroup found in all pelagophytes) and was likely acquired via a LGT from alpha-proteobacteria (54% identity) (SINGLETON_3273).As BioA genes are involved in biotin biosynthesis, having an additional copy of the enzyme could allow for increased biotin production capabilities, providing a competitive advantage in biotin-limited environments.

*Ac. anophagefferens* encoded a species specific LGT for the rate limiting step of pantothenate (vitamin B5) biosynthesis (ketopantoate reductase; PanE)[59]. The LGT-derived *PanE* in *Ac. anophagefferens* (N0_HOG0019626) had best blast hits to mostly delta/alpha-proteobacteria (37% identity) but was ultimately of unclear bacterial origin and possibly involved an additional eukaryote-to-eukaryote LGTs involving haptophytes or cryptophytes. One other orthogroup found ancestral to pelagophytes was identified as a PanE-encoding gene (N0_HOG0004448); this gene had homologs in some other stramenopiles as well as *Symbiodinium* and select haptophytes prior to branching within gamma-proteobacteria (38% identity), suggesting it may be a more ancient LGT or EGT. Having an additional copy of PanE could provide metabolic flexibility and increased production of pantothenate in *Ac. anophagefferens*. A similar role for gene duplication within the pantothenate biosynthesis pathway has been inferred in bacteria, particularly for PanE, which, as mentioned above, is the pathways rate limiting step[59].

### *LGTs related to nucleotide salvage and biosynthesis*

A few key differences in nucleotide biosynthesis and salvage pathways between the pelagophytes were also attributed to recent LGTs. One of the most notable cases was the unique presence of a bacterial thymidine kinase gene in *P. calceolata (*SINGLETON_933) that functions in the first step of the thymidine salvage pathway commonly observed in bacteria and plants (e.g., ref. [60]). Phylogenetic analyses of the thymidine kinase in *P. calceolata* showed it branching deeply embedded within alphaproteobacteria (Fig. 5A). The high identity of this recent LGT to its closest bacterial homologs (~70-72%) challenges the proposed '70% rule' for LGT in eukaryotes[61]. Other homologs to this gene were present in other strains of an independently sequenced genome of *P. calceolata* RC100[57] that branched together in corresponding phylogenetic trees, providing evidence to refute their presence being due to contamination. Long-reads from genome sequencing data were seen to span this gene and clearly eukaryotic neighboring genes, providing further support for its genuine integration within the genome. Furthermore, RNA-seq data showed that the thymidine kinase was highly expressed (50 TPM), providing support for its functionality. The presence of this gene, along with two predicted equilibrative nucleoside transporters that could uptake thymidine, could allow *P. calceolata* to effectively recycle thymidine from its environment rather than solely rely on *de novo* synthesis and thus would foreseeably provide a significant advantage over other algae who lack this pathway, especially during certain nutrient limitations.

In *Au. lagunensis*, an expressed (65 TPM) purine nucleoside permease (SINGLETON_3604) was acquired via LGT that could play an important role in salvaging purine nucleosides from its environment. Phylogenetic analysis of the predicted purine nucleoside permease showed *Au. lagunensis* and related Sarcinochrysidales species branching with homologs in select haptophyte and dinoflagellate species, which in turn branched within a large clade of alphaproteobacteria (40% identity). The cumulative absence of other stramenopile species and patchy distribution amongst these algal groups suggests that this was likely a bacterial gene originally acquired via LGT in eukaryotes that has been subsequently passed around via eukaryote-to-eukaryote LGT.

An enzyme for the final step of the biosynthesis pathway for the modified nucleoside queuosine was identified uniquely in *P. calceolata* (QueF) (SINGLETON_1610). Phylogenetic analyses of the QueF gene in *P. calceolata* suggested that it was likely acquired from either gamma-proteobacteria or Desulfobacteria (47% identity) and involved a eukaryote-to-eukaryote LGT either from or to haptophyte species involved. While both prokaryotes and eukaryotes use queuosine as tRNA modifications to stabilize codon interactions during protein synthesis (especially during oxidative stress), *de novo* queuosine biosynthesis is uncommon within eukaryotic cells, which typically salvage it from their environment instead[62]. Although other enzymes for earlier steps of the queuosine biosynthesis pathway were not found, the presence of

QueF suggests *P. calceolata* is uniquely able to rely at least partially on its own synthesis of queuosine instead of obtaining it solely from the environment[62]; this could be particularly advantageous when queuosine is in low abundance but the preQ1 precursor is present. As queuosine is involved in stabilizing codon interactions, this gene could be crucial in enhancing the accuracy and/or efficiency of protein synthesis in *P. calceolata*, particularly during oxidative stress[62].

### *Other LGTs associated with transporters*

While a handful of species-specific transporters were derived from recent LGT within *Au. lagunensis*, one orthogroup was interestingly related to metal ion transmembrane transport (N0_HOG0023616) and specifically annotated as an NRAMP family Mg2+/Fe2+ transporter (MntH). Phylogenetic analyses indicate that this gene was likely acquired from an actinobacteria (30% identity) and was unrelated to other NRAMP family genes in eukaryotes (data not shown). While three other orthogroups were identified within *Ac. anophagefferens* with this same domain, they were clearly more ancestral genes with homologs in related stramenopiles and/or other algae. The more recently acquired *MntH* in *Au. lagunensis* suggests differences in metal ion transport exist between the two HAB forming pelagophytes. Additionally present in *Ac. anophagefferens* was a monocarboxylate transporter acquired from alpha- proteobacteria (39% identity) (N0_HOG0011619) and a sugar phosphate permease (UhpC) (N0_HOG0016908) likely acquired from a Terrabacteria-group bacterium (40% identity) indirectly from a dinoflagellate via eukaryote-to-eukaryote LGT.

### *LGTs related to other aspects of metabolism*

Other recent LGTs specific to either HAB species were likely advantageous in phosphorus limited environments, such as peak pelagophyte blooms[63–65], by contributing to flexibility in organic phosphorous containing compounds used in metabolism. This included a highly expressed (250 TPM) bacterial ComB-like phosphatase in *Au. lagunensis* (SINGLETON_2873). Phylogenetic analysis of the ComB-like gene showed the *Au. lagunensis* homolog branches sister to a clade of dinoflagellates with maximum support that was then embedded within planctomycetes (40% identity), suggesting that the bacterial gene was subject to at least one eukaryote-to-eukaryote LGT after being initially acquired. ComB-like acid phosphatases are uncommon in eukaryotes and have only been annotated in *Entamoeba*, select dinoflagellates and a few Bilateria/Porifera species; instead, they are common in bacteria, where they are associated with the metabolism of phosphate from organic compounds containing phosphate esters; encoding this enzyme could be particularly advantageous in environments where phosphate is limited.

A 2-aminoethylphosphonate-pyruvate transaminase in *Ac. anophagefferens* (N0_HOG0016040) was likely acquired from a verrucomicrobia or CFB group bacteria (51% identity) and involved a eukaryote-to-eukaryote LGT with dinoflagellates. This enzyme produces both alanine (feeding into nitrogen metabolism and protein synthesis) and ethylphosphonate from 2-aminoethylphosphonate, a substrate that can be abundant in coastal marine ecosystems from anthropogenic origins such as pesticide use and agricultural run-off[66]. As such, the ability to uniquely utilize this phosphonate as an organophosphorus source could be particularly advantageous in *Ac. anophagefferens'* environment, particularly during nutrient limited conditions. One other class V aminotransferase found across all pelagophytes was predicted to be a recent LGT specifically involving an Ectoine hydroxylase-related dioxygenase (N0_HOG0007781). Phylogenetic analysis showed pelagophyte homologs branching with select dinoflagellates and haptophytes in a monophyletic clade that was then embedded within a large clade of predominately acidobacteria (46% identity). Nearly all of the rest of the class V aminotransferases predicted across pelagophytes (14/17) had strong bacterial signals with differing presence/absence patterns in other eukaryote species suggestive of more complicated gene transfers, more ancient transfers, or EGT type scenarios (e.g., N0_HOG0019884 which had close homologs in oomycetes and some other stramenopile species, green algae, bivalves and cyanobacteria). Overall, this suggests that this class of enzymes may be more frequently transferred from bacteria to and between algae, playing an important role in the range of amino acid and nitrogen metabolism found in different species.

One other species-specific recent LGT in *Au. lagunensis* included an expressed (40 TPM) bacterial-type ubiquitin activating enzyme (ThiF) (SINGLETON_2666) that branched with a variety of CFB-group bacteria (45% identity). In bacteria, ThiF is involved in thiamine biosynthesis via the activation of a sulfur carrier protein (ThiS) and can participate in processes that resemble eukaryotic ubiquitin-activating enzymes[67]. A ThiS-related gene was not annotated in *Au. lagunensis* nor were any regions of its genome identified as homologous to ThiS genes from a variety of bacteria; the ThiS gene, however, is typically short (80-90 amino acids) and thus could have gone undetected in gene prediction pipelines or be too divergent and short to produce significant blast results. Without an obvious *ThiS* gene present, it is unlikely that ThiF functions in similar thiamine biosynthesis processes within *Au. lagunensis.* Instead, it could be co-opted for a different metabolic function, possibly in pathways for the synthesis of another sulfur-containing compound or in a role analogous to eukaryotic ubiquitin-like enzymes.

One metabolic enzyme that appears to have been acquired via LGT on multiple separate occasions is taurine dioxygenase (TauD). TauD enzymes catalyze the oxidative cleavage of taurine, producing sulfate/sulfite and amino-acetaldehyde and are thus involved in both taurine sulfur metabolism within cells and enables them to use taurine as a carbon source; it also can play a role in neutralizing ROS. Over 20 orthogroups with this domain were annotated within the pelagophytes, including 10 found in *Ac. anophagefferens* specifically, suggesting it has an increased capacity to metabolize taurine, which may be important during specific nutrient limitations. Most of the TauD orthogroups found differentially amongst species appeared to be the result of differential duplications of ancestral pelagophyte TauD genes; many of these had complicated phylogenies with patchy distributions in eukaryotes, such that they were not confidently assigned as recent LGTs with the criteria used in this study despite having strong bacterial affiliations. Two TauD-containing orthogroups were confidently assigned as recent LGTs that were either specific to *P. calceolata* (N0_ HOG00105721; acquired from actinobacteria (36% identity)) or *Au. lagunensis* (N0_HOG00105722; acquired from alpha-proteobacteria (57% identity)). Low identity to one another (~20%) combined with non-overlapping blast hits suggested these species-specific LGTs of *TauD* were unlikely to have been acquired by the same LGT event.

Recent LGTs specific to *P. calceolata* included Myo-inositol oxygenase (SINGLETON_2242) that, on the basis of phylogenetic analyses, had unclear bacterial origin (35% identity). This recent LGT in *P. calceolata* appears to have replaced a more ancestral version of this gene identified in other pelagophytes and stramenopiles from which it was highly divergent (23% identity across 20% of its length). While it is tempting to speculate the recently acquired version of this Myo-inositol oxygenase results in improved functionality within *P. calceolata*'s environment, why it harbors a different Myo-inositol oxygenase than other pelagophytes and stramenopile algae is unclear. Also present is an enoyl-CoA hydratase/carnithine racemase (CaiD) involved in fatty acid degradation (N0_HOG0022084) that branched within a clade of *Acidimicrobiaceae* bacterium (42% identity) that could allow for increased flexibility/efficiency in utilizing fatty acids as an energy source within *P. calceolata*, and an acetoacetate decarboxylase (SINGLETON_1574) that branched sister to two haptophyte species within a larger clade of mixed bacteria, suggestive of eukaryote-to-eukaryote LGT of an originally acquired bacterial gene that could allow for enhanced energy production under low-oxygen conditions and varying carbon source availability (e.g., in plants[68]).

Other notable recent LGTs in *Au. lagunensis*-specifically included a malic enzyme (SfcA) (SINGLETON_3663) that branched specifically within Fibrobacteria species (68% identity), and separately from two other malic enzymes found within all pelagophytes that were more ancestral to stramenopiles (data not shown). While this gene was not found to be expressed under normal culture conditions, an identical homolog from an independently sequenced *Au. lagunensis* transcriptome was identified in the MMETSP database, providing support for its active transcription under some circumstances and for its legitimate presence within the genome. As malic enzymes play key roles in carbon metabolism, having an additional malic enzyme could provide numerous advantages related to energy generation during nutrient stress or fluctuating nutrient availability, conditions commonly encountered in HABs.

**Other LGTs in pelagophytes classified as transferases/associated with RNA modifications**

Outside of the LGTs discussed above, *Ac. anophagefferens*-specific LGTs included two protein-tyrosine sulfotransferases (N0_HOG0020162 and N0_HOG0020448) that have very low similarity to one another (<20% identity) and non-overlapping blast hits to different subsets of gamma-proteobacteria species (~30% identity), suggesting they were acquired separately. Also present is a *Ac. anophagefferens*-specific recent LGT annotated as a mannosyl-glycoprotein 4-beta-N- acetylglucosaminyltransferase (N0_HOG0020598) likely acquired from alpha/gamma-proteobacteria (38% identity) that had close homologs in a handful of dsDNA viruses and an uncultured phage within the smallest highly supported *Ac. anophagefferens*-bacterial clade.

Recent LGTs specific to *P. calceolata* were most significantly enriched in a variety of methyltransferases (e.g., two S- adenosyl dependent protein methyltransferases), thymidine kinase activity (discussed above) and the transfer of substituted phosphate groups. This included a protein-lysine N-methyltransferase (SINGLETON_1280) and an isoprenylcysteine carboxyl methyltransferase (SINGLETON_830) both likely obtained from cyanobacteria (30-37% identity) and involved in protein metabolism via post-translational modifications. Phylogenetic analyses of the isoprenylcysteine carboxyl methyltransferase gene showed the *P. calceolata* homolog branching directly with *Symbiodinium* spp. within a larger clade of green algae and bacteria; the overall species presence and specific branching pattern was strongly suggestive of *P. calceolata* and *Symbiodinium* having acquired this bacterial gene from green algae. The specific affiliation with cyanobacteria suggests that this gene is likely related to EGT within the green algae; the incredibly patchy presence amongst complex photosynthetic algal lineages is more consistent with a LGT scenario in these algae. The recent LGT specific to *P. calceolata* associated with the transfer of other substituted phosphate groups was predicted to be a CDP-alcohol phosphatidyltransferase (SINGLETON_841) acquired from cyanobacteria (33% identity) and involved in phospholipid biosynthetic processes. Phylogenetic analyses showed this *P. calceolata* gene branching within a large clade of mostly cyanobacteria with no other eukaryotes present.

Another methyltransferase predicted to be a recent *P. calceolata*-specific LGT encodes a tRNA(Val) N6-methyltransferase domain that plays a role in tRNA modification related to gene expression fused to a phytanoyl-CoA dioxygenase (PhyH) domain involved in fatty acid metabolism (N0_HOG0003702_2). Both domains have phylogenetic affinities to proteobacteria, although to largely different species. The tRNA methyltransferase domain had closet homologs in mostly alpha-proteobacteria (46% identity) while the PhyH domain had closest homologs in gamma-proteobacteria (30% identity), suggesting that the two domains may have been acquired separately and subsequently fused. Homology searches of the gene model using both the protein and nucleotide sequence as a query did not identify any adjacent homologs that contained both domains outside of other *P. calceolata* strains, providing further support for this unique gene fusion. Importantly, the gene model for the fusion-protein was found to be expressed (~25 TPM) with uniform RNA-seq mapping across its length, suggesting (together with long-read sequencing based support) that is in fact encoded by a single gene. The fusion of two domains with distinct functions into a single gene suggests that there might be some benefit to having their expression coordinated in *P. calceolata*.

Pseudouridine synthases, which are involved in generating RNA modifications via the isomerization of uridine to pseudouridine, were predicted to be encoded by 50 orthogroups within the pelagophytes, 34 of which were specifically annotated as belonging to the RsuA/RluA-like family that is associated with rRNA (and occasionally tRNA) modifications. RsuA/RluA-like pseudouridine synthases appeared to be frequent targets of LGT within pelagophytes, with strong evidence for four separates recent LGT events from a variety of bacterial groups. Most other pelagophyte RsuA/RluA-like pseudouridine synthases that had close homologs in at least other related stramenopiles also had more ancient bacterial signals that likely reflect their ancestral presence in both bacteria and eukaryotes[69]. Within the Pelagomonadales specifically, pseudouridine synthases appear to be expanded compared to *Au. lagunensis;* 40-47 of the orthogroups contain homologs in *P. calceolata* and/or *Ac. anophagefferens,* respectively, compared to only 32 in *Au. lagunensis.* This suggests that the Pelagomonadales, and to a greater extent *Ac. anophagefferens* specifically, produce higher levels of pseudouridine in various RNA species. This could serve to increase the overall stability of RNA molecules and enhance protein synthesis within these species, which could allow them to fine-tune their response to fluctuating environmental conditions.

### *Some LGTs are related to tolerance/detoxification of anthropogenic impacts*

Some recent LGTs in *Ac. anophagefferens* or *Au. lagunensis* could be linked to mitigating the impact of other environmental stresses commonly present in their respective niches due to anthropogenic activities. Pelagophyte HABs occur in shallow estuaries (*Ac. anophagefferens)* or lagoon regions (*Au. lagunensis*) that are anthropogenically modified (e.g., from wastewater/agricultural run-off) and, as a result, contain high concentrations of metals and other potentially harmful compounds (xenobiotics) that have to be tolerated and/or detoxified. One way to deal with these compounds is via transmembrane transporters that export them from the cell. A recent LGT specific to *Au. lagunensis* (N0_HOG0000641) encodes a putative multidrug resistance protein (MdtG) that facilitates the efflux of toxins out of the cell. The MdtG gene in *Au. lagunensis* formed a monophyletic group with dinoflagellate homologs in phylogenetic analyses, which then branched as a sister group to mostly Verrucomicrobia bacteria (30% identity); this topology suggests an initial transfer of *MtdG* from bacteria to dinoflagellates, followed by eukaryote-to-eukaryote LGT into *Au. lagunensis.*

*Ac. anophagefferens* encodes an LGT-derived, species-specific tellurite-resistance/dicarboxylate transporter (TehA) (N0_HOG0018027) involved in exporting the potentially toxic heavy metal and preventing ROS related damage. Phylogenetic analyses of the TehA gene identified a single homolog in a green algae and kinetoplastid species that branched with *Ac. anophagefferens*, suggesting that additional eukaryote-eukaryote gene transfers were involved in acquisition of an originally acquired gene from Gammaproteobacteria (32% identity) (Fig. 5F); the directionality of LGT between these eukaryotes, however, is unclear. Another recent LGT present in both HAB-forming species but absent in *P. calceolata* was an arsenite transmembrane transporter related to arsenic detoxification. Genes related to mitigating arsenic toxicity via transport were previously noted to be present in *Ac. anophagefferens*[15] but were not attributed to recent LGT. Two arsenic transmembrane transporters (ArsA) were identified within the *Ac. anophagefferens* pangenome – a core orthogroup of likely EGT origin present in all pelagophytes (N0_HOG0005839) and a clearly distinct accessory orthogroup acquired by recent LGT that has been subsequently lost in CCMP1850 that was also present in *Au. lagunensis* (N0_HOG0015620). While this accessory gene does not provide a novel function to strains, it could result in differences in their abilities to detoxify arsenic. Phylogenetic analyses showed that the LGT-derived ArsA genes in pelagophytes formed a monophyletic clade, which then branched exclusively with a variety of bacterial species (46% identity), making the exact donor lineage unclear. The presence of genes related to arsenic detoxification acquired via prokaryotic LGT has been observed in the red alga *Galdieria*, who also encodes a number of LGTs related to adaptation to its extreme environments[70], though they were not closely related to the LGT derived arsenite transporter in HAB-forming pelagophytes.

Other recent LGTs appeared to be involved in catalytic activities that detoxify/metabolize xenobiotic compounds, breaking them down into less harmful or more easily excretable compounds. This included bacterial arylsulfotransferase genes present in both HAB-forming pelagophyte species (N0_HOG0011980) and specifically in *Ac. anophagefferens* (N0_HOG0016186) via separate LGT events potentially associated with the detoxification of various xenobiotic compounds and phenolics through sulfation[71]. This gene family was highly expanded in *Ac. anophagefferens* compared to other pelagophytes, encoding 20 additional orthogroups that were largely derived from gene duplication of a species-specific LGT that have diverged substantially from one another, suggesting the recent LGT plays an important role in detoxification and/or sulfur metabolism. The absence of this type of gene in *P. calceolata* but presence in an additional LGT shared between *Au. lagunensis* and *Ac. anophagefferens,* as well as the overall expansion of this gene family within *Ac. anophagefferens*, suggests that bacterial-type arylsulfotransferases might be particularly advantageous in these HAB forming species.

Carboxylesterase type B genes (PrbA) were also acquired by LGT on at least two occasions within all pelagophytes and specifically within *Ac. anophagefferens* (N0_HOG0001696_2) likely derived from an alphaproteobacteria (38% identity) either directly or indirectly via a eukaryote-eukaryote LGT involving dinoflagellates or the haptophyte *Chrysochromulina tobinii*. As these enzymes generally function in lipid metabolism and detoxification of a wide range of xenobiotics by hydrolyzing ester bonds (e.g., pesticides and various drugs)[72], these LGTs likely help these algae cope with a variety of environmental toxins and contribute to the breakdown of lipids for energy production in the cell. The presence of a signal peptide suggests that this LGT could function extracellularly, degrading organic compounds for either defense against toxins or generating substrates for metabolism.

Some enzymes acquired by LGT were predicted to be involved in degrading various commonly used antibiotics that can be present in anthropogenically modified environments from wastewater[22,73], including an erythromycin esterase (N0_HOG0013682) (as previously noted to be present in the genome by [21]) acquired in the ancestor of the Pelagomonadales from beta/gamma-proteobacteria (37% identity to *Ac. anophagefferens*) and two separate LGTs of beta lactamases involved in the degradation of penicillin and cephalosporins in either *Ac. anophagefferens* (N0_HOG0021864) or *Au. lagunensis*. 17 other beta-lactam related orthogroups not attributed to recent LGTs were also identified across pelagophytes that largely appear to be the result of differential gene duplications of a beta-lactam gene found in a common ancestor of the pelagophytes – there was, however, a clear increase in copy number within *Ac. anophagefferens* compared to the other pelagophytes (16 orthogroups had homologs in *Ac. anophagefferens* vs. 5/19 in *P. calceolata* and 8/19 in *Au. lagunensis*). While the universal presence of these genes suggests all pelagophyte species may have the ability to metabolize beta-lactams, the increased copy number of beta-lactamase related genes in the case of *Ac. anophagefferens*, including an additional version acquired by recent LGT, The degradation of these antibiotics could serve both detoxification purposes (e.g., see ref. [74] and references therein) and/or provide substrates for further biosynthetic processes, acting as a source of carbon and/or nitrogen.

An additional recent LGT within *Au. lagunensis* was predicted to be involved in the metabolism of cocaine, a drug that could be present in its environment due to anthropogenic activity (e.g., ref. [75]). The presence of a CocE/NonD family hydrolase (SINGLETON_4314) acquired from spirochaetes (50% identity) suggests *Au. lagunensis* could potentially metabolize cocaine as an alternative carbon source as has been observed in some bacteria that also encode this enzyme[76].

### *Some other LGTs are related to dealing with oxidative and osmotic stress*

Pelagophyte HABs occur in environments with moderate to high levels of salinity (>25 ppt in *Ac. anophagefferens* and >40 ppt in *Au. lagunensis*) and high levels of ROS that increase during blooms. In order to tolerate varying degrees of osmotic and oxidative stress, algae rely on different mechanisms to regulate osmotic balance (e.g., osmolyte production, utilizing ion channels or medication of cell walls), remove ROS and repair oxidatively damaged DNA/proteins. In *Ac. anophagefferens*, a laterally acquired bestrophin-like chloride channel gene (yneE) (N0_HOG0017299) from CFB group bacteria (24% identity) was identified that has known roles in maintaining osmotic balance within plant cells and regulating photosynthesis during fluctuation light conditions[77].

On the other hand, a highly expressed (150 TPM) bacterial ) glycerol-1-phosphate dehydrogenase (G1PDH) was identified in *Au. lagunensis* (SINGLETON_2688) that could increase its tolerance to osmotic stress by contributing to cell membrane stability or the structure and composition of their EPS, a feature of *Au. lagunensis* that has been previously implicated in tolerance to salt related stress[78]. Phylogenetic analyses suggests that the G1PDH was acquired from a *Phycisphaerae* bacterium or another planctomycete (49% identity) in *Au. lagunensis* specifically. Glycerol-1-phosphate dehydrogenases are not commonly found in eukaryotes; instead, they typically encode glycerol-3-phosphate dehydrogenases (G3PDH) that are central to their lipid metabolism. In contrast, G1PDH enzymes are usually associated with archaea who use this enzyme for membrane lipid biosynthesis. The exact function of G1PDH homologs thought to have also been acquired by LGT in select bacteria like planctomycetes is still unclear[79]. The unique presence of a G1PDH in *Au. lagunensis* could result in enhanced glycerol metabolism (glycerolipid biosynthesis), or increased membrane stability and tolerance to osmotic and other environmental stress by utilizing both G1PDH and G3PDH products in their cell membrane[80].

The *Au. lagunensis* genome also encodes an expressed (50 TPM) bifunctional PutA gene (proline dehydrogenase and 1-pyrroline-5-carboxylate dehydrogenase) acquired by recent LGT from alpha-proteobacteria (46% identity) (SINGLETON_3749). Similarly to the G1PDH discussed above, PutA genes are not typically found in eukaryotes, with only a handful of distantly related homologs are annotated in other plants and algae (e.g., *Arabidopsis thaliana*[81]). Instead, they are commonly associated with bacteria where they are involved in proline catabolism to form glutamate, contributing to nitrogen metabolism and cellular osmotic balance[82]. While specific roles in *Au. lagunensis* are not known, such functions associated with PutA in bacteria could be highly advantageous to this algal species, particularly during HABs.

Subclasses of oxidoreductases involved in ROS mitigation – generally superoxide dismutases, catalases and peroxidases – were differentially present and abundant amongst the pelagophyte species, although largely they were not inferred to be recent LGTs. The exception was an additional Cu/Zn-superoxide dismutase specifically acquired in *Au. lagunensis* (SINGLETON_2577) and an alkyl hyperoxide reductase acquired in the common ancestor of all pelagophytes that has since been differentially duplicated twice within select *Ac. anophagefferens* strains (N0_HOG0020729, N0_HOG0022030). Compared to the other pelagophytes, the *Ac. anophagefferens* pangenome contains ~1.5X more orthogroups associated with antioxidant activity, ~15% of which were present in the accessory genome. This is likely linked to differential gene duplications of core genes and could allow for strains to differentially tailor their response to reactive oxygen species and detoxify various compounds related to oxidative stress, overall maintaining redox balance in the cell. Algal blooms of *Ac. anophagefferens* occur in anthropogenically modified environments where heavy metals and chemical pollutants are abundant, which in turn promote ROS accumulation – as such, it is likely that having additional copies of genes associated with antioxidant activities and/or response to oxidate stress would be advantageous.

In general, some genes associated with various metabolic pathways can have additional roles in reducing oxidative stress by utilizing potentially harmful metabolites as substrates. One example is A-like cyclophilins derived from LGT in *Ac. anophagefferens* (N0_HOG00082402) associated with peptidyl-prolyl isomerization. Cyclophilin-type domains were found in a large number of other orthogroups across the pelagophytes that were divergent from this *Ac. anophagefferens*-specific LGT on the basis of both sequence identity and largely non-overlapping best-blast hits. Cyclophilins have diverse roles in the cell from protein folding and chaperone type activity, to implications in stress response in plants[83], immune response (e.g., ref. [84]) and oxidative stress response in dinoflagellates[85]. Compared to the other pelagophytes, cyclophilin-type domains appeared highly expanded in *Ac. anophagefferens* with 22 species-specific orthogroups identified compared to four orthogroups specific to either *Au. lagunensis* or *P. calceolata*. Additional copies of the domain, including the recent species-specific LGT, may have a key role in allowing correct protein folding, particularly under oxidative stress (e.g., ref. [85]). The increased prevalence in high GC content species suggests its importance may be even more crucial in these cases given the higher frequency of 'GARP' amino acids they encode.

Similarly, glutathione-S- transferases have predicted functions in detoxification of ROS and other xenobiotic compounds by conjugating reduced glutathione to them. These enzymes were transferred on at least three separate occasions in different HAB-forming species, where acquisition was also followed by differential duplications of initially acquired bacterial genes. Glucose-methanol-choline oxidoreductases were also recently transferred into pelagophytes on at least three separate occasions – once within *P. calceolata* specifically (N0_HOG00049021) from gamma-proteobacteria (57% identity) and twice in *Ac. anophagefferens* from bacteria/euryarchaeotes (37% identity) (N0_HOG0014623) or actinobacteria (38% identity) (N0_HOG0004182_1) – and are associated with diverse metabolic activities related to carbon metabolism, as well as management of osmotic stress and ROS. Genes between these orthogroups align to one another with only 30-37% amino acid identity, have largely non-overlapping blast hits and branch separately within phylogenetic trees (data not shown), suggestive of multiple independent transfers. The increased gene copy number within *Ac. anophagefferens* partly attributed to species-specific LGTs (9 genes per strain) compared to *P. calceolata* (3 genes) combined with the presence of three orthogroups retained only in the two HAB forming species could suggest a particular advantage for these enzymes in the ecological context of HABs formed by *Ac. anophagefferens* and *Au. lagunensis*.

Polyamines have diverse metabolic roles in the cell related to nitrogen metabolism, regulating cellular processes and in stress response, including defense against oxidative damage by neutralizing ROS and enhancing antioxidant enzyme activity (e.g., ref. [86] and references therein). Both genes for putrescine biosynthesis (an agmatine deaminase (AguA)) and a nitrilase CPA domain (AguB)) were uniquely present in a single ORF in *Ac. anophagefferens*

((N0_HOG0015062). While there was no evidence for active transcription under normal culture conditions, the gene model is comprised of a single 730 amino-acid open-reading frame and has long-read genomic support, suggesting the two domains are part of the same gene. While some species of red algae, green algae and diatoms have been noted to encode both of the enzymes involved in putrescine biosynthesis[86], homologs to the *Ac. anophagefferens* gene in other eukaryotes that encoded the AguA domain did not also contain the AguB domain and vice-versa, suggesting that the AguA+AguB domain structure in a single protein-coding gene is unique within *Ac. anophagefferens* (and the amoeba *Vannella* sp.).

When database-retrieved homologs were required to span both domains, homologs were only identified in euryarchaeotes (31% identity) and a single amoeba species (*Vannella* sp.), which branched with *Ac. anophagefferens* sequences in phylogenetic analyses. When considered separately, the nitrilase domain (AguB) had best blast hits to firmicutes specifically (45% identity), while the AguA2 domain had best blast hits to different CFB-group bacteria (38% identity). Phylogenetic analyses of each domain considered in isolation showed eukaryotic homologs to the AguA domain branching somewhat nearby the *Ac. anophagefferens* domain (data not shown); on the other hand, eukaryotic homologs to the AguB domain branched separately from a highly supported *Aureococcus*-bacterial clade. Notably, in bacteria genes for AguA and AguB have been found to be encoded within the same operon (e.g., ref. [87]). This could suggest that this gene was initially acquired from a bacterial operon in euryarchaeotes and subsequently transferred to *Ac. anophagefferens.* It is also possible that just the AguB domain was acquired recently via LGT from bacteria – ultimately, however, the history is unclear as to how or when both domains were acquired and/or fused within *Ac. anophagefferens*. In any case, this suggests that putrescine biosynthesis is related to recent LGT in some way or another (although the exact history remains unclear) and has a significant role in the overall stress response in *Ac. anophagefferens* and the coordinated expression of both genes is important for this response.

A variety of different DNA/protein repair related LGTs were identified amongst the pelagophytes; similar patterns of increased LGT within DNA repair pathways has been observed in some other protist species ) and could be crucial for maintaining genome integrity (e.g., *Entamoeba histolytica*[88]). These LGTs included a unique DNA alkylation damage repair pathways (AdaB) and an additional AlkB gene in *Au. lagunensis*; this suggests LGTs have contributed to an increased ability for repair of alkylated DNA in *Au. lagunensis* that is induced by environmental stressors. Phylogenetic analyses of the highly expressed (250 TPM) O6-methylguanine-DNA- protein-cysteine methyltransferase (AdaB) (N0_HOG0023750) in *Au. lagunensis* showed this gene branching within alpha-proteobacteria (60% identity). Outside of some metazoa, this specific family of proteins is uncommonly annotated in eukaryotes, who rely on different repair pathways to handle DNA alkylation damage[89]; the unique presence of this gene in *Au. lagunensis* could therefore be advantageous by providing additional DNA repair mechanisms and an increased ability to repair damage induced by environmental stress common in algal blooms (e.g., environmental pollutants and ROS), thus helping maintain its genomic integrity.

Also related to DNA alkylation damage repair was an LGT derived orthogroup containing two highly expressed, nearly identical genes (>1,000 TPM) in *Au. lagunensis* annotated as alkylated DNA repair proteins (AlkB) (N0_HOG0021012_1). The LGT-derived AlkB genes had high identity to both viruses (45%) and gamma/beta-proteobacteria (40%). Phylogenetic analyses showed the *Au. lagunensis* homologs branching sister to an unclassified Euryarchaeota archaeon and within a larger, highly supported clade that otherwise contained sequences from mostly NCLDVs and a handful of unclassified bacteria, suggesting that gene transfer occurred from viruses into *Au. lagunensis* or vice versa. The presence of an additional AlkB gene in *Au. lagunensis* could result in an increased capacity to repair DNA alkylation related damage, again playing a role in maintaining its genome integrity.

Within *Ac. anophagefferens*, recent LGTs were responsible for an additional version of a peptide methionine sulfoxide reductase gene (MsrB) (N0_HOG0015743) involved in oxidative protein repair and a mismatch repair enzyme (MutS) involved in DNA mismatch repair in the *Ac. anophagefferens* accessory genome (N0_HOG0024196). Phylogenetic analysis of the MsrB orthogroup resolved *Ac. anophagefferens* homologs as directly sister to select dinoflagellates, which then branched separately from other eukaryote sequences within a highly supported bacterial clade of Gammaproteobacteria (52% identity). Five other orthogroups encoding a predicted MsrB domain were identified within the pelagophytes – two additional *Ac. anophagefferens*-specific orthogroups and three with differing presence/absence patterns in the other pelagophytes species that had non-overlapping blast hits and thus are not likely related by gene duplication. Phylogenetic analyses of the MutS accessory gene showed that the gene was likely acquired from a Campylobacterota bacterium (47% identity) and that the gene transfer was mediated by AaV or has been transferred to AaV from these strains subsequently.

A number of separate LGTs of uracil DNA glycosylases were identified across pelagophytes, and especially within the Pelagomonadales, that could contribute to more robust repair of oxidatively induced cysteine deamination, especially in the higher GC genomes of *Ac. anophagefferens* and *P. calceolata.* Uracil-DNA glycosylase family 1 proteins (UDG-F1) within the pelagophytes often had strong bacterial signals indicative of either recent LGT or more ancient LGT events present in other stramenopile and/or algal groups, including a number of duplications of recent LGTs that have contributed to differences within the accessory pangenome. Phylogenetic analyses supported at least three separate recent LGTs of this gene family – one shared in all pelagophyte species that was acquired from firmicutes (53% identity) (N0_H0G0007043), one shared between *P. calceolata* and *Ac. anophagefferens* acquired from an alpha-proteobacteria (37% identity) (N0_HOG0013717) and one exclusively in *P. calceolata* acquired from a different subset of firmicute species (43% identity) (SINGLETON_2148). One other UDG-F1 orthogroup in pelagophytes (N0_HOG0001990) had closely related homologs in other stramenopiles and was more anciently affiliated with CFB-group bacteria (45-54% identity) with phylogenies suggestive of more ancient LGT events. Notably, the other two UDG-F1 orthogroups found exclusively in *Ac. anophagefferens* are part of accessory genome (N0_HOG0017745 and N0_HOG0023927) and appeared to be separate duplications of the LGT found in all pelagophytes. Altogether this suggests that this group of DNA repair enzymes were horizontally acquired from bacteria and subsequently duplicated on multiple separate occasions within the pelagophytes. Notably, 3-4 more copies of UDG-F1 genes were identified in the pelagophyte species with high genome GC content (*Ac. anophagefferens* and *P. calceolata*) compared to *Au. lagunensis,* a pelagophyte species that has a substantially lower average genome GC content. The correlation of specific base-excision repair pathway presence/absence with low GC content has been observed previously in bacteria (e.g., ref. [90]) and eukaryotes (e.g., ref. [88]), while the presence of uracil-DNA glycosylases are specifically expected to favor high GC genomes (although this was not found to be a significant correlation in bacteria)[91].

### *LGTs potentially related to impacts on/interactions with co-occurring species*

Previous studies have aimed to assess how these pelagophytes directly impact other phytoplankton and bivalve species via the production a specific toxin that is not yet known (e.g., refs. [92,93]) and how they deter grazing (e.g., refs. [29,94–96]). Several Hydroxymethylglutaryl (HMG)-CoA synthases associated with polyketide synthase domains were identified specifically within *Ac. anophagefferens* that varied amongst strains and had a more ancient bacterial signal (i.e. they were not considered recent LGTs by our criteria). HMG-CoA synthase domain-containing regions within the accessory genome had 80-100% identity to one another within an orthogroup and appeared to be related by differential duplication in select strains (e.g., N0_HOG0004238, N0_HOG0000781). HMG-CoA synthase domains were closely associated with adjacent polyketide synthase and enoyl-CoA hydratase domains, as well as phosphopantetheine binding attachment sites; notably, these genes were located within larger regions of lower-than-average GC content (~53%) and often occurred in close proximity on a single contig (30-100 kbp in length), suggestive of a genomic island. The specific domain combinations in similarly large genomic regions have been recently linked to 'giant' polyketide synthesis in some haptophytes that are suggested to be related to toxin production[97]; these 'giant' polyketides in *P. parvum*[97] were intriguingly similar in structure and sequence to the polyketide synthase regions observed here in *Ac. anophagefferens*. These polyketide synthase-containing regions may play a similar role in *Ac. anophagefferens* as inferred in *P. parvum*, including potentially related to inter-strain differences in toxicity related to specific toxin production or dosage effects correlated with gene copy number variation. Further assessment of these polyketide synthase containing regions, including their overall domain structure and differing abundance between strains, is needed to draw conclusions related to similarity to the toxin associated giant polyketide synthase domains in *P. parvum*[97].

A T4-type lysozyme (RrrD) acquired by LGT in *Au. lagunensis* (SINGLETON_2610) could be involved in antibacterial activity as this has been noted to occur in bivalves that encode this enzyme[98]. Phylogenetic analysis showed the *Au. lagunensis* gene branching specifically as sister to an unclassified Halieaceae bacterium within a highly supported larger clade comprised of various bacterial species that were largely poorly resolved. Particularly striking was the fact that this gene in *Au. lagunensis* shares 79% amino acid identity with its closest bacterial homologs, again breaking the '70% rule'[61] and >65% identity to several phage sequences also present in the phylogenetic tree. Transcriptome data supported active transcription (16 TPM) of the T4-type lysozyme within *Au. lagunensis* and homologs were identified in independently sequenced *Au. lagunensis* transcriptomes from the MMETSP dataset; together with support from long reads within the genome sequencing data that spanned this gene into neighboring eukaryotic gene regions, there is strong evidence for the genuine integration of this lysozyme within the genome. T4-type lysozymes are most commonly associated with bacteriophages/viruses, where they are involved in degrading the peptidoglycan layer of bacterial cell walls during infection. Although they branched separately from *Au. lagunensis* homologs identified here, other eukaryotes have been identified with a horizontally acquired T4- type lysozyme including the amoebae *Dictyostelium discoideum*[99] and bivalves where they been shown to have antibacterial activities[98]. While speculative, this gene could also play a role in enhancing antibacterial activity in *Au. lagunensis* or could be related to the breakdown of organic matter for nutrient acquisition. A signal peptide, however, was not predicted to be encoded.

Additionally, the presence of two recently acquired sialic acid synthesis genes for N-Acetylneuraminic acid in pelagophytes could play a role in cell signalling and mediating viral and pathogen-host interactions through modifications of the surface glycoproteins/glycolipids (e.g., as inferred for sialic acids in some haptophytes and dinoflagellates[100]). One of these was an N- acetylneuraminic acid mutarotase (NanM) gene with a signal peptide that was found in all pelagophytes (N0_HOG0006327) and likely acquired from alphaproteobacteria (35% identity to *Ac. anophagefferens*). While the NanM gene had one homolog per strain in *Ac. anophagefferens* and *Au. lagunensis*, five homologs were identified in the *P. calceolata* genome (~40% identical to one another) suggesting that the gene has undergone species-specific gene duplication after initial acquisition. Within eukaryotes, NanM genes have primarily been described in fungi (e.g., ref. [101]); they are more typically associated with bacterial species where they convert anomers of N-acetylneuraminic acid (i.e. sialic acid), a component of cell surface glycoproteins/glycolipids with roles in cell-cell interactions and immune response. Functionally related to the presence of NanM was a second recent LGT related to biosynthesis of N-acetylneuraminic acids involving an orthogroup found in all pelagophytes (N0_HOG0006742) that was acquired from a CFB-group bacterium (69% identity to *Ac. anophagefferens*). N-acetylneuraminic synthase (NeuB) is involved in the *de novo* biosynthesis of N-acetylneuraminic acids in bacteria and, similarly to NanM, in eukaryotes is only rarely annotated outside of Metazoa.

**A single genomic region in Au. anophagefferens with three recent LGTs**

In general, recent LGTs examined herein did not show any obvious pattern of integration within the genome – they were not localized to sub-telomeric regions and typically were not located close to one another. An interesting exception was within one genomic region in all *Ac. anophagefferens* strains that encoded three consecutive protein-coding genes predicted to be recent LGTs, albeit from varying bacterial sources. The first recent LGT was a glycoside hydrolase family 28 protein with a pectin lyase fold present in all pelagophyte species and likely acquired from a verrucomicrobia bacterium (42% identity) (N0_HOG0010666); the second was a glutathione-S-transferase (GstA) found only in *Ac. anophagefferens* that was likely acquired from cyanobacteria (55% identity) (N0_HOG0019471); and the third recent LGT was a phosphoglycerate mutase 1 gene found in *Ac. anophagefferens* and *P. calceolata* that branched specifically with beta-proteobacteria (40% identity to *Ac. anophagefferens*) (N0_ HOG0014827).

While the homologs in 4/5 strains of *Ac. anophagefferens* for these LGTs were located consecutively in a single genomic region that was supported by long reads, homologs in *P. calceolata* (when present) were located on separate contigs. The consecutive gene region was not completely conserved in 1/5 of the *Ac. anophagefferens* strains (CCMP1850), where an additional ~5,000 bp present between the pectin lyase-encoding gene and the *GstA* that encodes an additional CCMP1850-specific gene predicted to contain an ankyrin repeat, a TPR repeat and polyketide synthase domain signature that has close homologs in fungi, bacteria and also some oomycetes. Differing potential gene donors and lack of consecutive presence of these three genes in other bacteria and *P. calceolata* could suggest separate LGT events occurred in potentially different regions of the genome that have since relocated in close proximity. Given the paucity of LGTs in the genome relative to the total number of genes, it seems more likely that the LGTs occurred in a single event from a donor whose genome was already mosaic and have since moved to different genomic locations in *P. calceolata*.
