## Supplementary Tables for "Pangenome biology and evolution in harmful algal-bloom-forming pelagophyte algae"

**Supplementary Table 1 Detailed summary of genome repeats identified in all seven pelagophyte genomes from RepeatMasker.** (A) All five *Aureococcus anophagefferens* strains, (B) *Pelagomonas calceolata* CCMP1756 and *Aureoumbra lagunensis* CCMP1510.

**(A)**

| | **CCMP 1984** | | **CCMP 1850** | | **CCMP 3368** | | **CCMP 1707** | | **CCMP 1708** | |
|---|---|---|---|---|---|---|---|---|---|---|
| **Bases / % Genome** | 8.05 Mbp / 15.20% | | 8.20 Mbp / 15.19% | | 5.72 Mbp / 11.02% | | 6.18 Mbp / 11.64% | | 5.90 Mbp / 11.13% | |
| | **#** | **kbp / %** | **#** | **kbp / %** | **#** | **kbp / %** | **#** | **kbp / %** | **#** | **kbp / %** |
| **SINEs** | - | - | - | - | - | - | 14 | 0.1 0.01% | 22 | 2116 0.01% |
| **LINEs** | - | - | 213 | 143 0.3% | 63 | 9 0.02% | 38 | 12 0.02% | - | - |
| LINE1 | - | - | - | - | 40 | 5 0.01% | - | - | - | - |
| L3/CR1 | - | - | 53 | 36 0.07% | - | - | - | - | - | - |
| **LTR elements** | 480 | 250 0.5% | 796 | 438 0.8% | 388 | 247 0.48% | 500 | 236 0.45% | 458 | 242 0.4% |
| **DNA elements** | 739 | 129 0.2% | 284 | 103 0.19% | 185 | 89 0.2% | 182 | 54 0.1% | 542 | 155 0.3% |
| hAT-Charlie | 574 | 70 0.1% | 28 | 9 0.02% | - | - | - | - | - | - |
| **Unclassified** | 20,971 | 7,486 14.1% | 20,970 | 714 14.5% | 19,585 | 5,107 9.8% | 19,238 | 5,520 10.4% | 17,640 | 5,174 9.8% |
| **Interspersed repeats** | | 7,864 14.6% | | 7,824 14.9% | | 5,453 10.5% | | 5,824 11.0% | | 5,573 10.5% |
| **Small RNA** | - | - | 26 | 64 0.1% | 21 | 60 0.1% | 19 | 87 0.16% | 17 | 41 0.08% |
| **Satellites** | - | - | 128 | 66 0.1% | 54 | 6 0.01% | 26 | 17 0.03% | 10 | 0.07 0.01% |
| **Simple repeats** | 846 | 222 0.4% | 778 | 257 0.5% | 524 | 207 0.4% | 866 | 264 0.5% | 844 | 295 0.6% |

**(B)**

| | CCMP1510 | | CCMP1756 | |
|---|---|---|---|---|
| **Bases / % Genome** | 9.11 Mbp / 22.05 % | | 1.94 Mbp / 6.07 % | |
| | **Number of elements** | **Length (bp) / (%)** | **Number of elements** | **Length (bp) / (%)** |
| **SINEs** | 0 | 0 | 60 | 7794 (0.02%) |
| **LINEs** | 1297 | 533,093 (1.29%) | 0 | 0 |
| LINE1 | 28 | 9693 (0.02%) | 0 | 0 |
| LINE2 | 26 | 34,632 (0.08%) | 0 | 0 |
| L3/CR1 | 0 | 0 | 0 | 0 |
| **LTR elements** | 1629 | 2,563,216 (6.2%) | 30 | 6284 (0.02%) |
| **DNA elements** | 283 | 295,527 (0.72%) | 160 | 84,739 (0.26%) |
| hAT-Charlie | 33 | 3993 (0.01%) | 0 | 0 |
| **Unclassified** | 17312 | 5,576,471 (13.49%) | 8484 | 1,757,324 (5.49%) |
| **Interspersed repeats** | | 8,968,407 (21.7 %) | | 1,856,141 (5.69 %) |
| **Small RNA** | 35 | 56,881 (0.14%) | 0 | 0 |
| **Satellites** | 274 | 127,622 (0.31%) | 13 | 2960 (0.01%) |
| **Simple repeats** | 68 | 11,244 (0.03%) | 257 | 91,427 (0.29%) |

**Supplementary Table 2 Characteristics of introns in protein-coding gene predictions for all pelagophyte genomes.** Only canonical GT-AG and non-canonical GC-AG splice signals were evaluated. When multiple isoforms of a gene were predicted, only the longest isoform was considered in the assessment.

| | Average Intron Size (bp) | Total Introns | Splice signal | | Average Introns per gene | Genes with # of Introns | | | |
|---|---|---|---|---|---|---|---|---|---|
| | | | GC-AG | GT-AG | | 0 | 1 | 2 | 3+ |
| *Au. lagunensis* CCMP1510 | 177 | 2463 | 8<br>0.32% | 2388 | 0.135 | 16384 | 1071 | 261 | 166 |
| *P. calceolata* CCMP1756 | 124 | 3411 | 489<br>14.34% | 2186 | 0.186 | 15402 | 1434 | 280 | 125 |
| *Ac. anophagefferens* CCMP1707 | 110 | 22288 | 3519<br>15.79% | 18214 | 0.917 | 16485 | 2857 | 1721 | 2878 |
| *Ac. anophagefferens* CCMP1708 | 103 | 18752 | 2679<br>14.29% | 15821 | 0.755 | 17524 | 2974 | 1647 | 2425 |
| *Ac. anophagefferens* CCMP1850 | 129 | 13252 | 2042<br>15.41% | 10945 | 0.467 | 20088 | 4072 | 1382 | 1242 |
| *Ac. anophagefferens* CCMP1984 | 121 | 13282 | 2213<br>16.66% | 10271 | 0.477 | 19888 | 3441 | 1409 | 1272 |
| *Ac. anophagefferens* CCMP3368 | 130 | 12017 | 1885<br>15.69% | 9842 | 0.443 | 20262 | 3545 | 1335 | 1114 |

**Supplementary Table 3 Molecular function-related GO term enrichment results for LGTs specific to *Ac. anophagefferens* (see associated REVIGO plot in Figure 3B).** Only terms that were found to be significantly enriched (p-value <0.05) that occurred >1 time are included.

| GO Term | Count | p-value | Name |
|---|---|---|---|
| GO:0003674 | 239 | 1 | molecular function |
| GO:0003824 | 187 | 9.55E-19 | catalytic activity |
| GO:0005488 | 97 | 1 | binding |
| GO:0016787 | 60 | 4.08E-05 | hydrolase activity |
| GO:0016740 | 59 | 0.0103 | transferase activity |
| GO:0036094 | 58 | 0.953 | small molecule binding |
| GO:0016491 | 47 | 5.14E-06 | oxidoreductase activity |
| GO:0043167 | 46 | 0.998 | ion binding |
| GO:0097159 | 41 | 1 | organic cyclic compound binding |
| GO:0005515 | 38 | 1 | protein binding |
| GO:1901363 | 34 | 0.87 | heterocyclic compound binding |
| GO:0046872 | 29 | 0.764 | metal ion binding |
| GO:0043169 | 29 | 0.796 | cation binding |
| GO:0043168 | 28 | 0.972 | anion binding |
| GO:0140096 | 27 | 0.766 | catalytic activity, acting on a protein |
| GO:0000166 | 23 | 0.99 | nucleotide binding |
| GO:1901265 | 23 | 0.99 | nucleoside phosphate binding |
| GO:0016741 | 20 | 2.33E-04 | transferase activity, transferring one-carbon groups |
| GO:0008168 | 19 | 3.60E-04 | methyltransferase activity |
| GO:0019842 | 18 | 7.93E-05 | vitamin binding |
| GO:0030554 | 18 | 0.985 | adenyl nucleotide binding |
| GO:0017076 | 18 | 0.998 | purine nucleotide binding |
| GO:0004553 | 17 | 3.03E-08 | hydrolase activity, hydrolyzing O-glycosyl compounds |
| GO:0016798 | 17 | 5.68E-07 | hydrolase activity, acting on glycosyl bonds |
| GO:0016788 | 15 | 0.0716 | hydrolase activity, acting on ester bonds |
| GO:0046914 | 15 | 0.292 | transition metal ion binding |
| GO:0030246 | 13 | 2.92E-04 | carbohydrate binding |
| GO:0008233 | 13 | 0.154 | peptidase activity |
| GO:0032559 | 12 | 0.999 | adenyl ribonucleotide binding |
| GO:0032555 | 12 | 1 | purine ribonucleotide binding |
| GO:0032553 | 12 | 1 | ribonucleotide binding |
| GO:0097367 | 12 | 1 | carbohydrate derivative binding |
| GO:0016614 | 11 | 3.15E-04 | oxidoreductase activity, acting on CH-OH group of donors |
| GO:0008484 | 10 | 8.82E-06 | sulfuric ester hydrolase activity |

| GO Term | Count | p-value | Name |
|---|---|---|---|
| GO:0016705 | 10 | 0.0227 | oxidoreductase, acting on paired donors, incorporation or reduction of molecular oxygen |
| GO:0005524 | 10 | 1 | ATP binding |
| GO:0003676 | 10 | 1 | nucleic acid binding |
| GO:0035639 | 10 | 1 | purine ribonucleoside triphosphate binding |
| GO:0016616 | 9 | 1.78E-03 | oxidoreductase, acting on CH-OH donors, NAD acceptors |
| GO:0017171 | 9 | 5.63E-03 | serine hydrolase activity |
| GO:0008236 | 9 | 5.63E-03 | serine-type peptidase activity |
| GO:0005509 | 9 | 0.927 | calcium ion binding |
| GO:0031418 | 8 | 0.0257 | L-ascorbic acid binding |
| GO:0043177 | 8 | 0.0273 | organic acid binding |
| GO:0031406 | 8 | 0.0273 | carboxylic acid binding |
| GO:0048029 | 8 | 0.0291 | monosaccharide binding |
| GO:0008146 | 8 | 0.0576 | sulfotransferase activity |
| GO:0016782 | 8 | 0.0846 | transferase activity, transferring sulphur-containing groups |
| GO:0005506 | 8 | 0.143 | iron ion binding |
| GO:0016757 | 8 | 0.258 | glycosyltransferase activity |
| GO:0016746 | 8 | 0.796 | acyltransferase activity |
| GO:0016747 | 7 | 0.245 | acyltransferase, transferring groups other than amino-acyl |
| GO:0016772 | 7 | 0.987 | transferase activity, transferring phosphorus-containing groups |
| GO:0022857 | 7 | 0.998 | transmembrane transporter activity |
| GO:0005215 | 7 | 0.999 | transporter activity |
| GO:0047834 | 6 | 3.38E-03 | D-threo-aldose 1-dehydrogenase activity |
| GO:0004175 | 6 | 0.311 | endopeptidase activity |
| GO:0003677 | 6 | 0.914 | DNA binding |
| GO:0016773 | 6 | 0.942 | phosphotransferase activity, alcohol group as acceptor |
| GO:0016301 | 6 | 0.971 | kinase activity |
| GO:0031177 | 5 | 2.44E-03 | phosphopantetheine binding |
| GO:0072341 | 5 | 2.70E-03 | modified amino acid binding |
| GO:0033218 | 5 | 6.98E-03 | amide binding |
| GO:0051287 | 5 | 9.31E-03 | NAD binding |
| GO:0016667 | 5 | 0.0205 | oxidoreductase activity, acting on a sulfur group of donors |
| GO:0070279 | 5 | 0.0291 | vitamin B6 binding |
| GO:0030170 | 5 | 0.0291 | pyridoxal phosphate binding |
| GO:0004252 | 5 | 0.0717 | serine-type endopeptidase activity |
| GO:0008757 | 5 | 0.0945 | S-adenosylmethionine-dependent methyltransferase activity |
| GO:0016829 | 5 | 0.13 | lyase activity |
| GO:0015267 | 5 | 0.805 | channel activity |
| GO:0022803 | 5 | 0.805 | passive transmembrane transporter activity |

| GO Term | Count | p-value | Name |
|---|---|---|---|
| GO:0008270 | 5 | 0.81 | zinc ion binding |
| GO:0004672 | 5 | 0.943 | protein kinase activity |
| GO:0140640 | 5 | 0.976 | catalytic activity, acting on a nucleic acid |
| GO:0016671 | 4 | 7.96E-03 | oxidoreductase, acting on sulfur donors, disulfide acceptors |
| GO:0008238 | 4 | 0.0445 | exopeptidase activity |
| GO:0008237 | 4 | 0.153 | metallopeptidase activity |
| GO:0050660 | 4 | 0.209 | flavin adenine dinucleotide binding |
| GO:0005216 | 4 | 0.871 | monoatomic ion channel activity |
| GO:0016853 | 4 | 0.881 | isomerase activity |
| GO:0015075 | 4 | 0.969 | monoatomic ion transmembrane transporter activity |
| GO:0008235 | 3 | 0.0218 | metalloexopeptidase activity |
| GO:0008483 | 3 | 0.0218 | transaminase activity |
| GO:0016769 | 3 | 0.0218 | transferase activity, transferring nitrogenous groups |
| GO:0004180 | 3 | 0.0285 | carboxypeptidase activity |
| GO:0016627 | 3 | 0.181 | oxidoreductase activity, acting on the CH-CH group of donors |
| GO:0016758 | 3 | 0.237 | hexosyltransferase activity |
| GO:0016810 | 3 | 0.254 | hydrolase, acting on carbon-nitrogen (not peptide) bonds |
| GO:0008080 | 3 | 0.313 | N-acetyltransferase activity |
| GO:0016410 | 3 | 0.325 | N-acyltransferase activity |
| GO:0016407 | 3 | 0.39 | acetyltransferase activity |
| GO:0016859 | 3 | 0.623 | cis-trans isomerase activity |
| GO:0003755 | 3 | 0.623 | peptidyl-prolyl cis-trans isomerase activity |
| GO:0140097 | 3 | 0.679 | catalytic activity, acting on DNA |
| GO:0015318 | 3 | 0.911 | inorganic molecular entity transmembrane transporter activity |
| GO:0043531 | 2 | 1.37E-03 | ADP binding |
| GO:0008476 | 2 | 1.37E-03 | protein-tyrosine sulfotransferase activity |
| GO:0046559 | 2 | 2.70E-03 | alpha-glucuronidase activity |
| GO:0004364 | 2 | 0.0224 | glutathione transferase activity |
| GO:0004181 | 2 | 0.0265 | metallocarboxypeptidase activity |
| GO:0004650 | 2 | 0.0309 | polygalacturonase activity |
| GO:0004930 | 2 | 0.0309 | G protein-coupled receptor activity |
| GO:0033743 | 2 | 0.0404 | peptide-methionine (R)-S-oxide reductase activity |
| GO:0008113 | 2 | 0.0682 | peptide-methionine (S)-S-oxide reductase activity |
| GO:0003690 | 2 | 0.0871 | double-stranded DNA binding |
| GO:0000030 | 2 | 0.1 | mannosyltransferase activity |
| GO:0005254 | 2 | 0.182 | chloride channel activity |
| GO:0052689 | 2 | 0.189 | carboxylic ester hydrolase activity |
| GO:0005253 | 2 | 0.189 | monoatomic anion channel activity |
| GO:0015108 | 2 | 0.197 | chloride transmembrane transporter activity |

| GO Term | Count | p-value | Name |
|---|---|---|---|
| GO:0008509 | 2 | 0.221 | monoatomic anion transmembrane transporter activity |
| GO:0016835 | 2 | 0.229 | carbon-oxygen lyase activity |
| GO:0050661 | 2 | 0.237 | NADP binding |
| GO:0004888 | 2 | 0.269 | transmembrane signaling receptor activity |
| GO:0016765 | 2 | 0.269 | transferase, transferring alkyl or aryl (other than methyl) groups |
| GO:0038023 | 2 | 0.309 | signaling receptor activity |
| GO:0060089 | 2 | 0.309 | molecular transducer activity |
| GO:0008170 | 2 | 0.309 | N-methyltransferase activity |
| GO:0015103 | 2 | 0.469 | inorganic anion transmembrane transporter activity |
| GO:0008173 | 2 | 0.483 | RNA methyltransferase activity |
| GO:0051536 | 2 | 0.66 | iron-sulfur cluster binding |
| GO:0051540 | 2 | 0.66 | metal cluster binding |
| GO:0016874 | 2 | 0.821 | ligase activity |
| GO:0140098 | 2 | 0.982 | catalytic activity, acting on RNA |
| GO:0003723 | 2 | 0.987 | RNA binding |
| GO:0050080 | 1 | 0.0216 | malonyl-CoA decarboxylase activity |
| GO:0030570 | 1 | 0.0216 | pectate lyase activity |
| GO:0033735 | 1 | 0.0216 | aspartate dehydrogenase activity |
| GO:0008988 | 1 | 0.0216 | rRNA (adenine-N6-)-methyltransferase activity |
| GO:0047304 | 1 | 0.0216 | 2-aminoethylphosphonate-pyruvate transaminase activity |
| GO:0016837 | 1 | 0.0216 | carbon-oxygen lyase activity, acting on polysaccharides |
| GO:0004668 | 1 | 0.0216 | protein-arginine deiminase activity |
| GO:0004096 | 1 | 0.0427 | catalase activity |
| GO:0004852 | 1 | 0.0427 | uroporphyrinogen-III synthase activity |
| GO:0000150 | 1 | 0.0427 | DNA strand exchange activity |
| GO:0004013 | 1 | 0.0427 | adenosylhomocysteinase activity |
| GO:0008716 | 1 | 0.0427 | D-alanine-D-alanine ligase activity |
| GO:0046508 | 1 | 0.0427 | hydrolase activity, acting on carbon-sulfur bonds |
| GO:0033699 | 1 | 0.0427 | DNA 5'-adenosine monophosphate hydrolase activity |

**Supplementary Table 4 Molecular function-related GO term enrichment results for LGTs specific to *Au. lagunensis* (see associated REVIGO plot in Figure 3C).** Only terms that were found to be significantly enriched (p-value <0.05) that occurred >1 time are included.

| GO Term | Count | p-value | Name |
|---|---|---|---|
| GO:0003674 | 56 | 1 | molecular function |
| GO:0003824 | 40 | 1.35E-03 | catalytic activity |
| GO:0005488 | 23 | 1 | binding |
| GO:0016787 | 15 | 0.0179 | hydrolase activity |
| GO:0036094 | 15 | 0.682 | small molecule binding |
| GO:0043167 | 12 | 0.863 | ion binding |
| GO:0016740 | 10 | 0.607 | transferase activity |
| GO:0097159 | 10 | 0.944 | organic cyclic compound binding |
| GO:1901363 | 9 | 0.605 | heterocyclic compound binding |
| GO:0016491 | 8 | 0.199 | oxidoreductase activity |
| GO:0030554 | 8 | 0.32 | adenyl nucleotide binding |
| GO:0140096 | 8 | 0.415 | catalytic activity, acting on a protein |
| GO:0017076 | 8 | 0.457 | purine nucleotide binding |
| GO:1901265 | 8 | 0.571 | nucleoside phosphate binding |
| GO:0000166 | 8 | 0.571 | nucleotide binding |
| GO:0005515 | 7 | 1 | protein binding |
| GO:0046872 | 6 | 0.787 | metal ion binding |
| GO:0043169 | 6 | 0.801 | cation binding |
| GO:0043168 | 6 | 0.897 | anion binding |
| GO:0016788 | 5 | 0.0812 | hydrolase activity, acting on ester bonds |
| GO:0005524 | 5 | 0.722 | ATP binding |
| GO:0032559 | 5 | 0.732 | adenyl ribonucleotide binding |
| GO:0035639 | 5 | 0.825 | purine ribonucleoside triphosphate binding |
| GO:0032555 | 5 | 0.832 | purine ribonucleotide binding |
| GO:0032553 | 5 | 0.844 | ribonucleotide binding |
| GO:0097367 | 5 | 0.85 | carbohydrate derivative binding |
| GO:0004553 | 4 | 7.35E-03 | hydrolase activity, hydrolyzing O-glycosyl compounds |
| GO:0016798 | 4 | 0.0146 | hydrolase activity, acting on glycosyl bonds |
| GO:0022857 | 4 | 0.553 | transmembrane transporter activity |
| GO:0005215 | 4 | 0.574 | transporter activity |
| GO:0016614 | 3 | 0.0374 | oxidoreductase activity, acting on CH-OH group of donors |
| GO:0042578 | 3 | 0.0764 | phosphoric ester hydrolase activity |
| GO:0016462 | 3 | 0.228 | pyrophosphatase activity |
| GO:0016818 | 3 | 0.241 | hydrolase, acting on phosphorus-containing aacid nhydrides |

| GO Term | Count | p-value | Name |
|---|---|---|---|
| GO:0016817 | 3 | 0.242 | hydrolase activity, acting on acid anhydrides |
| GO:0008233 | 3 | 0.383 | peptidase activity |
| GO:0016773 | 3 | 0.423 | phosphotransferase activity, alcohol group as acceptor |
| GO:0016301 | 3 | 0.491 | kinase activity |
| GO:0016772 | 3 | 0.635 | transferase activity, transferring phosphorus-containing groups |
| GO:0008081 | 2 | 8.71E-03 | phosphoric diester hydrolase activity |
| GO:0016903 | 2 | 0.0149 | oxidoreductase, acting on the aldehyde or oxo group of donors |
| GO:0000287 | 2 | 0.0198 | magnesium ion binding |
| GO:0051287 | 2 | 0.0368 | NAD binding |
| GO:0016829 | 2 | 0.13 | lyase activity |
| GO:0016616 | 2 | 0.136 | oxidoreductase, acting on CH-OH donors, NAD acceptors |
| GO:0016874 | 2 | 0.164 | ligase activity |
| GO:0016887 | 2 | 0.206 | ATP hydrolysis activity |
| GO:0016747 | 2 | 0.334 | acyltransferase, transferring groups other than amino-acyl |
| GO:0016757 | 2 | 0.415 | glycosyltransferase activity |
| GO:0017111 | 2 | 0.452 | ribonucleoside triphosphate phosphatase activity |
| GO:0140657 | 2 | 0.575 | ATP-dependent activity |
| GO:0016741 | 2 | 0.581 | transferase activity, transferring one-carbon groups |
| GO:0004672 | 2 | 0.616 | protein kinase activity |
| GO:0016746 | 2 | 0.691 | acyltransferase activity |
| GO:0003842 | 1 | 5.05E-03 | 1-pyrroline-5-carboxylate dehydrogenase activity |
| GO:0003908 | 1 | 5.05E-03 | methylated-DNA-[protein]-cysteine S-methyltransferase activity |
| GO:0004856 | 1 | 5.05E-03 | D-xylulokinase activity |
| GO:0050532 | 1 | 5.05E-03 | 2-phosphosulfolactate phosphatase activity |
| GO:0030248 | 1 | 0.0101 | cellulose binding |
| GO:0004609 | 1 | 0.0101 | phosphatidylserine decarboxylase activity |
| GO:0003796 | 1 | 0.0101 | lysozyme activity |
| GO:0061783 | 1 | 0.0101 | peptidoglycan muralytic activity |
| GO:0008172 | 1 | 0.0101 | S-methyltransferase activity |
| GO:0008239 | 1 | 0.0151 | dipeptidyl-peptidase activity |
| GO:0030247 | 1 | 0.0151 | polysaccharide binding |
| GO:0004657 | 1 | 0.0151 | proline dehydrogenase activity |
| GO:0016649 | 1 | 0.0201 | oxidoreductase, acting on the CH-NH group of donors, quinone or similar compound as acceptor |
| GO:0004470 | 1 | 0.0201 | malic enzyme activity |
| GO:0004471 | 1 | 0.0201 | malate dehydrogenase (decarboxylating) (NAD+) activity |
| GO:0030600 | 1 | 0.025 | feruloyl esterase activity |
| GO:0016615 | 1 | 0.025 | malate dehydrogenase activity |
| GO:0000774 | 1 | 0.025 | adenyl-nucleotide exchange factor activity |

| GO Term | Count | p-value | Name |
|---|---|---|---|
| GO:0042803 | 1 | 0.0299 | protein homodimerization activity |
| GO:0008864 | 1 | 0.0299 | formyltetrahydrofolate deformylase activity |
| GO:0004427 | 1 | 0.0446 | inorganic diphosphate phosphatase activity |
| GO:0004356 | 1 | 0.0446 | glutamine synthetase activity |
| GO:0042802 | 1 | 0.0494 | identical protein binding |
| GO:0016211 | 1 | 0.0494 | ammonia ligase activity |
| GO:0008417 | 1 | 0.0494 | fucosyltransferase activity |
| GO:0016880 | 1 | 0.0494 | acid-ammonia (or amide) ligase activity |
| GO:0004112 | 1 | 0.0494 | cyclic-nucleotide phosphodiesterase activity |

**Supplementary Table 5 Biological process-related GO term enrichment results for the *Ac. anophagefferens* accessory genome (see associated REVIGO plot in Extended Data Figure 6).** Only terms that were found to be significantly enriched (p-value <0.05) or highly abundant (i.e., within the top 50 occurring GO terms) are included.

| GO Term | Count | p-value | Name |
|---|---|---|---|
| GO:0008150 | 447 | 1 | biological process |
| GO:0008152 | 305 | 0.216 | metabolic process |
| GO:0071704 | 281 | 0.247 | organic substance metabolic process |
| GO:0009987 | 270 | 0.512 | cellular process |
| GO:0044238 | 267 | 0.0979 | primary metabolic process |
| GO:0043170 | 192 | 0.115 | macromolecule metabolic process |
| GO:1901564 | 152 | 0.687 | organonitrogen compound metabolic process |
| GO:0044237 | 149 | 0.0194 | cellular metabolic process |
| GO:0019538 | 119 | 0.405 | protein metabolic process |
| GO:1901360 | 102 | 0.107 | organic cyclic compound metabolic process |
| GO:0006139 | 93 | 0.0697 | nucleobase-containing compound metabolic process |
| GO:0009058 | 74 | 0.189 | biosynthetic process |
| GO:1901576 | 72 | 0.0658 | organic substance biosynthetic process |
| GO:0043412 | 72 | 0.923 | macromolecule modification |
| GO:0090304 | 70 | 0.138 | nucleic acid metabolic process |
| GO:0044249 | 66 | 2.64E-04 | cellular biosynthetic process |
| GO:0051179 | 64 | 0.998 | localization |
| GO:0036211 | 63 | 0.902 | protein modification process |
| GO:0051234 | 62 | 0.997 | establishment of localization |
| GO:0006810 | 61 | 0.997 | transport |
| GO:0006793 | 56 | 0.813 | phosphorus metabolic process |
| GO:0006796 | 55 | 0.838 | phosphate-containing compound metabolic process |
| GO:0055085 | 51 | 0.811 | transmembrane transport |
| GO:0044281 | 49 | 0.0998 | small molecule metabolic process |
| GO:1901566 | 43 | 0.0571 | organonitrogen compound biosynthetic process |
| GO:0016070 | 42 | 0.552 | RNA metabolic process |
| GO:0009059 | 41 | 9.14E-05 | macromolecule biosynthetic process |
| GO:0065007 | 36 | 0.797 | biological regulation |
| GO:0050789 | 35 | 0.746 | regulation of biological process |
| GO:0006629 | 33 | 0.219 | lipid metabolic process |
| GO:0050794 | 33 | 0.784 | regulation of cellular process |
| GO:0006412 | 31 | 5.34E-06 | translation |
| GO:0019752 | 29 | 0.0555 | carboxylic acid metabolic process |

| GO Term | Count | p-value | Name |
|---|---|---|---|
| GO:0043436 | 29 | 0.0608 | oxoacid metabolic process |
| GO:0006082 | 29 | 0.0695 | organic acid metabolic process |
| GO:0006259 | 28 | 0.04 | DNA metabolic process |
| GO:0080090 | 27 | 0.0572 | regulation of primary metabolic process |
| GO:0060255 | 27 | 0.142 | regulation of macromolecule metabolic process |
| GO:0019222 | 27 | 0.174 | regulation of metabolic process |
| GO:0016043 | 27 | 0.573 | cellular component organization |
| GO:0071840 | 27 | 0.686 | cellular component organization or biogenesis |
| GO:0016310 | 27 | 0.851 | phosphorylation |
| GO:0005975 | 26 | 0.61 | carbohydrate metabolic process |
| GO:0019219 | 26 | 0.0273 | regulation of nucleobase-containing compound metabolic process |
| GO:0009889 | 26 | 0.0915 | regulation of biosynthetic process |
| GO:0010556 | 26 | 0.0915 | regulation of macromolecule biosynthetic process |
| GO:0031326 | 26 | 0.0915 | regulation of cellular biosynthetic process |
| GO:0010468 | 26 | 0.0915 | regulation of gene expression |
| GO:0031323 | 26 | 0.157 | regulation of cellular metabolic process |
| GO:0006468 | 26 | 0.858 | protein phosphorylation |
| GO:0051252 | 25 | 0.0344 | regulation of RNA metabolic process |
| GO:0006355 | 25 | 0.0344 | regulation of DNA-templated transcription |
| GO:2001141 | 25 | 0.0344 | regulation of RNA biosynthetic process |
| GO:0032787 | 18 | 8.14E-03 | monocarboxylic acid metabolic process |
| GO:0006631 | 13 | 1.47E-04 | fatty acid metabolic process |
| GO:0072330 | 11 | 2.61E-03 | monocarboxylic acid biosynthetic process |
| GO:0006633 | 11 | 5.96E-04 | fatty acid biosynthetic process |
| GO:0006979 | 7 | 0.012 | response to oxidative stress |
| GO:0015074 | 5 | 6.68E-03 | DNA integration |
| GO:0006261 | 3 | 0.021 | DNA-templated DNA replication |
| GO:0033151 | 2 | 8.62E-03 | V(D)J recombination |
| GO:0002562 | 2 | 8.62E-03 | somatic diversification of immune receptors |
| GO:0016444 | 2 | 8.62E-03 | somatic cell DNA recombination |
| GO:0002200 | 2 | 8.62E-03 | somatic diversification of immune receptors |
| GO:0045337 | 2 | 0.0243 | farnesyl diphosphate biosynthetic process |
| GO:0045338 | 2 | 0.0243 | farnesyl diphosphate metabolic process |
| GO:0010142 | 2 | 0.0243 | farnesyl diphosphate biosynthetic process, mevalonate pathway |
| GO:0002376 | 2 | 0.0456 | immune system process |
| GO:0070070 | 2 | 0.0456 | proton-transporting V-type ATPase complex assembly |
| GO:0070072 | 2 | 0.0456 | vacuolar proton-transporting V-type ATPase complex assembly |
| GO:0006545 | 2 | 0.0456 | glycine biosynthetic process |

**Supplementary Table 6 Molecular function-related GO term enrichment results for the *Ac. anophagefferens* accessory genome (see associated REVIGO plot in Extended Data Figure 6).** Only terms that were found to be significantly enriched (p-value <0.05) or highly abundant (i.e., within the top 50 occurring GO terms) are included.

| GO Term | Count | p-value | Name |
|---|---|---|---|
| GO:0003674 | 1075 | 1 | molecular function |
| GO:0005488 | 686 | 0.239 | binding |
| GO:0003824 | 500 | 1 | catalytic activity |
| GO:0005515 | 354 | 0.0297 | protein binding |
| GO:0036094 | 264 | 1 | small molecule binding |
| GO:0097159 | 255 | 0.989 | organic cyclic compound binding |
| GO:0043167 | 229 | 1 | ion binding |
| GO:0016740 | 202 | 0.412 | transferase activity |
| GO:1901363 | 147 | 1 | heterocyclic compound binding |
| GO:0016787 | 137 | 0.998 | hydrolase activity |
| GO:0043168 | 137 | 1 | anion binding |
| GO:0003676 | 122 | 0.337 | nucleic acid binding |
| GO:1901265 | 120 | 1 | nucleoside phosphate binding |
| GO:0000166 | 120 | 1 | nucleotide binding |
| GO:0043169 | 116 | 0.995 | cation binding |
| GO:0046872 | 115 | 0.993 | metal ion binding |
| GO:0017076 | 111 | 1 | purine nucleotide binding |
| GO:0097367 | 106 | 1 | carbohydrate derivative binding |
| GO:0032553 | 104 | 1 | ribonucleotide binding |
| GO:0032555 | 102 | 1 | purine ribonucleotide binding |
| GO:0035639 | 100 | 1 | purine ribonucleoside triphosphate binding |
| GO:0140096 | 98 | 1 | catalytic activity, acting on a protein |
| GO:0030554 | 96 | 1 | adenyl nucleotide binding |
| GO:0016491 | 93 | 0.974 | oxidoreductase activity |
| GO:0032559 | 87 | 1 | adenyl ribonucleotide binding |
| GO:0005524 | 85 | 1 | ATP binding |
| GO:0016746 | 69 | 1.84E-04 | acyltransferase activity |
| GO:0046914 | 53 | 0.685 | transition metal ion binding |
| GO:0003677 | 52 | 0.0445 | DNA binding |
| GO:0022857 | 52 | 0.999 | transmembrane transporter activity |
| GO:0005215 | 52 | 1 | transporter activity |
| GO:0019842 | 45 | 7.32E-03 | vitamin binding |
| GO:0016772 | 45 | 0.997 | transferase activity, transferring phosphorus-containing groups |

| GO Term | Count | p-value | Name |
| --- | --- | --- | --- |
| GO:0005509 | 41 | 0.994 | calcium ion binding |
| GO:0016741 | 40 | 0.332 | transferase activity, transferring one-carbon groups |
| GO:0008168 | 38 | 0.356 | methyltransferase activity |
| GO:0016301 | 36 | 0.993 | kinase activity |
| GO:0140640 | 35 | 0.984 | catalytic activity, acting on a nucleic acid |
| GO:0016788 | 34 | 0.982 | hydrolase activity, acting on ester bonds |
| GO:0005198 | 32 | 1.98E-03 | structural molecule activity |
| GO:0008233 | 32 | 0.978 | peptidase activity |
| GO:0003735 | 31 | 2.39E-04 | structural constituent of ribosome |
| GO:0140657 | 29 | 0.957 | ATP-dependent activity |
| GO:0016773 | 29 | 0.998 | phosphotransferase activity, alcohol group as acceptor |
| GO:0016747 | 28 | 0.152 | acyltransferase, transferring groups other than amino-acyl |
| GO:0003723 | 28 | 0.66 | RNA binding |
| GO:0004672 | 28 | 0.983 | protein kinase activity |
| GO:0016818 | 27 | 0.932 | hydrolase, acting on phosphorus-containing acid anhydrides |
| GO:0016817 | 27 | 0.934 | hydrolase activity, acting on acid anhydrides |
| GO:0015075 | 27 | 0.991 | monoatomic ion transmembrane transporter activity |
| GO:0033218 | 20 | 1.61E-07 | amide binding |
| GO:0140110 | 20 | 0.0408 | transcription regulator activity |
| GO:0031177 | 19 | 6.47E-09 | phosphopantetheine binding |
| GO:0072341 | 19 | 1.10E-08 | modified amino acid binding |
| GO:0003700 | 19 | 0.0249 | DNA-binding transcription factor activity |
| GO:0046983 | 12 | 0.0178 | protein dimerization activity |
| GO:0016209 | 11 | 0.0282 | antioxidant activity |
| GO:0046982 | 10 | 3.33E-03 | protein heterodimerization activity |
| GO:0004315 | 9 | 5.40E-04 | 3-oxoacyl-[acyl-carrier-protein] synthase activity |
| GO:0005230 | 8 | 0.0189 | extracellular ligand-gated monoatomic ion channel activity |
| GO:0004601 | 7 | 0.0177 | peroxidase activity |
| GO:0016684 | 7 | 0.0177 | oxidoreductase activity, acting on peroxide as acceptor |
| GO:0008569 | 6 | 0.0114 | minus-end-directed microtubule motor activity |
| GO:0016846 | 5 | 2.81E-03 | carbon-sulfur lyase activity |
| GO:0045505 | 5 | 0.0207 | dynein intermediate chain binding |
| GO:0051959 | 5 | 0.0207 | dynein light intermediate chain binding |
| GO:0004602 | 4 | 0.0278 | glutathione peroxidase activity |
| GO:0004127 | 3 | 0.012 | cytidylate kinase activity |
| GO:0004844 | 3 | 0.0221 | uracil DNA N-glycosylase activity |
| GO:0008121 | 2 | 0.0127 | ubiquinol-cytochrome-c reductase activity |
| GO:0004421 | 2 | 0.0353 | hydroxymethylglutaryl-CoA synthase activity |
| GO:0004683 | 2 | 0.0353 | calmodulin-dependent protein kinase activity |

**Supplemental Table 7 Biological process-related GO term enrichment results for LGTs in the *Ac. anophagefferens* accessory genome (see associated REVIGO plot in Extended Data Figure 7).**

| GO Term | Count | p-value | name |
|---|---|---|---|
| GO:0008150 | 25 | 1 | biological process |
| GO:0008152 | 21 | 0.0437 | metabolic process |
| GO:0044238 | 18 | 0.088 | primary metabolic process |
| GO:0071704 | 18 | 0.185 | organic substance metabolic process |
| GO:0009987 | 11 | 0.969 | cellular process |
| GO:0043170 | 9 | 0.733 | macromolecule metabolic process |
| GO:0044237 | 8 | 0.442 | cellular metabolic process |
| GO:0006139 | 7 | 0.152 | nucleobase-containing compound metabolic process |
| GO:1901360 | 7 | 0.237 | organic cyclic compound metabolic process |
| GO:1901564 | 7 | 0.826 | organonitrogen compound metabolic process |
| GO:0009058 | 6 | 0.162 | biosynthetic process |
| GO:0006629 | 5 | 0.0194 | lipid metabolic process |
| GO:1901576 | 5 | 0.248 | organic substance biosynthetic process |
| GO:0090304 | 5 | 0.259 | nucleic acid metabolic process |
| GO:0019538 | 4 | 0.922 | protein metabolic process |
| GO:0006281 | 3 | 0.0335 | DNA repair |
| GO:0006974 | 3 | 0.0387 | DNA damage response |
| GO:0051716 | 3 | 0.0486 | cellular response to stimulus |
| GO:0033554 | 3 | 0.0486 | cellular response to stress |
| GO:0006950 | 3 | 0.0719 | response to stress |
| GO:0050896 | 3 | 0.0892 | response to stimulus |
| GO:0006259 | 3 | 0.0984 | DNA metabolic process |
| GO:0005975 | 3 | 0.189 | carbohydrate metabolic process |
| GO:1901566 | 3 | 0.294 | organonitrogen compound biosynthetic process |
| GO:0044281 | 3 | 0.406 | small molecule metabolic process |
| GO:0044249 | 3 | 0.448 | cellular biosynthetic process |
| GO:0001522 | 2 | 0.0215 | pseudouridine synthesis |
| GO:0016052 | 2 | 0.0337 | carbohydrate catabolic process |
| GO:0009150 | 2 | 0.0742 | purine ribonucleotide metabolic process |
| GO:0019693 | 2 | 0.0817 | ribose phosphate metabolic process |
| GO:0032787 | 2 | 0.104 | monocarboxylic acid metabolic process |
| GO:0006163 | 2 | 0.112 | purine nucleotide metabolic process |
| GO:0009451 | 2 | 0.117 | RNA modification |
| GO:0072521 | 2 | 0.123 | purine-containing compound metabolic process |
| GO:0006412 | 2 | 0.171 | translation |

| GO Term | Count | p-value | name |
|---|---|---|---|
| GO:0009117 | 2 | 0.177 | nucleotide metabolic process |
| GO:0006753 | 2 | 0.184 | nucleoside phosphate metabolic process |
| GO:1901135 | 2 | 0.23 | carbohydrate derivative metabolic process |
| GO:0044255 | 2 | 0.232 | cellular lipid metabolic process |
| GO:0055086 | 2 | 0.238 | nucleobase-containing small molecule metabolic process |
| GO:1901575 | 2 | 0.322 | organic substance catabolic process |
| GO:0009056 | 2 | 0.336 | catabolic process |
| GO:0019752 | 2 | 0.339 | carboxylic acid metabolic process |
| GO:0043436 | 2 | 0.343 | oxoacid metabolic process |
| GO:0006082 | 2 | 0.349 | organic acid metabolic process |
| GO:0019637 | 2 | 0.359 | organophosphate metabolic process |
| GO:0009059 | 2 | 0.363 | macromolecule biosynthetic process |
| GO:0006508 | 2 | 0.545 | proteolysis |
| GO:0016070 | 2 | 0.701 | RNA metabolic process |
| GO:0006796 | 2 | 0.876 | phosphate-containing compound metabolic process |
| GO:0006793 | 2 | 0.879 | phosphorus metabolic process |
| GO:0043412 | 2 | 0.96 | macromolecule modification |
| GO:0071722 | 1 | 5.20E-03 | detoxification of arsenic-containing substance |
| GO:0098754 | 1 | 0.0206 | detoxification |
| GO:0006672 | 1 | 0.0358 | ceramide metabolic process |
| GO:0006189 | 1 | 0.0358 | 'de novo' IMP biosynthetic process |
| GO:0045493 | 1 | 0.0459 | xylan catabolic process |
| GO:0044347 | 1 | 0.0459 | cell wall polysaccharide catabolic process |
| GO:0016998 | 1 | 0.0459 | cell wall macromolecule catabolic process |
| GO:2000895 | 1 | 0.0459 | hemicellulose catabolic process |
| GO:0006665 | 1 | 0.0459 | sphingolipid metabolic process |
| GO:0045491 | 1 | 0.0508 | xylan metabolic process |
| GO:0006188 | 1 | 0.0508 | IMP biosynthetic process |
| GO:0006298 | 1 | 0.0508 | mismatch repair |
| GO:0010410 | 1 | 0.0508 | hemicellulose metabolic process |
| GO:0044036 | 1 | 0.0508 | cell wall macromolecule metabolic process |
| GO:0010383 | 1 | 0.0508 | cell wall polysaccharide metabolic process |
| GO:0009168 | 1 | 0.0558 | purine ribonucleoside monophosphate biosynthetic process |
| GO:0008272 | 1 | 0.0607 | sulfate transport |
| GO:0009123 | 1 | 0.0656 | nucleoside monophosphate metabolic process |
| GO:0009124 | 1 | 0.0656 | nucleoside monophosphate biosynthetic process |
| GO:0000272 | 1 | 0.0704 | polysaccharide catabolic process |
| GO:0006289 | 1 | 0.0753 | nucleotide-excision repair |
| GO:0016051 | 1 | 0.113 | carbohydrate biosynthetic process |

| GO Term | Count | p-value | name |
|---|---|---|---|
| GO:0006643 | 1 | 0.118 | membrane lipid metabolic process |
| GO:0005976 | 1 | 0.132 | polysaccharide metabolic process |
| GO:0009152 | 1 | 0.15 | purine ribonucleotide biosynthetic process |
| GO:0009260 | 1 | 0.158 | ribonucleotide biosynthetic process |
| GO:0046390 | 1 | 0.158 | ribose phosphate biosynthetic process |
| GO:0015698 | 1 | 0.176 | inorganic anion transport |
| GO:0006633 | 1 | 0.185 | fatty acid biosynthetic process |
| GO:0072526 | 1 | 0.193 | pyridine-containing compound catabolic process |
| GO:0009261 | 1 | 0.193 | ribonucleotide catabolic process |
| GO:0009179 | 1 | 0.193 | purine ribonucleoside diphosphate metabolic process |
| GO:0009181 | 1 | 0.193 | purine ribonucleoside diphosphate catabolic process |
| GO:0009185 | 1 | 0.193 | ribonucleoside diphosphate metabolic process |
| GO:0009134 | 1 | 0.193 | nucleoside diphosphate catabolic process |
| GO:0009137 | 1 | 0.193 | purine nucleoside diphosphate catabolic process |
| GO:0009135 | 1 | 0.193 | purine nucleoside diphosphate metabolic process |
| GO:0046032 | 1 | 0.193 | ADP catabolic process |
| GO:0046031 | 1 | 0.193 | ADP metabolic process |
| GO:0019364 | 1 | 0.193 | pyridine nucleotide catabolic process |
| GO:0009191 | 1 | 0.193 | ribonucleoside diphosphate catabolic process |
| GO:0006096 | 1 | 0.193 | glycolytic process |
| GO:0009154 | 1 | 0.193 | purine ribonucleotide catabolic process |
| GO:0006195 | 1 | 0.193 | purine nucleotide catabolic process |
| GO:0072523 | 1 | 0.197 | purine-containing compound catabolic process |
| GO:0009132 | 1 | 0.197 | nucleoside diphosphate metabolic process |
| GO:0006164 | 1 | 0.202 | purine nucleotide biosynthetic process |
| GO:1901136 | 1 | 0.206 | carbohydrate derivative catabolic process |
| GO:0006631 | 1 | 0.21 | fatty acid metabolic process |
| GO:0072330 | 1 | 0.214 | monocarboxylic acid biosynthetic process |
| GO:0072522 | 1 | 0.218 | purine-containing compound biosynthetic process |
| GO:0006090 | 1 | 0.218 | pyruvate metabolic process |
| GO:0046034 | 1 | 0.234 | ATP metabolic process |
| GO:0009205 | 1 | 0.246 | purine ribonucleoside triphosphate metabolic process |
| GO:1901292 | 1 | 0.258 | nucleoside phosphate catabolic process |
| GO:0046496 | 1 | 0.27 | nicotinamide nucleotide metabolic process |
| GO:0019362 | 1 | 0.27 | pyridine nucleotide metabolic process |
| GO:0009141 | 1 | 0.274 | nucleoside triphosphate metabolic process |
| GO:0046434 | 1 | 0.274 | organophosphate catabolic process |
| GO:0072524 | 1 | 0.289 | pyridine-containing compound metabolic process |
| GO:0009165 | 1 | 0.289 | nucleotide biosynthetic process |

| GO Term | Count | p-value | name |
|---|---|---|---|
| GO:1901293 | 1 | 0.289 | nucleoside phosphate biosynthetic process |
| GO:1901137 | 1 | 0.307 | carbohydrate derivative biosynthetic process |
| GO:0034655 | 1 | 0.34 | nucleobase-containing compound catabolic process |
| GO:1901361 | 1 | 0.374 | organic cyclic compound catabolic process |
| GO:1901565 | 1 | 0.393 | organonitrogen compound catabolic process |
| GO:0032259 | 1 | 0.406 | methylation |
| GO:0043603 | 1 | 0.44 | amide metabolic process |
| GO:0009057 | 1 | 0.44 | macromolecule catabolic process |
| GO:0006457 | 1 | 0.446 | protein folding |
| GO:0090407 | 1 | 0.458 | organophosphate biosynthetic process |
| GO:0006091 | 1 | 0.466 | generation of precursor metabolites and energy |
| GO:0016053 | 1 | 0.48 | organic acid biosynthetic process |
| GO:0046394 | 1 | 0.48 | carboxylic acid biosynthetic process |
| GO:0034654 | 1 | 0.541 | nucleobase-containing compound biosynthetic process |
| GO:0008610 | 1 | 0.543 | lipid biosynthetic process |
| GO:0044283 | 1 | 0.576 | small molecule biosynthetic process |
| GO:0006520 | 1 | 0.62 | amino acid metabolic process |
| GO:2001141 | 1 | 0.626 | regulation of RNA biosynthetic process |
| GO:0006355 | 1 | 0.626 | regulation of DNA-templated transcription |
| GO:0010556 | 1 | 0.68 | regulation of macromolecule biosynthetic process |
| GO:0031326 | 1 | 0.68 | regulation of cellular biosynthetic process |
| GO:0010468 | 1 | 0.68 | regulation of gene expression |
| GO:0009889 | 1 | 0.68 | regulation of biosynthetic process |
| GO:0031323 | 1 | 0.704 | regulation of cellular metabolic process |
| GO:1901362 | 1 | 0.723 | organic cyclic compound biosynthetic process |
| GO:0065007 | 1 | 0.907 | biological regulation |
| GO:0006810 | 1 | 0.994 | transport |
| GO:0051179 | 1 | 0.995 | localization |

**Supplemental Table 8 Molecular function-related GO term enrichment results for LGTs in the *Ac. anophagefferens* accessory genome (see associated REVIGO plot in Extended Data Figure 7)**

| GO Term | Count | p-value | Name |
|---|---|---|---|
| GO:0003674 | 55 | 1 | molecular function |
| GO:0003824 | 37 | 0.0127 | catalytic activity |
| GO:0005488 | 25 | 0.997 | binding |
| GO:0036094 | 17 | 0.424 | small molecule binding |
| GO:0097159 | 15 | 0.503 | organic cyclic compound binding |
| GO:0016740 | 13 | 0.206 | transferase activity |
| GO:0043167 | 12 | 0.847 | ion binding |
| GO:0016787 | 11 | 0.228 | hydrolase activity |
| GO:0003676 | 9 | 0.141 | nucleic acid binding |
| GO:1901363 | 9 | 0.633 | heterocyclic compound binding |
| GO:0016491 | 8 | 0.199 | oxidoreductase activity |
| GO:1901265 | 7 | 0.73 | nucleoside phosphate binding |
| GO:0000166 | 7 | 0.73 | nucleotide binding |
| GO:0043168 | 7 | 0.821 | anion binding |
| GO:0030554 | 6 | 0.657 | adenyl nucleotide binding |
| GO:0046872 | 6 | 0.736 | metal ion binding |
| GO:0043169 | 6 | 0.753 | cation binding |
| GO:0017076 | 6 | 0.775 | purine nucleotide binding |
| GO:0016741 | 5 | 0.0406 | transferase activity, transferring one-carbon groups |
| GO:0003677 | 5 | 0.0587 | DNA binding |
| GO:0032559 | 5 | 0.744 | adenyl ribonucleotide binding |
| GO:0032555 | 5 | 0.843 | purine ribonucleotide binding |
| GO:0032553 | 5 | 0.853 | ribonucleotide binding |
| GO:0097367 | 5 | 0.858 | carbohydrate derivative binding |
| GO:0019842 | 4 | 0.0754 | vitamin binding |
| GO:0008168 | 4 | 0.108 | methyltransferase activity |
| GO:0005509 | 4 | 0.337 | calcium ion binding |
| GO:0005524 | 4 | 0.867 | ATP binding |
| GO:0035639 | 4 | 0.927 | purine ribonucleoside triphosphate binding |
| GO:0016616 | 3 | 0.0267 | oxidoreductase activity, acting on the CH-OH group of donors |
| GO:0016614 | 3 | 0.0361 | oxidoreductase activity, acting on CH-OH group of donors |
| GO:0008146 | 3 | 0.061 | sulfotransferase activity |
| GO:0016782 | 3 | 0.0758 | transferase activity, transferring sulphur-containing groups |
| GO:0016853 | 3 | 0.189 | isomerase activity |
| GO:0003723 | 3 | 0.194 | RNA binding |

| GO Term | Count | p-value | Name |
|---|---|---|---|
| GO:0016746 | 3 | 0.409 | acyltransferase activity |
| GO:0016788 | 3 | 0.423 | hydrolase activity, acting on ester bonds |
| GO:0140096 | 3 | 0.977 | catalytic activity, acting on a protein |
| GO:0005515 | 3 | 1 | protein binding |
| GO:0016811 | 2 | 0.0137 | hydrolase, acting on C-N (non-peptide) bonds, in linear amides |
| GO:0031177 | 2 | 0.0212 | phosphopantetheine binding |
| GO:0072341 | 2 | 0.0222 | modified amino acid binding |
| GO:0009982 | 2 | 0.0243 | pseudouridine synthase activity |
| GO:0033218 | 2 | 0.0336 | amide binding |
| GO:0047834 | 2 | 0.0497 | D-threo-aldose 1-dehydrogenase activity |
| GO:0016810 | 2 | 0.0541 | hydrolase activity, acting on carbon-nitrogen (but not peptide) bonds |
| GO:0016866 | 2 | 0.0571 | intramolecular transferase activity |
| GO:0008484 | 2 | 0.0728 | sulfuric ester hydrolase activity |
| GO:0008757 | 2 | 0.112 | S-adenosylmethionine-dependent methyltransferase activity |
| GO:0016829 | 2 | 0.129 | lyase activity |
| GO:0003735 | 2 | 0.2 | structural constituent of ribosome |
| GO:0016887 | 2 | 0.223 | ATP hydrolysis activity |
| GO:0005198 | 2 | 0.253 | structural molecule activity |
| GO:0017111 | 2 | 0.473 | ribonucleoside triphosphate phosphatase activity |
| GO:0016462 | 2 | 0.517 | pyrophosphatase activity |
| GO:0016818 | 2 | 0.53 | hydrolase activity, acting on acid anhydrides, in phosphorus-containing anhydrides |
| GO:0016817 | 2 | 0.532 | hydrolase activity, acting on acid anhydrides |
| GO:0008233 | 2 | 0.657 | peptidase activity |
| GO:0046914 | 2 | 0.788 | transition metal ion binding |
| GO:0022857 | 2 | 0.904 | transmembrane transporter activity |
| GO:0005215 | 2 | 0.913 | transporter activity |
| GO:0030570 | 1 | 5.78E-03 | pectate lyase activity |
| GO:1901683 | 1 | 5.78E-03 | arsenate ion transmembrane transporter activity |
| GO:0015446 | 1 | 5.78E-03 | ATPase-coupled arsenite transmembrane transporter activity |
| GO:0016837 | 1 | 5.78E-03 | carbon-oxygen lyase activity, acting on polysaccharides |
| GO:0043531 | 1 | 0.0115 | ADP binding |
| GO:0008864 | 1 | 0.0172 | formyltetrahydrofolate deformylase activity |
| GO:0046559 | 1 | 0.0229 | alpha-glucuronidase activity |
| GO:0004332 | 1 | 0.0229 | fructose-bisphosphate aldolase activity |
| GO:0016832 | 1 | 0.0286 | aldehyde-lyase activity |
| GO:0030145 | 1 | 0.0508 | manganese ion binding |
| GO:0030983 | 1 | 0.0563 | mismatched DNA binding |
| GO:0016742 | 1 | 0.0672 | hydroxymethyl-, formyl- and related transferase activity |

| GO Term | Count | p-value | Name |
|---|---|---|---|
| GO:0015116 | 1 | 0.0672 | sulfate transmembrane transporter activity |
| GO:0070006 | 1 | 0.0726 | metalloaminopeptidase activity |
| GO:1901682 | 1 | 0.0726 | sulfur compound transmembrane transporter activity |
| GO:0003887 | 1 | 0.0939 | DNA-directed DNA polymerase activity |
| GO:0008113 | 1 | 0.0939 | peptide-methionine (S)-S-oxide reductase activity |
| GO:0034061 | 1 | 0.0991 | DNA polymerase activity |
| GO:0004177 | 1 | 0.104 | aminopeptidase activity |
| GO:0003690 | 1 | 0.11 | double-stranded DNA binding |
| GO:0008235 | 1 | 0.15 | metalloexopeptidase activity |
| GO:0016671 | 1 | 0.165 | oxidoreductase, acting on a sulfur donors, disulfide acceptors |
| GO:0016830 | 1 | 0.165 | carbon-carbon lyase activity |
| GO:0016835 | 1 | 0.189 | carbon-oxygen lyase activity |
| GO:0051537 | 1 | 0.198 | 2 iron, 2 sulfur cluster binding |
| GO:0016209 | 1 | 0.261 | antioxidant activity |
| GO:0051287 | 1 | 0.265 | NAD binding |
| GO:0008238 | 1 | 0.286 | exopeptidase activity |
| GO:0015103 | 1 | 0.294 | inorganic anion transmembrane transporter activity |
| GO:0042626 | 1 | 0.307 | ATPase-coupled transmembrane transporter activity |
| GO:0015399 | 1 | 0.334 | primary active transmembrane transporter activity |
| GO:0008080 | 1 | 0.342 | N-acetyltransferase activity |
| GO:0070279 | 1 | 0.35 | vitamin B6 binding |
| GO:0030170 | 1 | 0.35 | pyridoxal phosphate binding |
| GO:0016410 | 1 | 0.35 | N-acyltransferase activity |
| GO:0016779 | 1 | 0.376 | nucleotidyltransferase activity |
| GO:0016407 | 1 | 0.379 | acetyltransferase activity |
| GO:0004252 | 1 | 0.387 | serine-type endopeptidase activity |
| GO:0051536 | 1 | 0.397 | iron-sulfur cluster binding |
| GO:0051540 | 1 | 0.397 | metal cluster binding |
| GO:0008237 | 1 | 0.401 | metallopeptidase activity |
| GO:0003700 | 1 | 0.458 | DNA-binding transcription factor activity |
| GO:0022804 | 1 | 0.488 | active transmembrane transporter activity |
| GO:0140110 | 1 | 0.497 | transcription regulator activity |
| GO:0017171 | 1 | 0.532 | serine hydrolase activity |
| GO:0008236 | 1 | 0.532 | serine-type peptidase activity |
| GO:0016859 | 1 | 0.537 | cis-trans isomerase activity |
| GO:0003755 | 1 | 0.537 | peptidyl-prolyl cis-trans isomerase activity |
| GO:0004553 | 1 | 0.54 | hydrolase activity, hydrolyzing O-glycosyl compounds |
| GO:0140097 | 1 | 0.563 | catalytic activity, acting on DNA |
| GO:0031418 | 1 | 0.598 | L-ascorbic acid binding |

| GO Term | Count | p-value | Name |
|---|---|---|---|
| GO:0043177 | 1 | 0.603 | organic acid binding |
| GO:0031406 | 1 | 0.603 | carboxylic acid binding |
| GO:0048029 | 1 | 0.607 | monosaccharide binding |
| GO:0016798 | 1 | 0.61 | hydrolase activity, acting on glycosyl bonds |
| GO:0004175 | 1 | 0.651 | endopeptidase activity |
| GO:0030246 | 1 | 0.655 | carbohydrate binding |
| GO:0016705 | 1 | 0.694 | oxidoreductase, acting on paired donors, with incorporation or reduction of molecular oxygen |
| GO:0016747 | 1 | 0.695 | acyltransferase, transferring groups other than amino-acyl |
| GO:0016757 | 1 | 0.716 | glycosyltransferase activity |
| GO:0005506 | 1 | 0.716 | iron ion binding |
| GO:0015318 | 1 | 0.723 | inorganic molecular entity transmembrane transporter activity |
| GO:0140657 | 1 | 0.863 | ATP-dependent activity |
| GO:0140640 | 1 | 0.918 | catalytic activity, acting on a nucleic acid |
| GO:0016772 | 1 | 0.966 | transferase activity, transferring phosphorus-containing groups |

**Supplemental Table 9 Biological process-related GO term enrichment results for LGTs specific to *Ac. anophagefferens* (see associated REVIGO plot in Extended Data Figure 10).** Only terms that were found to be significantly enriched (p-value <0.05) or that occurred >1 time are included.

| GO Term | Count | p-value | Name |
|---|---|---|---|
| GO:0008150 | 89 | 1 | biological process |
| GO:0008152 | 80 | 1.46E-07 | metabolic process |
| GO:0071704 | 74 | 4.21E-06 | organic substance metabolic process |
| GO:0044238 | 66 | 3.71E-04 | primary metabolic process |
| GO:1901564 | 38 | 0.0797 | organonitrogen compound metabolic process |
| GO:0043170 | 36 | 0.532 | macromolecule metabolic process |
| GO:0009987 | 33 | 1 | cellular process |
| GO:0019538 | 29 | 0.111 | protein metabolic process |
| GO:0005975 | 24 | 2.26E-10 | carbohydrate metabolic process |
| GO:0044237 | 22 | 0.8 | cellular metabolic process |
| GO:0043412 | 16 | 0.63 | macromolecule modification |
| GO:0009058 | 15 | 0.281 | biosynthetic process |
| GO:0036211 | 14 | 0.634 | protein modification process |
| GO:0006508 | 13 | 0.0107 | proteolysis |
| GO:1901360 | 11 | 0.98 | organic cyclic compound metabolic process |
| GO:1901576 | 10 | 0.732 | organic substance biosynthetic process |
| GO:0006793 | 9 | 0.88 | phosphorus metabolic process |
| GO:0006139 | 9 | 0.985 | nucleobase-containing compound metabolic process |
| GO:0006629 | 8 | 0.181 | lipid metabolic process |
| GO:0090304 | 8 | 0.933 | nucleic acid metabolic process |
| GO:0006796 | 8 | 0.934 | phosphate-containing compound metabolic process |
| GO:0044281 | 7 | 0.697 | small molecule metabolic process |
| GO:0043603 | 6 | 0.0149 | amide metabolic process |
| GO:0006790 | 6 | 0.0286 | sulfur compound metabolic process |
| GO:0018193 | 6 | 0.042 | peptidyl-amino acid modification |
| GO:0044255 | 6 | 0.0795 | cellular lipid metabolic process |
| GO:0006950 | 6 | 0.16 | response to stress |
| GO:0050896 | 6 | 0.223 | response to stimulus |
| GO:0044249 | 6 | 0.84 | cellular biosynthetic process |
| GO:0055085 | 6 | 0.978 | transmembrane transport |
| GO:0006810 | 6 | 1 | transport |
| GO:0051234 | 6 | 1 | establishment of localization |
| GO:0051179 | 6 | 1 | localization |
| GO:0006575 | 5 | 0.0254 | cellular modified amino acid metabolic process |
| GO:0006259 | 5 | 0.411 | DNA metabolic process |

| GO Term | Count | p-value | Name |
|---|---|---|---|
| GO:0006468 | 5 | 0.762 | protein phosphorylation |
| GO:1901566 | 5 | 0.774 | organonitrogen compound biosynthetic process |
| GO:0016310 | 5 | 0.794 | phosphorylation |
| GO:0006749 | 4 | 0.0507 | glutathione metabolic process |
| GO:0008610 | 4 | 0.252 | lipid biosynthetic process |
| GO:0006281 | 4 | 0.277 | DNA repair |
| GO:0006974 | 4 | 0.31 | DNA damage response |
| GO:0033554 | 4 | 0.369 | cellular response to stress |
| GO:0051716 | 4 | 0.369 | cellular response to stimulus |
| GO:0043436 | 4 | 0.602 | oxoacid metabolic process |
| GO:0009056 | 4 | 0.611 | catabolic process |
| GO:0006082 | 4 | 0.614 | organic acid metabolic process |
| GO:0072330 | 3 | 0.0505 | monocarboxylic acid biosynthetic process |
| GO:1901575 | 3 | 0.776 | organic substance catabolic process |
| GO:0065007 | 3 | 0.995 | biological regulation |
| GO:0016052 | 3 | 0.0743 | carbohydrate catabolic process |
| GO:0032787 | 3 | 0.286 | monocarboxylic acid metabolic process |
| GO:0016053 | 3 | 0.388 | organic acid biosynthetic process |
| GO:0046394 | 3 | 0.388 | carboxylic acid biosynthetic process |
| GO:0044283 | 3 | 0.569 | small molecule biosynthetic process |
| GO:0019752 | 3 | 0.788 | carboxylic acid metabolic process |
| GO:0019637 | 3 | 0.803 | organophosphate metabolic process |
| GO:0016070 | 3 | 0.99 | RNA metabolic process |
| GO:0018212 | 2 | 7.59E-04 | peptidyl-tyrosine modification |
| GO:0051923 | 2 | 7.59E-04 | sulfation |
| GO:0006478 | 2 | 7.59E-04 | peptidyl-tyrosine sulfation |
| GO:0006477 | 2 | 7.59E-04 | protein sulfation |
| GO:0007194 | 2 | 1.50E-03 | negative regulation of adenylate cyclase activity |
| GO:0051339 | 2 | 1.50E-03 | regulation of lyase activity |
| GO:0051350 | 2 | 1.50E-03 | negative regulation of lyase activity |
| GO:0031279 | 2 | 1.50E-03 | regulation of cyclase activity |
| GO:0031280 | 2 | 1.50E-03 | negative regulation of cyclase activity |
| GO:0045761 | 2 | 1.50E-03 | regulation of adenylate cyclase activity |
| GO:0044092 | 2 | 3.68E-03 | negative regulation of molecular function |
| GO:0043086 | 2 | 3.68E-03 | negative regulation of catalytic activity |
| GO:0044347 | 2 | 8.55E-03 | cell wall polysaccharide catabolic process |
| GO:2000895 | 2 | 8.55E-03 | hemicellulose catabolic process |
| GO:0045493 | 2 | 8.55E-03 | xylan catabolic process |
| GO:0010410 | 2 | 0.0128 | hemicellulose metabolic process |

| GO Term | Count | p-value | Name |
|---|---|---|---|
| GO:0045491 | 2 | 0.0128 | xylan metabolic process |
| GO:0010383 | 2 | 0.0128 | cell wall polysaccharide metabolic process |
| GO:0016998 | 2 | 0.0128 | cell wall macromolecule catabolic process |
| GO:0044036 | 2 | 0.0178 | cell wall macromolecule metabolic process |
| GO:0000272 | 2 | 0.0265 | polysaccharide catabolic process |
| GO:0065009 | 2 | 0.0692 | regulation of molecular function |
| GO:0050790 | 2 | 0.0692 | regulation of catalytic activity |
| GO:0006767 | 2 | 0.0881 | water-soluble vitamin metabolic process |
| GO:0006766 | 2 | 0.093 | vitamin metabolic process |
| GO:0006979 | 2 | 0.108 | response to oxidative stress |
| GO:0005976 | 2 | 0.108 | polysaccharide metabolic process |
| GO:0006310 | 2 | 0.124 | DNA recombination |
| GO:0033013 | 2 | 0.135 | tetrapyrrole metabolic process |
| GO:0006633 | 2 | 0.152 | fatty acid biosynthetic process |
| GO:0006631 | 2 | 0.186 | fatty acid metabolic process |
| GO:0046488 | 2 | 0.192 | phosphatidylinositol metabolic process |
| GO:0006650 | 2 | 0.233 | glycerophospholipid metabolic process |
| GO:0046486 | 2 | 0.251 | glycerolipid metabolic process |
| GO:0043414 | 2 | 0.335 | macromolecule methylation |
| GO:0006644 | 2 | 0.352 | phospholipid metabolic process |
| GO:0006486 | 2 | 0.399 | protein glycosylation |
| GO:0043413 | 2 | 0.399 | macromolecule glycosylation |
| GO:0070085 | 2 | 0.41 | glycosylation |
| GO:0032259 | 2 | 0.522 | methylation |
| GO:0009057 | 2 | 0.606 | macromolecule catabolic process |
| GO:0009451 | 2 | 0.606 | RNA modification |
| GO:0090407 | 2 | 0.611 | organophosphate biosynthetic process |
| GO:0006457 | 2 | 0.619 | protein folding |
| GO:0141188 | 2 | 0.77 | nucleic acid catabolic process |
| GO:0006520 | 2 | 0.842 | amino acid metabolic process |
| GO:0034660 | 2 | 0.848 | ncRNA metabolic process |
| GO:1901362 | 2 | 0.93 | organic cyclic compound biosynthetic process |
| GO:0006396 | 2 | 0.951 | RNA processing |
| GO:0006811 | 2 | 0.984 | monoatomic ion transport |
| GO:0019634 | 1 | 0.0161 | organic phosphonate metabolic process |
| GO:0009445 | 1 | 0.0161 | putrescine metabolic process |
| GO:0009446 | 1 | 0.0161 | putrescine biosynthetic process |
| GO:0019700 | 1 | 0.0161 | organic phosphonate catabolic process |
| GO:0006004 | 1 | 0.0161 | fucose metabolic process |

| GO Term | Count | p-value | Name |
|---|---|---|---|
| GO:1990505 | 1 | 0.0319 | mitotic DNA replication maintenance of fidelity |
| GO:1902298 | 1 | 0.0319 | cell cycle DNA replication maintenance of fidelity |
| GO:1990426 | 1 | 0.0319 | mitotic recombination-dependent replication fork processing |
| GO:0018095 | 1 | 0.0475 | protein polyglutamylation |
| GO:0009235 | 1 | 0.0475 | cobalamin metabolic process |

**Supplemental Table 10 Biological process-related GO term enrichment results for LGTs specific to *P. calceolata* (see associated REVIGO plot in Extended Data Figure 10).** Only terms that were found to be significantly enriched (p-value <0.05) or occurred >1 time are included.

| GO Term | Count | p-value | Name |
|---|---|---|---|
| GO:0008150 | 9 | 1 | biological process |
| GO:0071704 | 8 | 0.0781 | organic substance metabolic process |
| GO:0008152 | 8 | 0.133 | metabolic process |
| GO:0044238 | 6 | 0.395 | primary metabolic process |
| GO:1901564 | 5 | 0.172 | organonitrogen compound metabolic process |
| GO:0009987 | 4 | 0.896 | cellular process |
| GO:0044281 | 3 | 0.0399 | small molecule metabolic process |
| GO:0006139 | 3 | 0.206 | nucleobase-containing compound metabolic process |
| GO:1901360 | 3 | 0.262 | organic cyclic compound metabolic process |
| GO:0044237 | 3 | 0.488 | cellular metabolic process |
| GO:0043170 | 3 | 0.775 | macromolecule metabolic process |
| GO:1901135 | 2 | 0.0392 | carbohydrate derivative metabolic process |
| GO:0055086 | 2 | 0.0414 | nucleobase-containing small molecule metabolic process |
| GO:0019637 | 2 | 0.0649 | organophosphate metabolic process |
| GO:1901576 | 2 | 0.328 | organic substance biosynthetic process |
| GO:0006796 | 2 | 0.354 | phosphate-containing compound metabolic process |
| GO:0006793 | 2 | 0.357 | phosphorus metabolic process |
| GO:0009058 | 2 | 0.375 | biosynthetic process |
| GO:0019538 | 2 | 0.731 | protein metabolic process |
| GO:0008616 | 1 | 1.63E-03 | queuosine biosynthetic process |
| GO:0015878 | 1 | 1.63E-03 | biotin transport |
| GO:0046116 | 1 | 1.63E-03 | queuosine metabolic process |
| GO:0006481 | 1 | 3.25E-03 | C-terminal protein methylation |
| GO:0046164 | 1 | 3.25E-03 | alcohol catabolic process |
| GO:0018410 | 1 | 3.25E-03 | C-terminal protein amino acid modification |
| GO:0019310 | 1 | 3.25E-03 | inositol catabolic process |
| GO:0046174 | 1 | 3.25E-03 | polyol catabolic process |
| GO:1901616 | 1 | 3.25E-03 | organic hydroxy compound catabolic process |
| GO:0006020 | 1 | 4.87E-03 | inositol metabolic process |
| GO:0051180 | 1 | 4.87E-03 | vitamin transport |
| GO:0042455 | 1 | 6.49E-03 | ribonucleoside biosynthetic process |
| GO:1901659 | 1 | 6.49E-03 | glycosyl compound biosynthetic process |
| GO:0009163 | 1 | 6.49E-03 | nucleoside biosynthetic process |
| GO:0009119 | 1 | 6.49E-03 | ribonucleoside metabolic process |

| GO Term | Count | p-value | Name |
|---|---|---|---|
| GO:0034404 | 1 | 6.49E-03 | nucleobase-containing small molecule biosynthetic process |
| GO:0015718 | 1 | 6.49E-03 | monocarboxylic acid transport |
| GO:0015849 | 1 | 8.11E-03 | organic acid transport |
| GO:0046942 | 1 | 8.11E-03 | carboxylic acid transport |
| GO:0042886 | 1 | 0.0113 | amide transport |
| GO:0034035 | 1 | 0.0129 | purine ribonucleoside bisphosphate metabolic process |
| GO:0015969 | 1 | 0.0129 | guanosine tetraphosphate metabolic process |
| GO:0015711 | 1 | 0.0129 | organic anion transport |
| GO:0019751 | 1 | 0.0162 | polyol metabolic process |
| GO:0072348 | 1 | 0.0226 | sulfur compound transport |
| GO:0006284 | 1 | 0.0321 | base-excision repair |
| GO:1901657 | 1 | 0.0384 | glycosyl compound metabolic process |
| GO:0009116 | 1 | 0.0384 | nucleoside metabolic process |
| GO:0034032 | 1 | 0.04 | purine nucleoside bisphosphate metabolic process |
| GO:0033875 | 1 | 0.04 | ribonucleoside bisphosphate metabolic process |
| GO:0006066 | 1 | 0.04 | alcohol metabolic process |
| GO:0033865 | 1 | 0.04 | nucleoside bisphosphate metabolic process |
| GO:0044282 | 1 | 0.0478 | small molecule catabolic process |
| GO:0008213 | 1 | 0.0493 | protein alkylation |
| GO:0006479 | 1 | 0.0493 | protein methylation |

**Supplemental Table 11 Molecular function-related GO term enrichment results for LGTs specific to *P. calceolata* (see associated REVIGO plot in Extended Data Figure 10).** Only terms that were found to be significantly enriched (p-value <0.05) or occurred >1 time are included.

| GO Term | Count | p-value | Name |
|---|---|---|---|
| GO:0003674 | 28 | 1 | molecular function |
| GO:0003824 | 21 | 7.92E-03 | catalytic activity |
| GO:0005488 | 9 | 1 | binding |
| GO:0016491 | 8 | 5.15E-03 | oxidoreductase activity |
| GO:0016740 | 7 | 0.253 | transferase activity |
| GO:0043167 | 5 | 0.909 | ion binding |
| GO:0036094 | 5 | 0.939 | small molecule binding |
| GO:0140096 | 4 | 0.481 | catalytic activity, acting on a protein |
| GO:0097159 | 4 | 0.957 | organic cyclic compound binding |
| GO:0016614 | 3 | 5.76E-03 | oxidoreductase activity, acting on CH-OH group of donors |
| GO:0008168 | 3 | 0.0636 | methyltransferase activity |
| GO:0016741 | 3 | 0.0709 | transferase activity, transferring one-carbon groups |
| GO:0046872 | 3 | 0.75 | metal ion binding |
| GO:0043169 | 3 | 0.762 | cation binding |
| GO:1901265 | 3 | 0.791 | nucleoside phosphate binding |
| GO:0000166 | 3 | 0.791 | nucleotide binding |
| GO:0016787 | 3 | 0.822 | hydrolase activity |
| GO:0043168 | 3 | 0.843 | anion binding |
| GO:1901363 | 3 | 0.869 | heterocyclic compound binding |
| GO:0005515 | 3 | 0.998 | protein binding |
| GO:0004867 | 2 | 6.12E-05 | serine-type endopeptidase inhibitor activity |
| GO:0061135 | 2 | 9.17E-05 | endopeptidase regulator activity |
| GO:0030414 | 2 | 9.17E-05 | peptidase inhibitor activity |
| GO:0004866 | 2 | 9.17E-05 | endopeptidase inhibitor activity |
| GO:0061134 | 2 | 1.28E-04 | peptidase regulator activity |
| GO:0140678 | 2 | 2.19E-04 | molecular function inhibitor activity |
| GO:0004857 | 2 | 2.19E-04 | enzyme inhibitor activity |
| GO:0008276 | 2 | 0.0106 | protein methyltransferase activity |
| GO:0030234 | 2 | 0.0247 | enzyme regulator activity |
| GO:0008757 | 2 | 0.0322 | S-adenosylmethionine-dependent methyltransferase activity |
| GO:0098772 | 2 | 0.0338 | molecular function regulator activity |
| GO:0016616 | 2 | 0.0411 | oxidoreductase, acting on CH-OH donors, NAD acceptors |
| GO:0046914 | 2 | 0.444 | transition metal ion binding |
| GO:0016772 | 2 | 0.486 | transferase activity, transferring phosphorus-containing groups |

| GO Term | Count | p-value | Name |
|---|---|---|---|
| GO:0005524 | 2 | 0.813 | ATP binding |
| GO:0032559 | 2 | 0.818 | adenyl ribonucleotide binding |
| GO:0030554 | 2 | 0.852 | adenyl nucleotide binding |
| GO:0035639 | 2 | 0.87 | purine ribonucleoside triphosphate binding |
| GO:0032555 | 2 | 0.874 | purine ribonucleotide binding |
| GO:0032553 | 2 | 0.881 | ribonucleotide binding |
| GO:0097367 | 2 | 0.884 | carbohydrate derivative binding |
| GO:0017076 | 2 | 0.899 | purine nucleotide binding |
| GO:0004797 | 1 | 2.53E-03 | thymidine kinase activity |
| GO:0033739 | 1 | 2.53E-03 | preQ1 synthase activity |
| GO:0019136 | 1 | 2.53E-03 | deoxynucleoside kinase activity |
| GO:0015225 | 1 | 2.53E-03 | biotin transmembrane transporter activity |
| GO:0019206 | 1 | 5.05E-03 | nucleoside kinase activity |
| GO:0003880 | 1 | 5.05E-03 | protein C-terminal carboxyl O-methyltransferase activity |
| GO:0050113 | 1 | 5.05E-03 | inositol oxygenase activity |
| GO:0016920 | 1 | 7.56E-03 | pyroglutamyl-peptidase activity |
| GO:0004764 | 1 | 7.56E-03 | shikimate 3-dehydrogenase (NADP+) activity |
| GO:0090482 | 1 | 7.56E-03 | vitamin transmembrane transporter activity |
| GO:0008028 | 1 | 7.56E-03 | monocarboxylic acid transmembrane transporter activity |
| GO:0008242 | 1 | 0.0101 | omega peptidase activity |
| GO:0046857 | 1 | 0.0101 | oxidoreductase, acting on nitrogenous compounds as donors, with NAD(P) as an acceptor |
| GO:0046943 | 1 | 0.0101 | carboxylic acid transmembrane transporter activity |
| GO:0005342 | 1 | 0.0101 | organic acid transmembrane transporter activity |
| GO:0042887 | 1 | 0.0151 | amide transmembrane transporter activity |
| GO:0097506 | 1 | 0.02 | deaminated base DNA N-glycosylase activity |
| GO:0004844 | 1 | 0.02 | uracil DNA N-glycosylase activity |
| GO:0010340 | 1 | 0.025 | carboxyl-O-methyltransferase activity |
| GO:0051998 | 1 | 0.025 | protein carboxyl O-methyltransferase activity |
| GO:0008514 | 1 | 0.0274 | organic anion transmembrane transporter activity |
| GO:0016646 | 1 | 0.0299 | oxidoreductase, acting on the CH-NH donors, NAD acceptors |
| GO:0016780 | 1 | 0.0348 | phosphotransferase, for other substituted phosphate groups |
| GO:1901682 | 1 | 0.0372 | sulfur compound transmembrane transporter activity |
| GO:0019104 | 1 | 0.0372 | DNA N-glycosylase activity |
| GO:0005044 | 1 | 0.0372 | scavenger receptor activity |
| GO:0016645 | 1 | 0.0397 | oxidoreductase activity, acting on the CH-NH group of donors |
| GO:0038024 | 1 | 0.0397 | cargo receptor activity |
| GO:0019205 | 1 | 0.0494 | nucleobase-containing compound kinase activity |

**Supplemental Table 12 Biological process-related GO term enrichment results for LGTs specific to *Au. lagunensis* (see associated REVIGO plot in Extended Data Figure 10).** Only terms that were found to be significantly enriched (p-value <0.05) or occurred >1 time are included

| GO Term | Count | p-value | Name |
|---|---|---|---|
| GO:0008150 | 34 | 1 | biological process |
| GO:0008152 | 28 | 0.0284 | metabolic process |
| GO:0071704 | 25 | 0.0887 | organic substance metabolic process |
| GO:0044238 | 23 | 0.124 | primary metabolic process |
| GO:0009987 | 19 | 0.732 | cellular process |
| GO:1901564 | 12 | 0.552 | organonitrogen compound metabolic process |
| GO:0044237 | 11 | 0.353 | cellular metabolic process |
| GO:0043170 | 11 | 0.871 | macromolecule metabolic process |
| GO:0006139 | 8 | 0.25 | nucleobase-containing compound metabolic process |
| GO:1901360 | 8 | 0.367 | organic cyclic compound metabolic process |
| GO:0044281 | 7 | 0.0284 | small molecule metabolic process |
| GO:0006796 | 6 | 0.318 | phosphate-containing compound metabolic process |
| GO:0006793 | 6 | 0.322 | phosphorus metabolic process |
| GO:0019538 | 6 | 0.916 | protein metabolic process |
| GO:1901576 | 5 | 0.454 | organic substance biosynthetic process |
| GO:0009058 | 5 | 0.544 | biosynthetic process |
| GO:0006810 | 5 | 0.778 | transport |
| GO:0051234 | 5 | 0.789 | establishment of localization |
| GO:0051179 | 5 | 0.814 | localization |
| GO:0055086 | 4 | 0.0355 | nucleobase-containing small molecule metabolic process |
| GO:0006259 | 4 | 0.0739 | DNA metabolic process |
| GO:0005975 | 4 | 0.147 | carbohydrate metabolic process |
| GO:0055085 | 4 | 0.65 | transmembrane transport |
| GO:0090304 | 4 | 0.701 | nucleic acid metabolic process |
| GO:0090407 | 3 | 0.0421 | organophosphate biosynthetic process |
| GO:1901135 | 3 | 0.122 | carbohydrate derivative metabolic process |
| GO:0019637 | 3 | 0.218 | organophosphate metabolic process |
| GO:0006629 | 3 | 0.35 | lipid metabolic process |
| GO:1901566 | 3 | 0.443 | organonitrogen compound biosynthetic process |
| GO:0006508 | 3 | 0.448 | proteolysis |
| GO:0036211 | 3 | 0.938 | protein modification process |
| GO:0043412 | 3 | 0.968 | macromolecule modification |
| GO:0009084 | 2 | 7.15E-03 | glutamine family amino acid biosynthetic process |
| GO:1901657 | 2 | 9.29E-03 | glycosyl compound metabolic process |

| GO Term | Count | p-value | Name |
|---|---|---|---|
| GO:0009116 | 2 | 9.29E-03 | nucleoside metabolic process |
| GO:0009064 | 2 | 0.0125 | glutamine family amino acid metabolic process |
| GO:0015074 | 2 | 0.0143 | DNA integration |
| GO:0170038 | 2 | 0.0583 | proteinogenic amino acid biosynthetic process |
| GO:0170034 | 2 | 0.0583 | L-amino acid biosynthetic process |
| GO:1901607 | 2 | 0.0649 | alpha-amino acid biosynthetic process |
| GO:0008652 | 2 | 0.0665 | amino acid biosynthetic process |
| GO:0009165 | 2 | 0.0717 | nucleotide biosynthetic process |
| GO:1901293 | 2 | 0.0717 | nucleoside phosphate biosynthetic process |
| GO:0170039 | 2 | 0.0935 | proteinogenic amino acid metabolic process |
| GO:0170033 | 2 | 0.0935 | L-amino acid metabolic process |
| GO:1901605 | 2 | 0.133 | alpha-amino acid metabolic process |
| GO:0016053 | 2 | 0.21 | organic acid biosynthetic process |
| GO:0046394 | 2 | 0.21 | carboxylic acid biosynthetic process |
| GO:0009117 | 2 | 0.26 | nucleotide metabolic process |
| GO:0034654 | 2 | 0.267 | nucleobase-containing compound biosynthetic process |
| GO:0006281 | 2 | 0.272 | DNA repair |
| GO:0006753 | 2 | 0.272 | nucleoside phosphate metabolic process |
| GO:0006974 | 2 | 0.292 | DNA damage response |
| GO:0044283 | 2 | 0.311 | small molecule biosynthetic process |
| GO:0051716 | 2 | 0.329 | cellular response to stimulus |
| GO:0033554 | 2 | 0.329 | cellular response to stress |
| GO:0006520 | 2 | 0.354 | amino acid metabolic process |
| GO:0006950 | 2 | 0.412 | response to stress |
| GO:1901575 | 2 | 0.461 | organic substance catabolic process |
| GO:0050896 | 2 | 0.467 | response to stimulus |
| GO:0019752 | 2 | 0.471 | carboxylic acid metabolic process |
| GO:0043436 | 2 | 0.475 | oxoacid metabolic process |
| GO:0009056 | 2 | 0.481 | catabolic process |
| GO:0006082 | 2 | 0.483 | organic acid metabolic process |
| GO:1901362 | 2 | 0.485 | organic cyclic compound biosynthetic process |
| GO:0006468 | 2 | 0.703 | protein phosphorylation |
| GO:0016310 | 2 | 0.725 | phosphorylation |
| GO:0050794 | 2 | 0.83 | regulation of cellular process |
| GO:0044249 | 2 | 0.834 | cellular biosynthetic process |
| GO:0050789 | 2 | 0.846 | regulation of biological process |
| GO:0065007 | 2 | 0.865 | biological regulation |
| GO:0005997 | 1 | 6.14E-03 | xylulose metabolic process |
| GO:0010133 | 1 | 6.14E-03 | proline catabolic process to glutamate |

| GO Term | Count | p-value | Name |
|---|---|---|---|
| GO:0006562 | 1 | 0.0183 | proline catabolic process |
| GO:0009065 | 1 | 0.0244 | glutamine family amino acid catabolic process |
| GO:0007156 | 1 | 0.0303 | homophilic cell adhesion via plasma membrane adhesion molecules |
| GO:0006561 | 1 | 0.0303 | proline biosynthetic process |
| GO:0098742 | 1 | 0.0303 | cell-cell adhesion via plasma-membrane adhesion molecules |
| GO:0098609 | 1 | 0.0363 | cell-cell adhesion |
| GO:0006542 | 1 | 0.0363 | glutamine biosynthetic process |
| GO:0006536 | 1 | 0.0422 | glutamate metabolic process |
| GO:0006560 | 1 | 0.0422 | proline metabolic process |
| GO:0006307 | 1 | 0.0481 | DNA alkylation repair |
