## Supplementary Figure 1 for "Pangenome biology and evolution in harmful algal-bloom-forming pelagophyte algae"

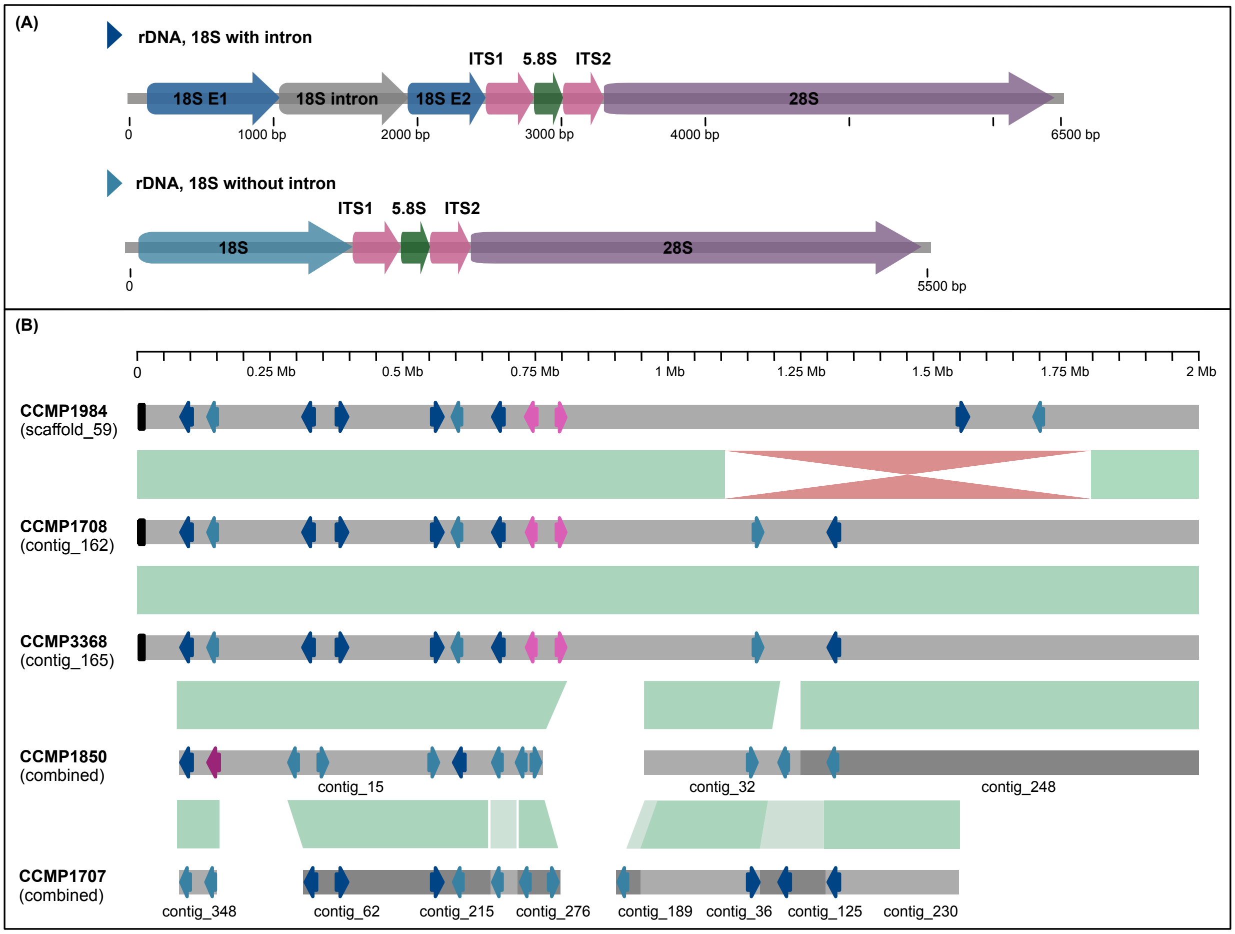


**Supplemental Figure 1 Ribosomal DNA structure and arrangement within *Aureococcus anophagefferens* strains**. (A) A schematic showing the two different rDNA versions observed within the *Aureococcus anophagefferens* genome that differ in the presence of a predicted ~900 bp intron in the 18S rDNA. (B) The genomic location of rDNA copies in the genome assembly of each strain. Contigs between the strains are oriented and linked to indicate synteny (green = linear match, red = inversion). Black boxes at the end of contigs represent telomere regions. Colored arrows indicate the direction of the rDNA copy, while dark or light blue indicates whether the 18S gene contains the predicted intron or not, as indicated in (A). Dark or light pink colours indicate similar 18S gene structures but more divergent 18S rDNA copies (~93% identity). For simplicity, only the first 2 Mbp of the rDNA containing contig in CCMP1984, CCMP1708 and CCMP3368 are shown, and only regions localized to rDNA copies on multiple contigs in CCMP1707 and CCMP1850 are shown.
